# Combinatorial docking and molecular generation to navigate over 100-billion molecules for prospective ligand discovery

**DOI:** 10.64898/2026.06.07.730716

**Authors:** Jianxiang Zhang, Chao Yang, Yang Zhang, Xiao Chen, Brandon Lam, Claire Bryant, Shabareesh Pidathala, Yangzhi Wang, Yurii S. Moroz, Dmytro S. Radchenko, Assaf Alon, Chia-Hsueh Lee, Zhe Zhang, Jiankun Lyu

**Affiliations:** The Evnin Family Laboratory of Computational Molecular Discovery, The Rockefeller University, New York, NY, USA; The David Rockefeller Graduate Program in Bioscience, The Rockefeller University, New York NY, USA; State Key Laboratory of Natural and Biomimetic Drugs, Department of Molecular and Cellular Pharmacology, School of Pharmaceutical Sciences, Peking University, Beijing, China; Peking University-Yunnan Baiyao International Medical Research Center, Beijing, China; School of Life Sciences, Center for Life Sciences, Academy for Advanced Interdisciplinary Studies, Peking University, Beijing, China; Structural Biology, St. Jude Children’s Research Hospital, Memphis, TN, USA; School of Medicine, Yale University, New Haven, CT, USA; Tri-Institutional PhD Program in Computational Biology and Medicine, New York, NY, USA; Enamine Ltd (www.enamine.net), Winston Churchill Street 78, Kyїv 02094, Ukraine; Taras Shevchenko National University of Kyiv, Volodymyrska Street 60, 01601, Kyїv, Ukraine; Chemspace LLC (www.chem-space.com), Winston Churchill Street 85, Kyїv 02094, Ukraine

## Abstract

Commercially available make-on-demand libraries now exceed 100 billion compounds, requiring over 50 years to screen on 2,000 CPU cores using conventional docking. We present two complementary approaches to address this challenge. CombiDOCK, a combinatorial docking framework, enables exhaustive screening at the 100-billion scale within 40 days. MINT-Dock, a generative framework, accelerates navigation of this space by integrating CombiDOCK with Monte Carlo Tree Search. Benchmarked on 46 diverse targets, CombiDOCK matched full-molecule docking accuracy, and MINT-Dock achieved a 4,800-fold enrichment over random selection. Compared with prior billion-scale brute-force campaigns against σ_2_, VMAT2, and VAChT, prospective CombiDOCK screens of the 100-billion-molecule library yielded higher hit rates and more potent ligands, while MINT-Dock achieved comparable outcomes across single- and multi-target objectives with >20-fold computational cost reductions. Docking-predicted poses of the best VAChT-binding compounds were confirmed by cryo-EM structures. These methods provide exhaustive and generative paths for navigating the trillion-molecule frontier of drug discovery.

## Introduction

Recently, tangible virtual libraries fueled by diverse building block catalogs and robust parallel synthesis reactions (>80% success rates)^1,2^ have become major drivers in the expansion of commercially available chemical space. These fast-growing make-on-demand chemical libraries now exceed 100 billion structurally diverse and stereogenic molecules, offering an unprecedented opportunity for ligand and drug discovery. Recent prospective studies further suggest that well-predicted protein models can extend ligand discovery beyond experimentally determined structures, thereby broadening the range of targets accessible to virtual screening ^3–7^. Such large tangible libraries can only be screened computationally, and even then, require efficient and accurate methods ^8,9^. Many groups have leveraged these libraries for ligand discovery, including more than 45 prospective virtual screening campaigns,^1,4,10–43^ the majority of which rely on brute-force screening methods. When directly compared, larger libraries consistently identify more potent hits than screening smaller libraries,^1,4,25,26,35,39,40^ emphasizing the importance of utilizing larger libraries for virtual campaigns as they have become available.

Even with these advances, structure-based virtual screening (SBVS) remains fundamentally constrained by its underlying computational architecture. SBVS workflows typically rely on either on-the-fly ligand construction or pre-built conformer libraries.^44^ Both workflows have distinct computational and storage demands; regardless, a 100-billion compound screen would take over 50 years on a 2000-core cluster for either approach (**Fig. 1A**, see details in **Table S1**),^45–51^ emphasizing that these strategies cannot feasibly scale to libraries on the order of 100 billion molecules. This scaling challenge is further amplified when discovery goals go beyond single-target binding to multi-target optimization, such as achieving selectivity over antitargets or polypharmacology across related receptors, goals that are central to many drugs and chemical probes^52–61^ but require separate exhaustive screens against each target, multiplying already prohibitive computational cost.

**Figure 1.**
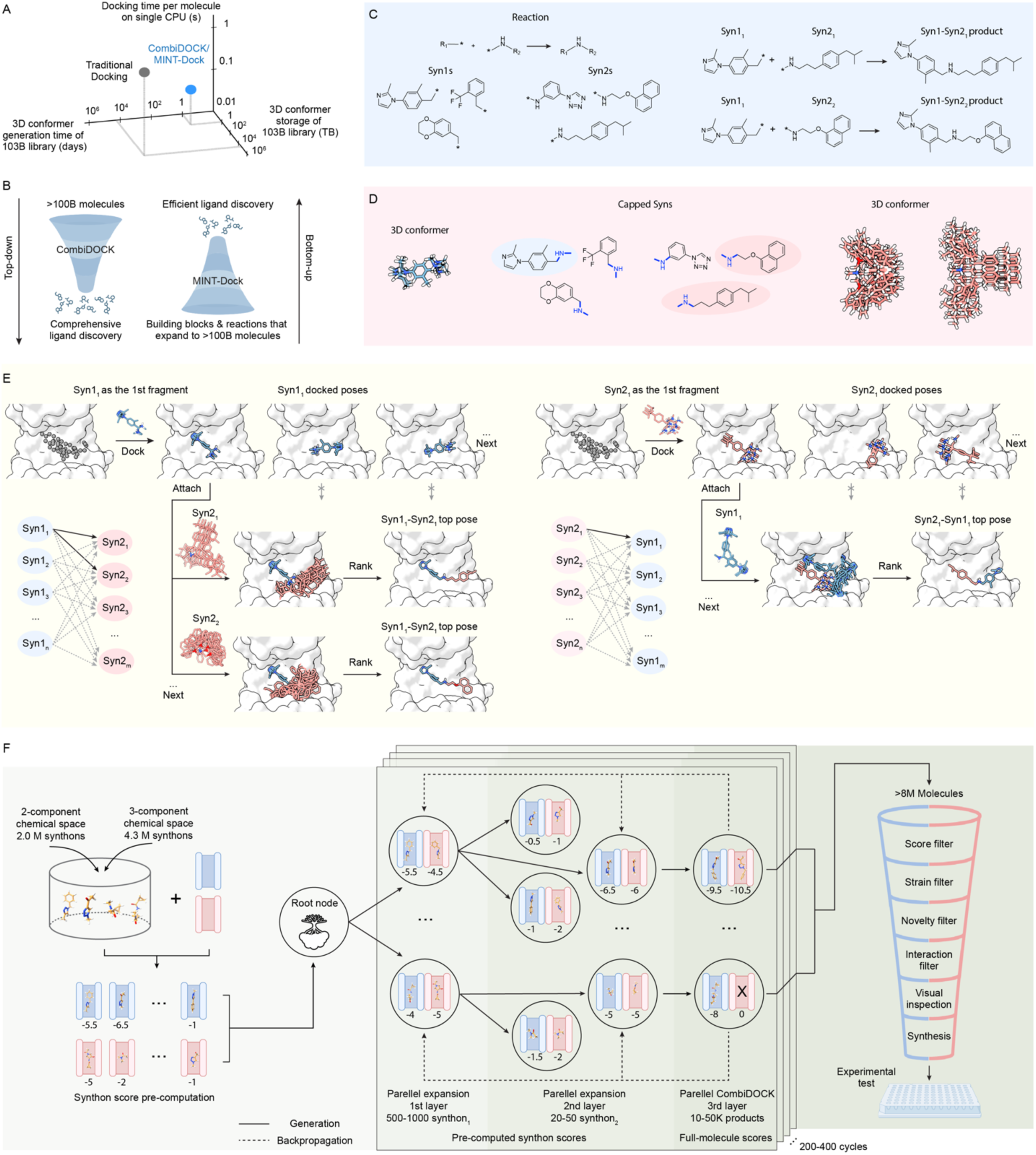
CombiDOCK and MINT-Dock enable exhaustive and guided navigation of ultra-large combinatorial chemical space. (A) Comparison of the CombiDOCK/MINT-Dock workflow with the traditional full-molecule docking workflow, represented here by UCSF DOCK, across three major computational dimensions: time required for 3D conformer generation, storage required for the library, and docking time per molecule. CombiDOCK/MINT-Dock improves efficiency across all three dimensions relative to conventional full-molecule docking. Data are available in **Table S1**. (B) Conceptual relationship between the two methods. CombiDOCK performs top-down, exhaustive screening of ultra-large chemical libraries, whereas MINT-Dock performs bottom-up, guided exploration of the same chemical space for efficient ligand discovery. (C) The REAL Space virtual library was constructed combinatorially from 304 chemical reactions and 2.3 million synthons. One representative reaction type is shown, illustrating how synthons (Syn1s and Syn2s) can be combined to generate diverse products. (D) Examples of capped synthons and the hierarchical organization of their precomputed 3D conformers. (E) Overview of the CombiDOCK workflow, showing how docking poses of the first synthon are reused across multiple product combinations to accelerate virtual screening. Nonpolar hydrogens are hidden in this panel for clarity. (F) The Overview of the MINT-Dock workflow, shown here for a multi-target optimization task. Precomputed synthon scores guide Monte Carlo Tree Search through synthon space, while full-molecule scores from the CombiDOCK engine are used for product evaluation and backpropagation, enabling efficient sampling of a small, high-value subset of ultra-large chemical space.

Despite the clear advantages of the larger libraries, their vastness poses substantial challenges for conventional docking approaches which struggle to scale beyond a few billion molecules. Several strategies have been developed to reduce this burden, including fragment-enrichment methods, such as Chemical Space Docking or V-SYNTHES,^10,12^ and machine-learning (ML) surrogate approaches.^41,43,62,63^ These approaches applied prospectively have identified sub-micromolar to low micromolar binders across multiple targets,^10,12–14,16–20,42,43^ all while screening far smaller libraries. Despite their efficiency, these approaches inevitably sacrifice accuracy: because they do not evaluate every molecule explicitly with the scoring function, relying instead on predicted or fragment-derived docking scores to filter the library, inaccuracies can propagate and many true positives are likely to be missed. In parallel, generative methods have emerged as an alternative way to explore only high-value regions of chemical space, enabling increasingly efficient discovery of novel and potent bioactives.^64–83^ However, current structure-based generative methods face a fundamental tradeoff among synthesizability,^69,70,72^ 3D pose prediction^74,77,82^ and generation throughput.^71,81^ As a result, relatively few studies have carried designs through synthesis and experimental testing at scale.

These challenges motivate us to develop two complementary computational goals for exploring over 100 billion-molecule make-on-demand chemical space (**Fig. 1B**). One is top-down, comprehensive virtual screening: to dock every molecule in a combinatorial library as accurately as possible without explicit full-molecule enumeration, thereby preserving the exhaustive logic of conventional docking while overcoming its scaling bottlenecks. The other is bottom-up, efficient navigation: to start from the underlying building blocks and reactions and use structure-guided search to identify a much smaller set of high-value candidates while retaining explicit 3D evaluation of full molecules.

Here, we introduce two methods that achieve these complementary goals. For the top-down approach, we present CombiDOCK, a combinatorial docking approach that enables efficient, exhaustive screening of ultra-large chemical space by decomposing molecules into synthons and reconstructing full molecules on-the-fly during docking. Here, a synthon is defined as a building block fragment expressed in its product form, with reactive attachment points that specify how it can combine with other synthons under well-defined reaction rules.^10^ CombiDOCK is not a “fragment-enrichment then enumerate” workflow. It eliminates explicit full-molecule enumeration throughout both library representation and docking, enabling full-molecule-equivalent docking at >100 billion scale. For the bottom-up approach, building on CombiDOCK’s combinatorial docking engine, we introduce MINT-Dock (MCTS Integrated Navigation Tool for Docking), a structure-based generative docking framework that couples Monte Carlo Tree Search (MCTS) with CombiDOCK to navigate make-on-demand chemical space by assembling reaction-compatible synthons and evaluating full molecules in 3D on the fly. While CombiDOCK exhaustively evaluates all possible combinations, MINT-Dock provides an adaptive, on-demand exploration that generates only the most promising products under explicit design objectives, including multi-target optimization for selectivity and polypharmacology, tasks that are otherwise computationally prohibitive at the ultra-large library scale.

To demonstrate the broad utility of CombiDOCK and MINT-Dock, we performed both retrospective testing against 46 diverse drug targets as well as prospective testing of up to 103-billion molecules in single and multi-target objective campaigns against the Sigma-2 (σ_2_) receptor, vesicular monoamine transporter 2 (VMAT2) and vesicular acetylcholine transporter (VAChT). These three targets provide clinically relevant and methodologically stringent test cases of the method, and to our knowledge, these are the first docking campaigns against any target spanning over 100-billion compounds. To prospectively assess the hypothesis of whether bigger chemical libraries improve outcomes of ligand discovery, we synthesized and tested hundreds of top-ranking molecules from each screen. Hit rates and ligand affinities were compared with those from smaller library screens.

### CombiDOCK efficiently addresses combinatorial virtual libraries through its modular design

Inspired by molecular fragmentation strategies,^10,12^ we designed CombiDOCK (implemented in DOCK6^45^), a new docking method that builds ligands in the protein binding site by combining synthons using well-defined chemical transformations. Next, to further accelerate the navigation of ultra-large combinatorial chemical space, we introduce MINT-Dock, a structure-based generative framework that couples Monte Carlo Tree Search with CombiDOCK. Both methods operate on the same synthon-based representation of make-on-demand libraries and use the same CombiDOCK engine for evaluating full molecules in 3D, but differ in search strategy, with CombiDOCK enabling exhaustive exploration and MINT-Dock enabling adaptive navigation.

Using synthons and reactions to represent modern combinatorial libraries provides a far more compact and scalable representation than explicit enumeration of every possible product (**Fig. 1C**). This abstraction enabled us to encode 103 billion stereochemically rich and diverse molecules from Enamine REAL Space using a relatively small set of 2.3 million synthons and 304 reaction schemes. Because each building block can participate in one or more reaction schemes, we denote the combination of a building block–reaction pair as a synthon, yielding 6.3 million synthons in total. The resulting chemical space is partitioned into a two-component space (2.0 million synthons, 276 reactions; 5.5 billion molecules) and a three-component space (4.3 million synthons, 28 reactions; 97.7 billion molecules). For three-component reactions, because they involve a dependency in which one synthon serves as a linker connecting the other two, we pre-react the two smallest dependent synthon sets by merging the linker with one of the other synthons, generating a pseudo two-component representation compatible with both CombiDOCK and MINT-Dock, which currently combines only two synthons to assemble a full molecule. To enable structure-based modeling, synthons are capped according to the Markush structure of the reaction they participate in,^10^ preserving the local chemical environment of the final product while providing an alignment handle for assembling two 3D synthons into a full molecule (**Fig. 1C**). These capped synthons are then stereochemically expanded into 3D conformers, resulting in a diverse library of 103 billion molecules. This synthon-based representation, which more closely reflects the chemical environment of the final products than a building-block representation, plays an important role in both methods (**Fig. 1D**).

In comparison to the pre-sampling methods such as DOCK3^50^ and its DOCK6 re-implementation,^84^ CombiDOCK utilizes a computationally efficient on-the-fly approach capable of exhaustively evaluating the 50-fold larger combinatorial virtual libraries. (**Fig. 1E**). To avoid the redundancy of the pre-sampling methods, CombiDOCK decouples sampling: rather than pre-sampling the conformers of the entire compound at once, synthon 1 (Syn1) and synthon 2 (Syn2), which together constitute the full molecule, are sampled independently to generate conformational ensembles (**Fig. S1**). From here, the algorithm performs similar hierarchical growth to DOCK3, however, CombiDOCK decomposes the process into synthon-level growth steps: capped Syn1 is docked first, followed by capped Syn2, which is assembled under the corresponding reaction geometry. Because this procedure simply breaks down the same hierarchical growth process into synthon-defined stages, CombiDOCK is designed to produce results equivalent to full-molecule docking, while avoiding redundant sampling of identical substructures across the library. The power of CombiDOCK lies in its ability to scale with the growing size of combinatorial libraries. Instead of enumerating and docking every possible full molecule, which becomes intractable at >10^10^ scale, CombiDOCK docks each synthon fragment once to generate its binding poses within the receptor. These cached synthon poses are then reused to assemble full molecules on-the-fly, ensuring that fragment-level and product-level docking remain consistent. As illustrated in the **Fig. 1E**, Syn1 is first docked into the binding site to produce an ensemble of poses. Partner synthons (Syn2s) are then introduced and combined with the cached Syn1 poses to yield full Syn1-Syn2 products, which are subsequently ranked by docking score. The process is repeated with Syn2 as the first fragment, ensuring both orientations are explored, after which all possible Syn1-Syn2 combinations can be reconstructed without redundant calculations.

### MINT-Dock: Structure-based generative docking with Monte Carlo Tree Search

While CombiDOCK exhaustively evaluates the full combinatorial space, MINT-Dock uses a Monte Carlo Tree Search (MCTS) architecture to traverse this space while remaining strictly within synthesizable, reaction-defined combinations. Synthon docking scores are precomputed once using DOCK6 HDB docking on reaction-capped synthons^85^ (see **Methods**), and these scores guide early-stage exploration without requiring product-level docking at every step. The search is organized as a three-layer tree. In the first two layers, each node corresponds to a synthon, and each path specifies a unique combination of reaction-compatible synthons that can be merged into a product node in the third (final) layer. As illustrated in **Fig. 1F**, the tree expands in parallel each cycle, typically branching to 500 to 1,000 synthon_1_ choices at the first layer and 20 to 50 synthon_2_ choices at the second layer, prioritized by synthon scores. When a branch reaches the product layer, the selected synthons are reacted *in silico* to form full molecules, and 10,000 to 50,000 products are docked in parallel using CombiDOCK to obtain full-molecule scores (**Fig. 1F**).

For each synthon combination, only the top-ranked stereoisomer is retained to promote diversity in the generated pool. The resulting product-level score is used as the MCTS reward and is backpropagated to the corresponding synthon nodes, updating their expected values and biasing subsequent rollouts toward higher-reward branches. Repeating this process over 200 to 400 cycles separately on two-component and three-component space typically yields over 8 million docked, synthesizable molecules per campaign, which are then passed through downstream triage (score, ligand strain, novelty to knowns, and interaction filters), followed by visual inspection and compound synthesis (**Fig. 1F**). For multi-target tasks such as selectivity or polypharmacology, MINT-Dock defines a design function that maps per-target docking scores to a single scalar reward used for node selection and backpropagation. For selectivity, the design function increases with the score of the intended on-target and decreases with the score of the off-target, whereas for polypharmacology, it increases with the scores of both targets. During search, the design function is evaluated using the precomputed synthon docking scores for each target, and during backpropagation it is evaluated using the full-molecule docking scores for each target.

Next, using σ_2_ and VMAT2 as test cases, we evaluated two features that improve MINT-Dock search performance. We first compared the accumulation of well-scored molecules with and without backpropagation across rollouts and found that backpropagation consistently increased the yield of well-scoring molecules across multiple score cutoffs, and the advantage became more pronounced as the number of cycles increased (**Fig. S2** and **S3**, see details in **Methods**). We then compared MINT-Dock to an ML-guided MCTS baseline (“ML-MCTS”, adapted from Synthemol^67^) to assess whether coupling MCTS to physics-based docking improves generation relative to an ML surrogate. In both targets, MINT-Dock produced substantially more well-scored molecules above any given score threshold, yielding higher enrichment across thresholds (**Fig. S4**, see details in **Methods**).

### Benchmarking CombiDOCK/MINT-Dock against full-molecule docking on DUDE-Z benchmark set

To assess the generalizability of CombiDOCK’s performance, we evaluated it against 43 diverse protein targets from the widely used DUDE-Z benchmark set (dudez.docking.org),^86^ along with 3 prospectively tested targets, for a total of 46. These targets span diverse protein classes (enzymes, proteases, GPCRs, kinases, nuclear receptors, among others) and were curated on the basis of the availability of complete resolved structures and known ligands, as well as a broad range of preferred ligand charge states. This benchmark set ensures that performance is measured across a broad range of pharmacologically relevant targets with varied binding site characteristics (see **Table S2**) and enables direct comparison across different docking engines^87–90^ (see **Table S3**). Each target includes experimentally tested actives and computationally generated decoys that are property-matched to the actives but topologically dissimilar. To represent this benchmarking set in synthon space for direct comparison to conventional full-molecule docking, each full molecule was systematically decomposed into its constituent synthons and capped (**Fig. 2A**). The resulting capped synthon libraries were then organized by reaction type, serving as reusable inputs for CombiDOCK docking. Across the 46 targets, docking scores between full-molecule docking and CombiDOCK were highly correlated, with a mean Pearson correlation coefficient (*R*) of 0.82 (**Fig. 2B**). Enrichment of known actives over decoys was nearly identical, with a mean Δlog-adjusted area under the curve (ΔlogAUC) of 0.97 between full-molecule docking and CombiDOCK (**Fig. 2C**). For the three prospective targets: σ_2_ receptor, VMAT2, and VAChT, docking scores were again strongly correlated (*R* = 0.91 for σ_2_, 0.82 for VMAT2, and 0.91 for VAChT), with nearly identical enrichment values (**Fig. 2B-C**). These findings demonstrate that CombiDOCK achieves accuracy comparable to full-molecule docking while offering a fundamentally more scalable framework.

**Figure 2.**
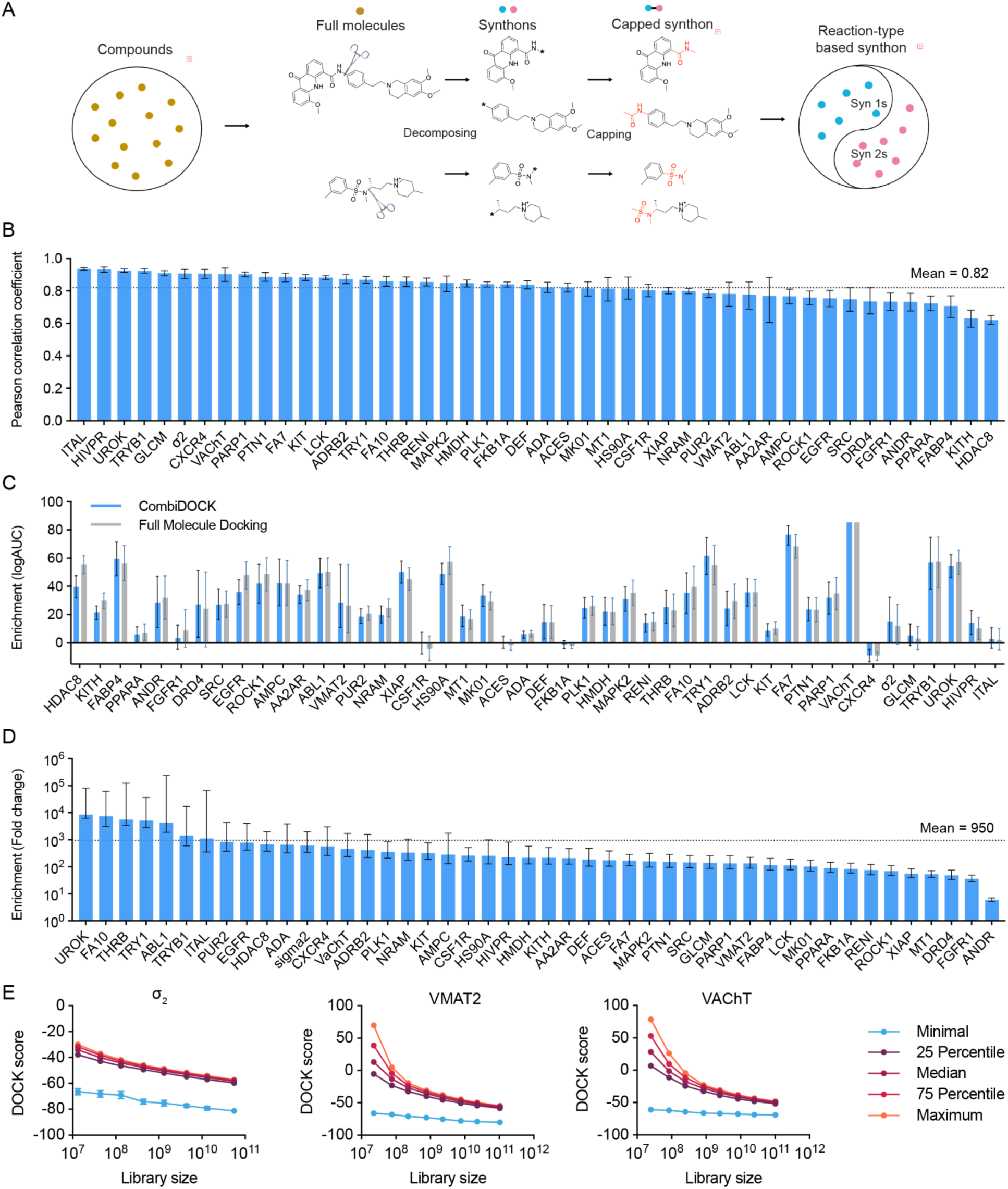
CombiDOCK approach to navigate Enamine REAL Space. (A) Compounds from pre-assembled libraries were decomposed into pairwise synthons, enabling combinatorial reassembly during docking. (B-C). Across 46 protein targets — including 43 from the DUDE-Z benchmark as well as the σ_2_ receptor (TMEM97), VMAT2 (SLC18A2), and VAChT (SLC18A3) — CombiDOCK achieved performance comparable to full-molecule docking, as assessed by B, Pearson correlation of docking scores and C, logAUC-based early enrichment. Error bars represent 90% confidence intervals, estimated by bootstrapping (1,000 iterations with replacement). (D) Summary plot of the adjustment enrichment ratio between molecules generated by MINT-Dock and molecules randomly sampled at a designated score threshold for all 46 targets, where the random library was docked by CombiDOCK. Error bars represent 90% confidence intervals, estimated by bootstrapping (1,000 iterations with replacement). (E) Docking score improves with increasing library size, illustrated for σ_2_ (56 billion), VMAT2 (103 billion), and VAChT (103 billion).

Next, to quantify the computational efficiency of MINT-Dock, we benchmarked it on the same DUDE-Z set for its ability to enrich well-scored molecules relative to docking a random subset from the same library (see details in **Methods**). For each of the 46 targets, we computed an adjusted enrichment that compares the number of molecules exceeding a given score threshold between MINT-Dock and the random subset while accounting for differences in sample size and upweighting more favorable scores (see details in **Methods**). **Fig. 2D** summarizes the adjusted enrichment at the designated score threshold across all 46 targets, spanning orders of magnitude and resulting in a mean improvement of 950-fold over random selection when the random subset was docked with CombiDOCK. We report this comparison because CombiDOCK is optimized for combinatorial libraries and is the same product-level docking engine used inside MINT-Dock, making it a particularly stringent and method-matched comparison. In parallel, we also benchmarked against one of the fastest conventional full-molecule docking methods, DOCK6 HDB, to provide a more general comparison for library-agnostic reference. Under this baseline, the mean enrichment increased by 4,810-fold (**Fig. S5A**). A more fine-grained enrichment analysis on three representative targets showed that MINT-Dock consistently produced more molecules above any given score cutoff than random sampling, with the strongest enrichment at the most stringent thresholds, indicating that the search preferentially concentrates docking on productive regions of chemical space (**Fig. S5B**).

### Docking scores continue to improve with library size at the >100-billion-molecule scale

A central question for make-on-demand combinatorial libraries is whether expanding their size continues to yield more favorable molecules or whether performance eventually saturates beyond 5 billion molecules. This question has been difficult to address because the computational cost of exhaustive screening at this scale is prohibitive for conventional methods. CombiDOCK overcomes this barrier, enabling brute-force screening of entire libraries and thus allowing this scaling question to be tested directly. As a proxy, we examined how docking scores improve with increasing library size.^91^ Although docking scores are imperfect predictors of binding affinity, they have been shown to correlate with experimental hit rates,^25^ and they remain the primary criterion for compound selection in virtual screening.

Before scaling to 100+-billion-molecule libraries, we first optimized the two sampling parameters that govern CombiDOCK’s speed-accuracy tradeoff (**Fig. S6**, see details in **Methods**). With these parameters fixed, we docked progressively larger subsets of Enamine REAL Space against three prospective targets: σ_2_ receptor, VMAT2, and VAChT to test scaling behavior. From the largest enumerated sets (56 billion for σ_2_, 103 billion for VMAT2, and 103 billion for VAChT), we sampled 20 random subsets spanning approximately 10^7^ to 10^11^ molecules in half-logarithmic increments. For each subset, we analyzed the top-ranked molecules: 300,000 for σ_2_ and 1 million for VMAT2 and VAChT, calculating their minimum, maximum, median, and quartile docking scores. As shown in **Fig. 2E**, the docking scores of top-ranking molecules improved monotonically and approximately log-linearly with increasing library size across all three targets. This improvement was consistent across quartiles, indicating that the entire distribution of high-ranking compounds steadily shifts toward better fits as the library grows. Importantly, no signs of score saturation were observed, suggesting that even at scales beyond 10^11^ molecules, further expansion continues to yield better-fitting candidates. The computational cost of these screens was substantial but tractable with modern resources: using a cluster of 2,000 CPU cores, docking required 40 cluster days for σ_2_ (56B molecules), 33 cluster days for VMAT2 (103B molecules), and 26 cluster days for VAChT (103B molecules). These results highlight that both the quality of top-ranking molecules and the feasibility of screening at trillion scale improve hand-in-hand, supporting the case for continued library expansion.

### Prospective testing on combinatorial docking and molecular generation

To test whether these computational gains translate into ligand discovery with improved outcomes, we conducted prospective screens using conventional docking method DOCK3, CombiDOCK and MINT-Dock across both well-studied and challenging targets for structure-based drug design. Here, we sought to evaluate whether guided search of the 100 billion molecule library continues to yield novel and more potent chemotypes with substantially less computational cost relative to conventional docking methods. We address these questions target by target, beginning with the σ_2_ receptor.

### Prospective combinatorial docking and molecular generation against the σ_2_ receptor

To address this question in a setting with a well-established large-scale docking baseline, we chose σ_2_ as benchmark target. Because σ_2_ has a previous prospective campaign that screened 490 million molecules,^27^ this enables an apples-to-apples comparison across both library sizes and search strategies. In this earlier effort, 138 high-scoring compounds from top 300,000 were tested in a competition binding assay with [^3^H]-DTG and the known σ_2_ ligand PB28 was included as a positive control. The hit rate was 51% (70/138 displaced >50% [^3^H]-DTG at 1 µM) (**Fig. 3A-B**). The top 21 potent ligands exhibiting Ki values of 30 nM to the low µM range (6 with Ki < 50 nM, none < 10 nM) (**Fig. 3C-D**). This campaign serves as the baseline for evaluating both CombiDOCK and MINT-Dock below.

**Figure 3.**
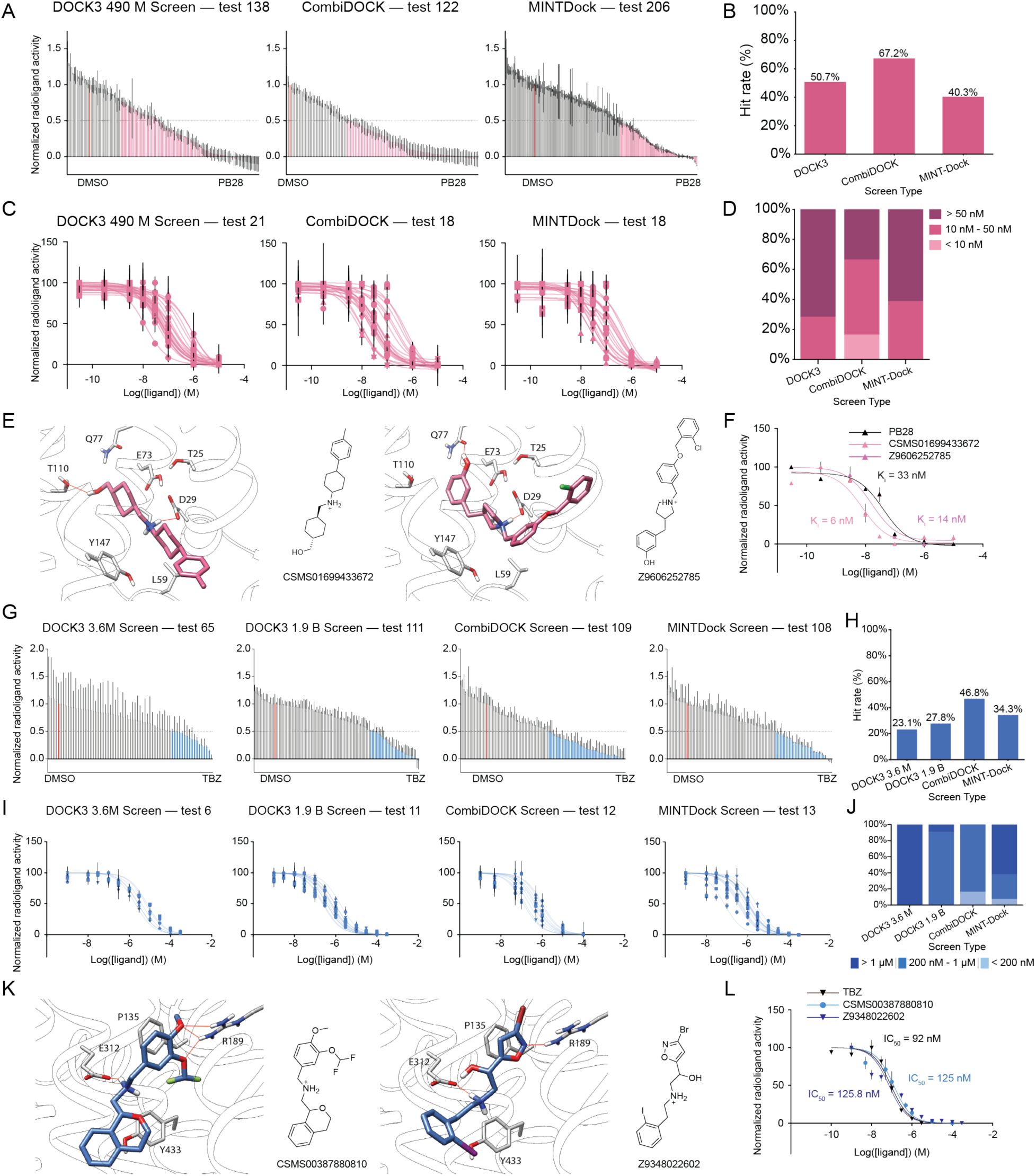
Summary of virtual screening results against σ_2_ and VMAT2. (A) In a [^3^H]-DTG binding assay, using > 50% [^3^H]-DTG displacement as a hit threshold, out of 138 tested compounds from the 490 million screen against σ_2_, 70 hits were identified (left) representing a hit rate of 51%. From the 122 tested compounds from the 56 billion screen 82 hits were identified (middle), representing a hit rate of 67%. From the 206 compounds tested from the MINT-Dock 103 billion screen, 83 hits were identified (right), representing a hit rate of 40.3%. (B) Hit rates for 490 million, CombiDOCK 56 billion, and MINT-Dock 103 billion sized library screens. (C) Dose-response curves of 21 re-tested compounds from the previous 490 million screen (left),18 tested compounds from the CombiDOCK screen (middle), and 18 tested compounds from the MINT-Dock screen. Curve of positive control PB28 (red curve) is in dose-response curves from the previous 490 million screen (left). (D) The potency distribution shifts towards higher potency compounds from the 490 million screen to the larger CombiDOCK and MINT-Dock screens. (E) Predicted pose and structure of the most potent compound from the CombiDOCK screen (left), and the MINT-Dock screen (right). (F) The dose-response curves of the most potent compound from the CombiDOCK screen and the MINT-Dock screen, with PB28 as the control. (G) In a [^3^H]-dopamine uptake assay, using > 50% [^3^H]-dopamine displacement as a hit threshold, 15 out of 65 tested compounds from the 3.6 million screen were hits representing a hit rate of 23% (top left). In the 1.9 billion screen against VMAT2, 30 out of 111 tested compounds were hits representing a hit rate of 27% (middle-left). In the 103 billion CombiDOCK screen against VMAT2, 51 out of 109 tested compounds were hits representing a hit rate of 47% (middle right). In the 103 billion MINT-Dock guided screen, 37 out of 108 tested compounds were hits representing a hit rate of 34% (right). (H) Hit rate steadily improves as screening size grows. (I) Dose-response curves of tested compounds across screening size. (J) The potency distribution shifts towards higher potency compounds as the screening size grows. (K) Predicted poses, and structures of the most potent compounds from the 103 billion screen against VMAT2 using CombiDOCK (left) and MINT-Dock (right). (L) The dose-response curves of the most potent compound from the CombiDOCK screen and the MINT-Dock screen, with TBZ as the control.

#### CombiDOCK exhaustive screen (56 billion molecules)

Using CombiDOCK, we screened 56 billion compounds from the REAL Space virtual library against the same σ_2_ receptor crystal structure in complex with cholesterol (PDB ID: 7MFI).^27^ On a 2,000-CPU-core cluster, docking required 40 cluster-days, despite the 100-fold increase in library size relative to our previous conventional docking campaign, in which 490 million make-on-demand compounds from the ZINC20 library^2^ were screened using DOCK3^27^ in three cluster-days. To match the selection depth used in the prior 490-million-compound campaign, 122 high-ranking molecules were synthesized and tested from the top 300,000 scoring molecules. For consistency, we used the same post-docking filters for selecting molecules from the previous 490 million molecule campaign^27^ (detailed in the **Methods**) and the same binding assay conditions, including PB28 as a positive control. This yielded a hit rate of 67% (82/122 displaced >50% [^3^H]-DTG). Strikingly, the top 18 actives spanned affinities ranging from 6 to 473 nM, with 12 ligands < 50 nM and three < 10 nM (**Fig. 3C-D**). The highest affinity molecule CSMS01699433672 has a K_i_ value of 6 nM, whose docked pose forms key salt bridge interaction with D29 as well as an additional hydrogen bond with T110 (**Fig. 3E**). Its dose-response curve is presented in **Fig. 3F**.

#### MINT-Dock guided search (103 billion molecules)

Starting from the synthons of the 103 billion Enamine REAL Space, MINT-Dock generated 10.3 million compounds for downstream processing in 4 days with approximately 100 CPU cores. From the top 300,000 ranked compounds from MINT-Dock screen, 206 compounds were synthesized and tested. Eighty-three compounds displaced more than 50% of [^3^H]-DTG (**Fig. 3A**), corresponding to a hit rate of 40% (**Fig. 3B**). Among the top 18 compounds ranked by [^3^H]-DTG displacement, 7 of 18 hits achieved K_i_ below 50 nM, with the best compound possessing a K_i_ of 14 nM (**Fig. 3C-D**), comparable to PB28 (K_i_ = 33 nM measured in the same assay), whereas in the prior campaign 6 of 21 compounds achieved Ki < 50 nM and the best Ki was 30 nM (**Fig. 3C-D**). The most potent compound’s docked pose forms key salt bridge interaction with D29 as well as an additional hydrogen bond with Q77 (**Fig. 3E**). Its dose-response curve is presented in **Fig. 3F**.

#### Comparative analysis across σ_2_ campaigns

Across the three σ_2_ campaigns, screening larger chemical space improved both hit rate and potency distribution. CombiDOCK achieved the highest hit rate, 67% versus 51% in the prior 490-million baseline, and improved the best potency from 30 nM to 6 nM, making its top ligand also about 5-fold more potent than the well-characterized σ_2_ ligand PB28 (Ki = 33 nM). MINT-Dock achieved a 40% hit rate with a best Ki of 14 nM, thus delivering comparable or improved potency relative to the prior 490-million campaign while requiring 22-fold less docking time and approximately 240-fold less storage (1 TB for synthon conformers versus 240 TB for full-molecule conformers).

### Prospective combinatorial docking and molecular generation against SLC transporters VMAT2 & VAChT

We next asked whether the scaling effect observed with σ_2_ generalized to challenging neurotransmitter solute carrier (SLC) transporters.^92–97^ We selected VMAT2 and VAChT as a paired test case because they are in the same family, share similar methodological constraints, and are both highly clinically relevant. They also adopt the major facilitator superfamily (MFS) fold, a widely conserved structural class within the SLC superfamily that remains underexplored by large-scale virtual docking. To our knowledge, few prospective docking campaigns at the >1-billion-compound scale, if any, have been reported for MFS transporters. Both are challenging targets for structure-based ligand discovery due to their extensive conformational dynamics,^98–101^ and historically limited structural information available for SLC proteins. Only recently have their high-resolution experimental structures been determined,^101,102^ enabling structure-based discovery of new chemotypes. This limited structural knowledge in the protein family makes pose prediction especially challenging. Notably, although AlphaFold3 (AF3) performs comparably to DOCK3 for σ_2_ (RMSD < 2Å), its performance does not generalize to VMAT2 or VAChT.^103^ Both DOCK3 and CombiDOCK accurately reproduced the cryo-EM binding poses of known ligands (RMSD < 2 Å), whereas AF3 failed to generate qualitatively correct binding modes (**Fig. S7A-B**). This trend held across a broader panel of 154 SLC transporters, revealing a systematic limitation of AF3: restricted generalizability to protein-ligand complexes underrepresented or absent in its training data (**Fig. S7C)**. In contrast, physics-based approaches such as DOCK3 and CombiDOCK maintained robust performance across these out-of-distribution targets, underscoring why advancing scalable physics-based docking remains conceptually important even for ligandable targets.

Beyond their methodological challenges, both targets present pressing therapeutic needs that motivate the discovery of new chemotypes beyond their existing privileged scaffolds. The Vesicular Monoamine Transporter 2 (VMAT2, SLC18A2) is an antiporter that actively loads monoamine neurotransmitters from the cytosol into synaptic vesicles.^104^ It is a key drug target for movement disorders, including tardive dyskinesia and chorea associated with Huntington’s disease.^105–107^ Although VMAT2 is a clinically validated target, important therapeutic gaps remain.^108^ Inter-patient variability in CYP2D6 metabolism can alter exposure and tolerability of current VMAT2 inhibitor scaffolds,^109^ motivating the search for new chemotypes with improved pharmacological properties. The Vesicular Acetylcholine Transporter (VAChT, SLC18A3) performs an analogous function, specifically loading acetylcholine into synaptic vesicles in cholinergic neurons,^110,111^. While it is a promising diagnostic target for Alzheimer’s disease, existing vesamicol-class ligands lack selectivity over σ receptors, limiting their clinical imaging utility.^112^ For both targets, structure-based screening of ultra-large chemical space offers a path to move beyond these long-established scaffold classes and identify fundamentally new chemotypes that can potentially address these pressing clinical and translational needs.

We conducted prospective screens against VMAT2 (PDB ID: 8T69, tetrabenazine-bound complex)^113^ and VAChT (PDB ID: 8ZMR, vesamicol-bound complex)^101^ across libraries of increasing size (3.6 million (DOCK3.8), 1.9 billion (DOCK3.8), and 103 billion (CombiDOCK and MINT-Dock)). We present the VMAT2 results below, followed by VAChT in the next section.

#### DOCK3 screens (3.6 million and 1.9 billion molecules)

Using full-molecule docking with DOCK3, we screened 3.6 million and then 1.9 billion compounds against VMAT2. Full-molecule docking of the 1.9-billion-compound library required 11 cluster-days on a 2,000-CPU-core cluster. Across these screens, we selected top-scoring compounds for experimental testing from the top-ranked subset (top 100K for 3.6M; top 1M for 1.9B) using the same post-docking filters detailed in **Methods**. We used a top-100K cutoff for the 3.6-million screen, rather than top 1M, because selecting 1M compounds from a 3.6M library would be both impractical and unnecessary; instead, we capped selection at the top 3% of the ranked list, already a generous cutoff given that prior in-stock screens typically select compounds from the top 0.8-1.0%.^35,39^ These campaigns resulted in the synthesis and testing of 65 and 111 compounds, respectively. In a [^3^H]-dopamine ([^3^H]-DA) uptake assay at 10 µM, VMAT2 hit rates were 23% in the 3.6-million screen (15/65 displaced >50% [^3^H]-DA) and 27% in the 1.9-billion screen (30/111 displaced >50% [^3^H]-DA) (**Fig. 3G-H**). In the 3.6-million screen, IC_50_ values ranged from 2 µM to 7 µM (six with IC_50_ > 1 µM, none with IC_50_ between 200 nM and 1 µM, none with IC_50_ < 200 nM). In the 1.9-billion screen, IC_50_ values ranged from 230 nM to 2 µM (one with IC_50_ > 1 µM, ten with IC_50_ between 200 nM and 1 µM, none with IC_50_ < 200 nM) (**Fig. 3I-J**).

#### CombiDOCK exhaustive screen (103 billion molecules)

CombiDOCK was used to screen 103 billion molecules against VMAT2, requiring 33 cluster-days on a 2,000-CPU-core cluster. This campaign resulted in the synthesis and testing of 109 compounds from the top 1 million ranked molecules. The hit rate further improved to 47% (51/109 displaced >50% [^3^H]-DA) (**Fig. 3G**). IC_50_ values ranged from 125 nM to 1 µM (one with IC_50_ = 1 µM, nine with IC_50_ between 200 nM and 1 µM, two with IC_50_ < 200 nM). The top compound has IC_50_ of 125 nM (**Fig. 3I-J**), whose docked pose preserves all key interactions with the VMAT2 receptor (E312, R189, F135) (**Fig. 3K**). Its dose-response curve is presented in **Fig. 3L**.

#### MINT-Dock guided search (103 billion molecules)

Starting from the synthons of the 103 billion Enamine REAL Space, MINT-Dock generated 15.6 million docked, synthesizable compounds for downstream processing in 4 days using approximately 95 CPU cores. 108 compounds were selected and tested from the top 1 million ranked molecules. Thirty-seven compounds displaced more than 50% of [^3^H]-DA (**Fig. 3G**), corresponding to a 34% hit rate **(Fig. 3G-H)**. We further characterized the top 13 hits with dose-response curves, obtaining an overall potency profile in which 5 of 13 compounds had IC_50_ < 1 μM and 12 of 13 had IC_50_ < 2 μM (**Fig. 3I-J**). The most potent compound had an IC_50_ of 126 nM, whose docked pose similarly preserves all key interactions with the VMAT2 receptor (E312, R189, F135) (**Fig. 3K**).

#### Comparative analysis across VMAT2 campaigns

Across the four VMAT2 campaigns, screening larger chemical space improved both hit rate and potency distribution. CombiDOCK achieved the highest hit rate, 47% versus 23% and 27% in the 3.6M and 1.9B brute-force campaigns, and improved the best potency from 230 nM in the 1.9B screen to 125 nM, approaching the potency of the FDA-approved VMAT2 inhibitor tetrabenazine (TBZ; IC_50_ = 92 nM, **Fig. 3L**). MINT-Dock achieved a 34% hit rate, exceeding the prior 1.9B brute-force campaign, with a best IC_50_ of 126 nM, also slightly better than the best compound from the 1.9B screen and likewise comparable to TBZ, while requiring 64-fold less docking computation and approximately 930-fold less storage.

Having established that both CombiDOCK and MINT-Dock improve ligand discovery outcomes for VMAT2, we next turned to VAChT, the second vesicular SLC transporter in our paired test case. As described above, existing vesamicol-class ligands, the privileged scaffold for VAChT, lack selectivity over σ receptors, limiting their clinical utility as imaging agents for Alzheimer’s disease.^114,115^ VAChT therefore presents not only a single-target discovery challenge but also a multi-target optimization problem: identifying compounds that are potent against VAChT while selectively avoiding σ_2_ activity, or alternatively, identifying dual-active compounds for polypharmacology applications. We conducted progressive single-target screens (3.6M, 1.9B, 103B) using DOCK3 and CombiDOCK, and additionally leveraged MINT-Dock’s multi-target capabilities for explicit selectivity and polypharmacology campaigns against VAChT and σ_2_.

#### DOCK3 screens (3.6 million and 1.9 billion molecules)

Using full-molecule docking with DOCK3, we screened 3.6 million and then 1.9 billion compounds against VAChT (PDB ID: 8ZMR, vesamicol-bound complex).^101^ These campaigns resulted in the synthesis and testing of 64 and 108 compounds, respectively. In a [^3^H]-vesamicol competition binding assay at 7.5 µM, hit rates were 25% in the 3.6-million screen (16/64 displaced >50% [^3^H]-vesamicol) and 28% in the 1.9-billion screen (30/108 displaced >50% [^3^H]-vesamicol) (**Fig. 4A**). In the 3.6-million screen, IC_50_ values ranged from 200 nM to the low µM range (two with IC_50_ > 1 µM, three with IC_50_ between 100 nM and 1 µM, none with IC_50_ < 100 nM). In the 1.9-billion screen, IC_50_ values ranged from 103 nM to the low µM range (two with IC_50_ > 1 µM, 13 with IC_50_ between 100 nM and 1 µM, none with IC_50_ < 100 nM) (**Fig. 4B**).

**Figure 4.**
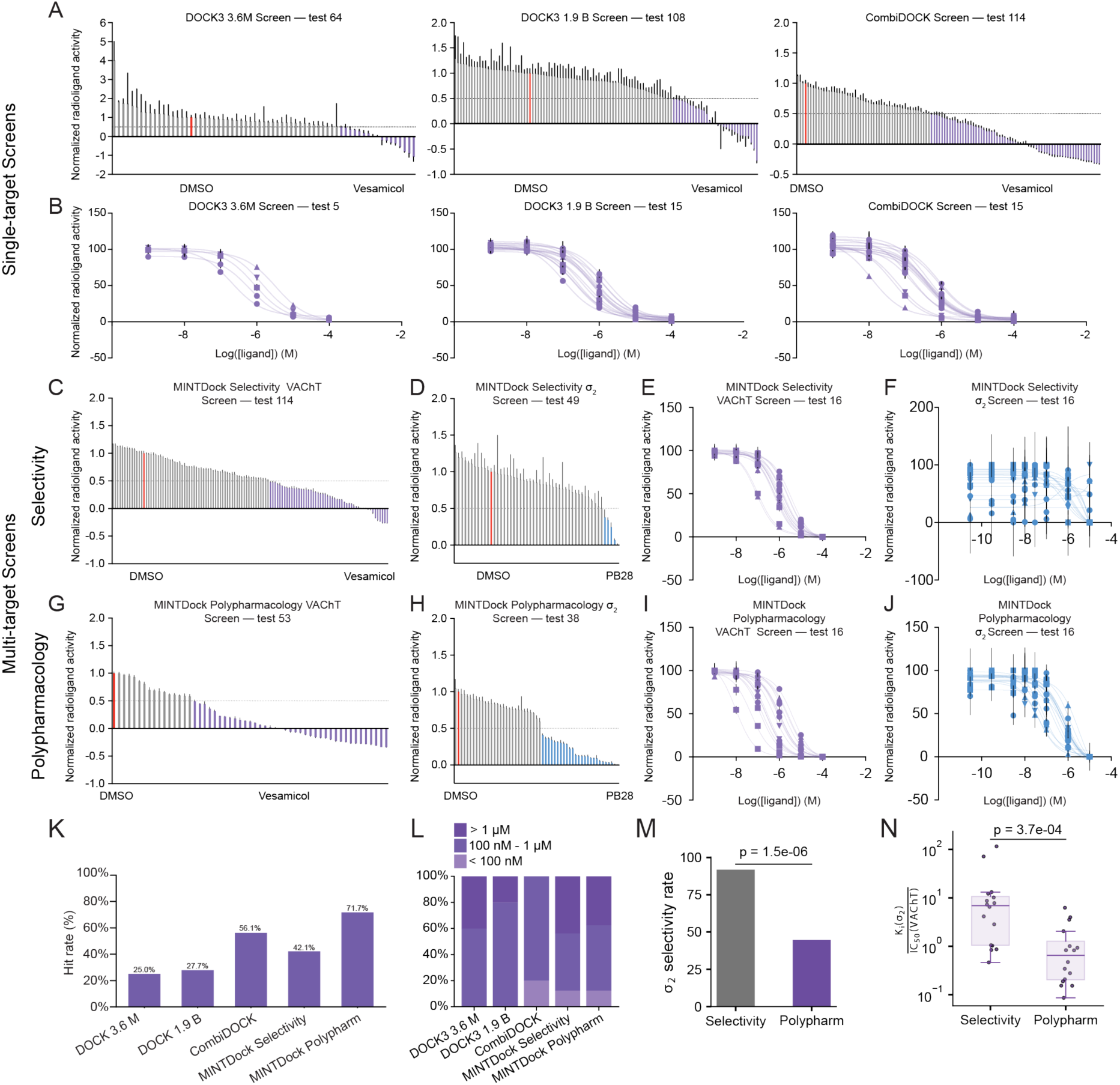
Summary of virtual screening results against VAChT. (A) VAChT compounds from the DOCK3 and CombiDOCK screens were tested in a [^3^H]-vesamicol uptake assay, using > 50% [^3^H]-vesamicol displacement as a hit threshold. 16 out of 64 tested compounds from the 3.6 million screen were hits, representing a hit rate of 25% (left). 15 out of 108 tested compounds from the 1.9 billion screen were hits, representing a hit rate of 28% (middle). 64 out of the 114 tested compounds were hits in the 103 billion CombiDOCK screen, representing a hit rate of 56% (right). (B) Dose-response curves of tested compounds from the DOCK3 and CombiDOCK screens. (C) 49 out of 114 tested compounds from the 103 billion MINT-Dock selectivity screen were found to be hits when tested in a [^3^H]-vesamicol uptake assay, representing a hit rate of 43%. (D) 45 of the 49 VAChT hits from the MINT-Dock selectivity screen displaced <50% [^3^H]-DTG in a [^3^H]-DTG binding assay. (E) VAChT dose-response curves of compounds from MINT-Dock selectivity screen. (F) σ_2_ dose-response curves of compounds from MINT-Dock selectivity screen. (G) 38 of the 53 tested compounds from the MINT-Dock polypharmacology screen were found to be hits when tested in a [^3^H]-vesamicol uptake assay, representing a hit rate of 72%. (H) 21 of the 38 VAChT hits from the MINT-Dock polypharmacology screen displaced >50% [^3^H]-DTG in a [^3^H]-DTG binding assay. (I) VAChT dose-response curves of compounds from MINT-Dock polypharmacology screen. (J) σ_2_ dose-response curves of compounds from MINT-Dock polypharmacology screen. (K) Hit rate steadily improves as library size increases. (L) The potency distribution shifts towards higher potency compounds as the screening size grows. (M) Selectivity hit rate comparison between the two MINT-Dock multi-target campaigns. P-value was calculated by the two-sample proportion Z-test. (N) Selectivity fold comparison between the two MINT-Dock multi-target campaigns. P-value was calculated by the two-sided Mann-Whitney U test.

#### CombiDOCK exhaustive screen (103 billion molecules)

CombiDOCK was used to screen 103 billion molecules against VAChT, requiring 26 cluster-days on a 2,000-CPU-core cluster. This campaign resulted in the synthesis and testing of 114 compounds from the top 1 million ranked molecules. The hit rate further improved to 56% (64/114 displaced >50% [^3^H]-vesamicol) (**Fig. 4A**). IC_50_ values ranged from 12 nM to 847 nM (none with IC_50_ > 1 µM, 12 with IC_50_ between 100 nM and 1 µM, three with IC_50_ < 100 nM). Notably, sub-200 nM ligands emerged only at the 103-billion scale. The top compound has an IC_50_ value of 12 nM (**Fig. 4B**).

#### MINT-Dock selectivity campaign (VAChT selective over σ_2_)

Beyond single-target screening, MINT-Dock’s multi-target capabilities enabled explicit optimization for VAChT selectivity over σ_2_: a key clinical objective that cannot be addressed by exhaustive single-target screening alone. MINT-Dock generated 12.9 million compounds for downstream processing in 3.5 days with approximately 110 CPU cores. With the same post-docking filtering criteria used for the CombiDOCK and prior campaigns (detailed in **Methods**), 114 were synthesized and tested for VAChT activity. In a [^3^H]-vesamicol displacement assay, 49 of 114 compounds displaced >50% of radioligand (**Fig. 4C**), corresponding to a 43% VAChT hit rate. To assess selectivity, we then tested these VAChT hits against σ_2_ using a [^3^H]-DTG displacement assay. Forty-five of 49 compounds displaced <50% of [^3^H]-DTG (**Fig. 4D**), indicating that 92% of VAChT actives were σ_2_ selective under this criterion. Overall, 45 of 114 synthesized compounds satisfied the selectivity profile (VAChT active and σ_2_ inactive), yielding an overall hit rate of 40%. We advanced 16 selectivity hits for dose-response curves against both targets. These compounds showed VAChT potencies spanning the sub-100 nM to low-micromolar range, with the two most potent compounds achieving VAChT IC_50_ = 86 nM (Z9424360192) and 100 nM (Z2406174633) (**Fig. 4E**). In parallel σ_2_ profiling, most compounds remained weak binders, with 9 of 16 exhibiting σ_2_ Ki > 10 μM, consistent with the intended selectivity task (**Fig. 4F**).

#### MINT-Dock polypharmacology campaign (VAChT plus σ_2_)

For the polypharmacology campaign, MINT-Dock generated 8.8 million compounds for downstream processing in 3 days with approximately 75 cores. 53 compounds were synthesized and tested. In the [^3^H]-vesamicol displacement assay, 38 of 53 compounds displaced >50% of radioligand (**Fig. 4G**), corresponding to a 72% VAChT hit rate. We then evaluated these VAChT hits against σ_2_, and 21 of 38 displaced >50% of [^3^H]-DTG (F**ig. 4H**), corresponding to a 55% σ_2_ hit rate among VAChT actives and yielding an overall polypharmacology hit rate of 21 of 53 (40%). Across the polypharmacology-tested compounds, higher VAChT displacement was associated with higher σ_2_ displacement (**Fig. S8A**), and vice versa (**Fig. S8B**), consistent with the expectation that the polypharmacology objective enriches dual-target binders rather than producing VAChT and σ_2_ activity independently by chance. Among the top 16 polypharmacology hits followed up with dose-response measurements, IC_50_ values on VAChT spanned from single-digit nanomolar to low micromolar, with the most potent compound reaching VAChT IC_50_ = 10 nM (Z9506560893) and multiple additional compounds in the sub-200 nM range (**Fig. 4I**). Consistent with the dual-target task, σ_2_ binding for this set also shifted into a more potent range, with Ki values extending to tens of nanomolar for the strongest binders (**Fig. 4J**).

#### Comparative analysis across VAChT campaigns

Across the five VAChT campaigns, screening larger chemical space improved both hit rate and potency distribution, while MINT-Dock further enabled objective-specific multi-target discovery (**Fig. 4K,L**). CombiDOCK achieved the highest single-target hit rate, 56% versus 25% and 28% in the 3.6M and 1.9B brute-force campaigns, respectively, and improved the best potency from 103 nM in the 1.9B screen to 12 nM, a roughly 150-fold improvement over [+]-vesamicol in the same assay. MINT-Dock achieved higher VAChT hit rates than the 1.9B brute-force campaign in both multi-target modes, 43% for the selectivity campaign and 72% for the polypharmacology campaign, with more than 80-fold less computational cost. Its best compounds reached 86 nM in the selectivity campaign and 10 nM in the polypharmacology campaign, while steering discovery toward distinct endpoints: the selectivity campaign yielded compounds with significantly higher selectivity indices, whereas the polypharmacology campaign produced dual VAChT/σ_2_ binders (**Fig. 4M,N**). Unexpectedly, even without explicit selectivity optimization, the top CombiDOCK compound, CSMS00228273820, exhibited a Ki of >10 µM for σ_2_, corresponding to >800-fold selectivity over σ_2_. This exceeds the selectivity of [+]-vesamicol (**Fig. 5A**), prior reported values for (±)-benzovesamicol (73-fold) and spiroindolines (<8-fold),^115^ and the best selectivity hit from the MINT-Dock campaign (116-fold, **Fig. 5A**). Together, these hit-rate, potency, and selectivity profiles show that MINT-Dock can steer discovery toward qualitatively distinct multi-target endpoints, while CombiDOCK can independently uncover highly selective compounds through exhaustive coverage of chemical space.

**Figure 5.**
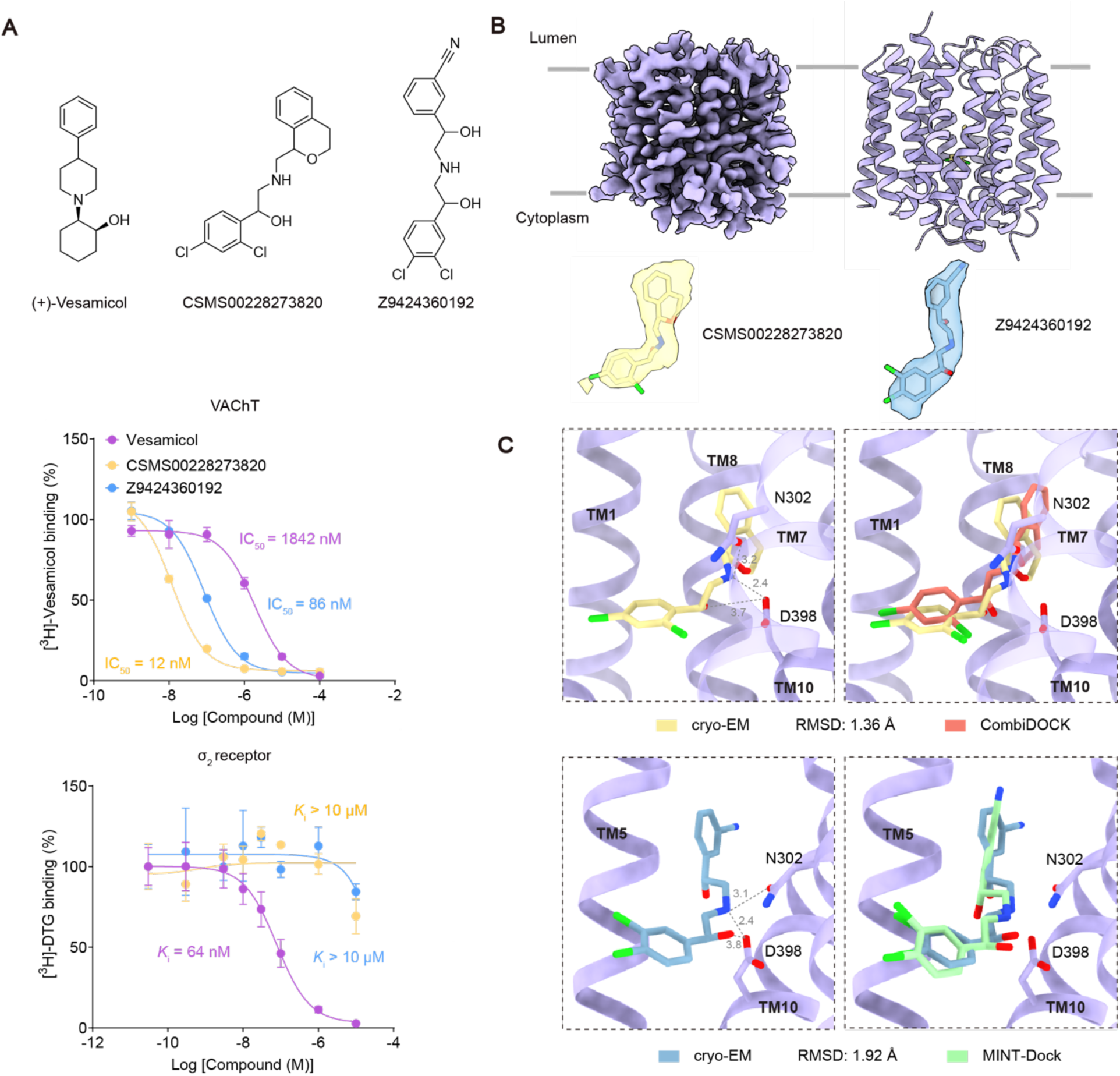
Structural determination of VAChT bound with ligand CSM00228273820 and Z9424360192. (A) The 2D structure of vesamicol, CSM00228273820 and Z9424360192, along with dose-response curves of vesamicol (blue), CSM00228273820 (purple) and Z9424360192 (yellow) against VAChT (top) and σ_2_ (bottom). (B) CryoEM density map of ligand-bound VAChT (top). The model of CSM00228273820 and Z9424360192 built into density is shown (bottom). (C) Experimental poses of CSM00228273820 (top) and Z9424360192 (bottom) are shown to conserve key interaction with D398. High-fidelity between the experimental pose and the CombiDOCK predicted pose of both CSM00228273820 and Z9424360192 is observed with an RMSD < 2 Å.

#### Structural validation of CombiDOCK/MINT-Dock-predicted poses

Unlike conventional docking that operates on pre-enumerated full molecules, CombiDOCK assembles synthons into full molecules on the fly, completely eliminating the enumeration step. Because the docking poses are constructed through this synthon-staged assembly rather than sampled from precomputed full-molecule conformers, it is important to validate that the resulting predicted poses are structurally faithful. We therefore selected the top compound from each method: the most potent from the CombiDOCK exhaustive screen and the most selective from the MINT-Dock selectivity campaign, and we determined cryo-EM structures of VAChT bound to each molecule.

For the CombiDOCK-derived compound CSMS00228273820 (IC_50_ = 12 nM, **Fig. 5A**), we obtained a 3.6 Å-resolution cryo-EM structure of its complex with VAChT in an outward-open state (**Fig. 5B** and **Fig. S9** and **Table S4**). The local resolution around the ligand-binding site reached approximately 3.3 Å, allowing unambiguous determination of the ligand pose and surrounding side-chain conformations (**Fig. S10A**). The experimentally resolved pose closely matched the CombiDOCK prediction (RMSD = 1.4 Å; **Fig. 5C**), confirming bidentate interactions with D398. In contrast, AF3 failed to accurately predict the binding mode (best RMSD of 3.8 Å; **Fig. S11A**). Combined with >800-fold selectivity over σ_2_, these data highlight its promise for future therapeutic development (**Fig. 5A**, bottom panel).

For the MINT-Dock-derived selective compound Z9424360192 (IC_50_ = 86 nM), we determined a 3.7 Å-resolution cryo-EM structure of its VAChT complex (**Fig. S12** and **Table S4**). Local resolution near the ligand-binding site was similarly high (3.3 Å), enabling clear pose assignment (**Fig. S10A**). Superimposition with the MINT-Dock prediction showed close agreement (RMSD = 1.9 Å; **Fig. 5C**), verifying bidentate interactions with D398 and additional bidentate contacts with N302. In this case, the top-confidence AF3 pose was also close to the experimental structure, with an RMSD of 2.2 Å (**Fig. S11B**). Across the three experimental VAChT structures (vesamicol, CSMS00228273820, and Z9424360192), the 33% success rate of AF3 pose reproduction followed the same overall accuracy trend shown in **Fig. S7C**.

The independent cryo-EM validation of both predicted poses confirms the structural fidelity of the shared combinatorial docking framework. We further performed MicroScale Thermophoresis (MST) to validate the binding of CSMS00228273820 to VAChT and to confirm the functional significance of the two key interaction residues, D398 and N302 (**Fig. S10B**).

#### Analogue optimization

To assess the capability of the combinatorial strategy for analogue optimization, we pursued analogue searches around top hits from both campaigns. For VMAT2, we applied a synthon-based approach in which close analogues were rapidly identified by searching matching synthons around parent hits. Across four primary CombiDOCK hits, 103 analogues were synthesized and tested, of which 72 (70%) exhibited >50% inhibition in the same primary assay described above. Dose-response follow-up identified more potent analogues for three of the four ligands, with potency improvements of up to 9-fold and a best IC_50_ of 77 nM (**Fig. S13A-E**), comparable to the FDA-approved VMAT2 inhibitor tetrabenazine. For VAChT, we searched Enamine REAL Space with [Small World](https://swp.docking.org/) for close analogs around the top CombiDOCK hit and two MINT-Dock selective hits. These analogue searches yielded high hit rates and identified affinity improvements while maintaining selectivity. Notably, two series converged on the same optimized scaffold, which reached a VAChT IC_50_ of 8 nM and 649-fold selectivity over σ_2_ (**Fig. S13F**), exceeding prior reported selectivity values for (±)-benzovesamicol (73-fold) and spiroindolines (<8-fold).^115^ Across five optimization series spanning both methods, three showed 8- to 12-fold improvements, supporting that the workflow can efficiently progress from initial hit discovery to improved potency without bespoke synthesis campaigns.

#### Scaffold novelty analysis

Bemis-Murcko scaffold analysis^116^ across the σ_2_, VMAT2, and VAChT campaigns showed that screening previously intractable chemical space with either CombiDOCK or MINT-Dock consistently uncovered previously unseen chemotypes (**Fig. S14**). For σ_2_, scaffold overlap between the smaller DOCK3 screen and the larger CombiDOCK and MINT-Dock screens was minimal, with only at most two pairwise shared scaffolds and just one scaffold shared across all three campaigns (**Fig. S14A-B**). Similarly low overlap was observed for VMAT2 and VAChT between the smaller in-stock/ZINC library screens performed with DOCK3 and the larger CombiDOCK and MINT-Dock campaigns, where shared scaffolds were rare and most tested scaffolds were unique to each screen (**Fig. S14C-H**). Even when CombiDOCK and MINT-Dock searched the same underlying chemical space, they recovered largely distinct scaffold sets, indicating that exhaustive and guided strategies sample different productive regions of that space (**Fig. S14D,H**). Together, this structural diversity across methods and objectives confirms that both exhaustive and guided approaches reveal novel chemotypes at previously inaccessible scale.

### Fragment-enrichment based virtual screening versus CombiDOCK/MINT-Dock: a comparative analysis

Both CombiDOCK and MINT-Dock operate on synthon-based chemical space, a representation shared by fragment-enrichment methods such as V-SYNTHES. However, the three approaches differ fundamentally in how they exploit this representation. Fragment-enrichment methods dock individual synthons to identify high-scoring anchor fragments, enumerate only products containing those anchors, and then dock the reduced product set at the full-molecule level. CombiDOCK instead docks capped synthons once and recombines cached poses to assemble and score all full molecules on the fly. MINT-Dock further couples this combinatorial docking engine with MCTS-guided search to concentrate docking on the most promising regions of chemical space, and its multi-target optimization additionally enables selectivity-driven optimization: a task that fragment-enrichment methods cannot address. We evaluated how fragment-enrichment performs relative to both methods, first retrospectively in recall of top-scoring molecules and then prospectively in recovery of experimental actives reported in this study.

To evaluate the fidelity of our approach with combinatorial libraries, we retrospectively benchmarked CombiDOCK against full-molecule docking across σ_2_, VMAT2, and VAChT using a 10-million-compound library constructed from 5,000 and 2,000 randomly selected Syn1s or Syn2s, respectively. Overall, CombiDOCK outperforms existing fragment-enrichment protocol adapted from V-SYNTHES by 7- to 20-fold in recall across all three targets, while maintaining close agreement with full docking rankings (Pearson/Spearman *R* = 0.67 to 0.89) (**Fig. S15A**, more details in **Methods**). While fragment-enrichment significantly reduces computation, it does so at the cost of losing context-dependent interactions that emerge when synthons are covalently joined, leading to poor overlap with full molecule docking results. By instead docking capped synthons only once and recombining cached poses, CombiDOCK preserves the full context of binding poses while avoiding redundant computations. For MINT-Dock, we compared it to the fragment-enrichment protocol above under matched-scale outputs for both σ_2_ and VMAT2. As shown in **Fig. S15B-C**, MINT-Dock’s guided search, which uses product-level docking feedback to refine its exploration, generates molecules that are not only well-scored but also more likely to satisfy the specific interaction patterns required for productive binding. **Fig. S16** further provides composition analysis that explains the increased enrichment for σ_2_ after interaction-based triage.

These differences translated into prospective performance (**Fig. S15D**; see **Methods**): across σ_2_, VMAT2, and VAChT, fragment-enrichment recovered only a small fraction of the true actives identified by CombiDOCK. Even under generous conditions, where the top 6 million molecules were enumerated and docked as full compounds, fragment-enrichment still missed more than 95% of the experimental hits from the CombiDOCK campaigns. Only two σ_2_ hits were recovered under the same chemical space. A similar pattern was observed for VMAT2, where fragment-enrichment identified only one hit (25%, 1/4) and no ligands with potency better than 1 μM. The VAChT campaign showed the same trend: fragment-enrichment yielded only five hits (38%, 5/13), with affinities remaining near the 1 μM range. Recall improved only when very large fractions of the library were enumerated and fully docked (**Fig. S15D** and **Table S5**). For VAChT, complete recovery of CombiDOCK actives required enumeration of more than 1 billion products, increasing the end-to-end cost to 179 cluster-days on 2,000 CPU cores, compared with 26 cluster-days for CombiDOCK.

### Computational efficiency of both methods

For Enamine REAL Space (103 billion molecules), we estimated that conventional full-molecule workflows would require approximately 18,000 CPU-days and 50,000 TB of storage for conformer preparation, whereas CombiDOCK reduced this to about 1 cluster-day and 1 TB by representing the library at the synthon level (**Fig. 1A**). Average docking speed improved from 1 s per molecule for full-molecule docking (DOCK3) to 0.07 s per molecule for CombiDOCK. This corresponds to >10-fold acceleration over conventional full-molecule docking, 14-fold acceleration over AutoDock Vina GPU (1 s per molecule), and at least 300-fold acceleration over recent deep-learning co-folding methods such as Boltz-2 (20 s per molecule) and AF3 (120 s per molecule).^103,117–119^ MINT-Dock further reduced cost by docking only selected synthon combinations, typically generating >10 million docked, synthesizable molecules in 3-4 cluster days using 75-110 CPU cores. Across prospective campaigns, this corresponded to 22-fold less docking computation than the prior 490M σ_2_ brute-force campaign, 64-fold less than the 1.9B VMAT2 campaign, and >80-fold less for the multi-target 1.7B VAChT campaigns. Rank-enrichment analyses against brute-force baselines showed strong concentration of well-scored molecules, including up to 660-fold rank enrichment for σ_2_, up to 130-fold enrichment for VMAT2, up to 200-fold enrichment for the VAChT selectivity task, and up to 700-fold enrichment for the VAChT polypharmacology task (**Fig. S17**), while maintaining or improving ligand-discovery outcomes relative to these prior billion-scale screens. MINT-Dock’s practical advantage extends beyond reduced docking time to a major reduction in storage burden: by replacing full-molecule conformations with synthon conformations, storage requirements decrease from approximately 240 TB (490M molecules for σ_2_) or 930 TB (1.9B molecules for VMAT2 and VAChT) to only 1 TB for synthon conformers, a reduction of up to nearly 1,000-fold.

The recently published DrugCLIP is able to score over trillions of ligand-protein interactions per cluster day using 8 GPUs, which has enabled genome-wide docking, discovering potent binders prospectively for three biological targets.^6^ However, extrapolating DrugCLIP’s reported scaling (see details in **Methods**) suggests that screening 100 billion molecules would require 200 cluster-days for retrieval plus 116 cluster-days for conformer generation, for a total of at least 316 cluster-days under optimistic assumptions. By contrast, CombiDOCK required 1 cluster-day for synthon preparation and 30 cluster-days for screening. DrugCLIP also failed to recover the canonical ligand-binding sites for σ_2_, VMAT2, and VAChT, and its top hits scored substantially worse than CombiDOCK/MINT-Dock candidate hits after re-docking (**Fig. S11C,D**).

To test whether the improved prospective outcomes from larger libraries could be explained by simple differences in ligand size or selection depth across campaigns, we analyzed ligand heavy-atom counts and performed matched comparisons under overlapping ranking constraints. Larger libraries continued to yield higher hit rates for σ_2_, VMAT2, and VAChT across increasing heavy-atom cutoffs and under shared-ranking analysis, whereas no correlation was observed between ligand size and binding affinity among active compounds (**Fig. S18**). These results indicate that the improved performance was not driven by larger ligands or selection depth, but instead reflects improved ligand discovery outcomes in larger chemical spaces, enabled by our newly developed methods.

## Discussion

Altogether, we present over 250 billion scored complexes (56 billion for σ_2_, 103 billion for VMAT2, and 103 billion for VAChT). This is almost 40-fold larger than that reported in lsd.docking.org (6.3 billion),^120^ a dedicated database for large-scale docking. Additionally, we also present experimental activities for 1300 molecules across the three targets. This experimental dataset includes both active and inactive compounds, representing successful docking predictions as well as false positives. Moreover, we see great value in these datasets for benchmarking and training ML models for virtual screening,^121,122^ and they may further support advances in computational protein-binder design.^123,124^ Nevertheless, not all ML approaches share equal promise for scaling to screening billions of molecules. Recent benchmarking on co-folding methods such as AF3^103^ and Boltz2,^119^ reveals that these models perform poorly on out-of-distribution targets relative to their training set,^125^ a result that is borne out in our own comparisons of AF3 with CombiDOCK. CombiDOCK’s strong performance in both retrospective benchmarks and prospective docking screens highlights the advantages of this physics-based method, which offers superior scalability and generalizability for prospective screening of ultra-large libraries. These advantages also enable CombiDOCK to serve a dual role as a docking engine to quickly generate large datasets of docking scores for benchmarking and training ML models. Looking forward, as new reactions and diverse building blocks are discovered, the modular nature of make-on-demand libraries will support continued combinatorial growth. In this context, and in line with the “bigger is better” paradigm, docking at the trillion-molecule scale becomes increasingly feasible through both methods. CombiDOCK can complete a one-trillion-molecule screen in just one year on 2,000 CPU cores, with further acceleration achievable through cloud resources such as AWS.^126^ These computational speedups extend beyond overcoming the limitations of brute-force molecular docking. Even for approximation strategies scaled to 1 trillion, such as ML-based or fragment-based docking, top 10% full-molecule docking remains necessary for optimal performance, making 100-billion-scale redocking computationally prohibitive. We believe CombiDOCK can serve as a practical and scalable docking engine to enable efficient docking of such enriched subsets of chemical space. This naturally leads to MINT-Dock, which leverages CombiDOCK’s pre-computed synthon conformations to perform on-the-fly sampling and generation during docking while maintaining its computational speedups, and further extends this capability to multi-target design objectives that would otherwise require multiple exhaustive screens. MINT-Dock further strikes a practical balance among existing generative approaches by combining four goals that are rarely achieved simultaneously: (i) 3D, structure-based discovery through physics-based docking in the binding site, (ii) strict synthesizability through combinatorial chemistry, (iii) computational efficiency through search that evaluates only a small, high-value subset of an ultra-large library and (iv) practical throughput, enabling generation of over 100 million docked, synthesizable designs from trillion-scale chemical library with 2-3 days using 2,000 CPU cores.

Several caveats should be noted. First, three-component reactions were handled here as pseudo-two-component reactions by pre-reacting dependent synthon sets; this worked well at the current scale and remains practical at the trillion-molecule scale, although a more general hierarchical treatment of three-component assembly will become important as libraries continue to grow beyond 10 trillion molecules. Second, our multi-target studies used scalar design functions, which worked well here because the objectives were naturally commensurate, namely docking scores produced by the same docking method, making additive or monotonic combinations reasonable. Although we did not rely on explicit multi-objective MCTS for the multi-target studies presented here, MINT-Dock includes a multi-objective MCTS mode for settings in which maintaining the Pareto front is necessary.

While these limitations exist, they do not diminish the key conclusions drawn from this study. CombiDOCK and MINT-Dock together define a unified framework for navigating the trillion-molecule frontier of drug discovery. For single-target campaigns, the two methods can be deployed in a staged workflow: MINT-Dock first provides efficient directed exploration to rapidly identify promising chemotypes with modest computational resources, followed by CombiDOCK’s exhaustive screen of the full combinatorial space to maximize hit rates and ligand potency, as demonstrated by CombiDOCK’s consistently higher hit rates and best-in-series affinities across all three targets in this study. For multi-target objectives, including selectivity, polypharmacology, and multi-property design, MINT-Dock’s ability to embed explicit design functions into the search process makes it the method of choice, enabling discovery modes that are otherwise computationally out of reach even at the billion-molecule scale, let alone at the trillion-molecule frontier.

## Methods

### Preparation of DOCKable files for synthon enumeration library

The reactions and corresponding synthons comprising the enumeration library (the version of April 2023) were provided by Enamine. This database consists of 304 chemical reactions with 2.3 million synthons, theoretically enabling the enumeration of 36 billion compounds without considering stereoisomer expansion. All the reactions in the database can be separated into two categories: two-component and three component reactions. Since the combination of synthons in three component reactions has the dependency that one synthon is the linker connecting other two synthons, we simplified them into “pseudo two component reactions” by merging the linker synthon with one of the other two synthons. Moreover, we excluded 4 chemical reactions that generate the covalent compound libraries.

At the synthon level, to ensure the retention of reaction-formed functional groups, all the synthons in the database were capped according to their corresponding reaction-formed functional groups. These capping groups also served as chemically meaningful junctions for aligning pairwise synthons and subsequently reconstructing full molecules during docking. The capped synthons were represented in SMILES strings and were subsequently used to generate hierarchical trees of precomputed conformers in DB2 format.^127^ The ligand conformer preparation pipeline involved protonation state and tautomer enumeration using Jchem (v.22.22.0, https://chemaxon.com/), stereoisomers expansion via RDKit (v2023.09.1),^128^ three-dimensional structures using Corina (v.4.4.0026, https://mn-am.com/products/corina/), and conformational ensemble generation with OMEGA (v.2019.10.2, https://www.eyesopen.com/omega).^129^ Atomic partial charges and desolvation energies^130,131^ were computed as previously described.^132^

The resulting curated library of 3D capped synthons enables the theoretical construction of 103.2 billion compounds: 5.5 billion from two component reactions and 97.7 billion from pseudo two component reactions. Applying a heavy atom count threshold of 34 for pseudo two component products reduced the final enumerated docking library to approximately 56 billion compounds. For the σ_2_ receptor, we screened these 56 billion compounds, while for VMAT2 and VAChT, we screened the whole library of 103.2 billion compounds.

### Combinatorial docking approach

Our combinatorial docking approach, implemented in the version 6.14 of DOCK6,^45^ follows a hierarchical sampling strategy using capped synthons that sequentially assemble into final products. The process begins by docking a hierarchical tree of precomputed conformers of the *first* synthon (Syn1) into the target protein pocket, generating multiple Syn1 docking poses per orientation (this is a tunable docking parameter called *number per first search*, default value is 10). The hierarchical tree conformer of the *second* synthon (Syn2) is then grown onto these docked Syn1 poses to generate feasible docking poses for the complete molecule (governed by another tunable docking parameter called *number per second search*, default value is 5). Each resulting pose is scored using the DOCK 3.5 scoring function (ChemGrid Score),^50^ and the best-scoring pose of each full molecule is retained for VS. The combinatorial docking parameters for CombiDOCK are illustrated in **Table S6**, and the technical details were presented in the later sections.

### Docking models and parameters for large-scale docking

The σ_2_ receptor bound to cholesterol (PDB ID: 7MFI),^27^ the VMAT2 receptor bound to tetrabenazine (PDB ID: 8T69),^113^ and the VAChT receptor bound to vesamicol (PDB ID:8ZMR)^101^ were used as the 3D structural input for the σ_2_, VMAT2, and VAChT docking campaigns, respectively. In the σ_2_ receptor docking model, the structure was prepared as previously described. In the VMAT2 docking model, 45 matching spheres were calculated around the tetrabenazine ligand atoms, and AMBER united atom charges were assigned to the receptor structure. The volume of the low dielectric and the desolvation volume was extended out 1.2 Å and 0.1 Å, respectively, from the surface of protein using spheres calculated by SPHGEN. Energy grids were pre-generated with AMBER force fields using CHEMGRID for van der Waals potential,^133^ QNIFFT^134^ for Poisson-Boltzmann-based electrostatic potentials and SOLVMAP^131^ for ligand desolvation. This docking model was evaluated for its ability to enrich 5 known VMAT2 ligands (DTBZ, TBZOH, tetrabenazine, valbenazine, deutetrabenazine) over 250 property-matched decoys. Decoys are unlikely to bind to the receptor because despite their similar physical properties to known ligands, they are topologically dissimilar. The docking model described above was able to achieve a logAUC of 2.18 and to recover the crystal pose of tetrabenazine with RMSD values of 0.39 Å. We also constructed an “extrema” set of 28,086 molecules using the DUDE-Z webserver (http://tldr.docking.org) ^86^ to ensure that molecules with extreme physical properties were not enriched. This docking model enriched close to 94% monocations among the top 1,000 ranking molecules. Since the docking grids were originally generated using DOCK 3.7 Blastermaster for DOCK3, we converted them to DOCK6 readable grids using the method described by Balius and colleagues^135^ and using their script (available here: https://github.com/tbalius/teb_docking_test_sets.git).^84^ The VAChT docking model was similarly prepared as described above. Matching sphere perturbation was skipped. The low dielectric and desolvation volumes were extended out 1.7 Å and 0.3 Å, respectively. Energy grids were pre-generated using CHEMGRID,^133^ QNIFFT,^134^ and SOLVMAP.^131^ Using retrospective docking, the docking model was evaluated on how well it enriched four known VAChT ligands (vesamicol, CHEMBL1089205, CHEMBL20730, CHEMBL3587324) over 200 property-matched decoys. The model achieved a high logAUC of 64.26, and successfully recapitulated the crystal pose of vesamicol with an RMSD value of 1.61 Å. An “extrema” set of 79,815 molecules was generated and docked, revealing over 92% of the top 1,000 ranking molecules to be monocations. Finally, the scoring grids from the docking model, generated using DOCK3.7 Blastermaster, were converted into a DOCK6 readable format using the method/script described above.

Because CombiDOCK evaluates full molecules through on-the-fly assembly guided by pre-docked synthon ensembles, performance depends on two sampling parameters: the number per first search (how many top poses of the first synthon per orientation are retained for potential extension to partner synthons, averaging 2,500 orientations per molecule) and the number per second search (multiplied by the number per first search to determine how many top poses of the fully combined molecule are passed into the minimizer). On retrospective σ_2_ and VMAT2 datasets, docking score distributions and Pearson correlations with full docking rose sharply at small values of number per first search and then plateaued (**Fig. S6A-C**). Similar trends were observed as number per second search increased (**Fig. S6D-F**). Specifically, performance converged after sampling the top 10 first-synthon poses per orientation and the top 10 combined molecules per minimizer. Since larger parameter values proportionally increase docking runtime, for the three prospective campaigns below, we therefore set the number per first search to 10 and the number per second search to 5, which together capture near-maximal agreement with full docking (Pearson *R* = 0.86 for σ_2_ and 0.74 for VMAT2) while minimizing computational cost.

### Benchmarking datasets for evaluating CombiDOCK performance

To evaluate the performance of CombiDOCK, we first benchmarked it against conventional full-molecule docking using three curated ligand-decoy datasets: σ_2_, VMAT2, and VAChT. The σ_2_ dataset comprised 10 known ligands and 550 property-matched decoys derived from a previous study. The VMAT2 dataset consisted of 5 actives and 250 matched decoys, as described above in the VMAT2 docking model evaluation. The VAChT dataset contained 4 active compounds and 200 decoys. After stereoisomer enumeration and successful 3D conformer generation, all molecules were stored in DB2 format for full-molecule docking. For CombiDOCK benchmarking, each molecule was decomposed into pairwise capped synthons according to common reaction types, such as ether, amide, or sulfonamide linkages. These capped groups served as chemically meaningful junctions for aligning synthons and reconstructing complete molecules during the docking process. To ensure conformer consistency, 3D conformers of full molecules were used to derive the conformers of their corresponding synthons, which were also stored in DB2 format for combinatorial docking.

It should be noted that 57 molecules in the σ_2_ dataset—including 3 actives and 54 decoys—could not be feasibly decomposed into synthons. For these compounds, full molecule docking scores were directly incorporated into the CombiDOCK benchmarking analysis. Additionally, we evaluated the impact of the key docking parameters on docking performance. As shown in **Fig. S6C**, increasing the *number per first search* parameter improved the correlation between CombiDOCK and full-molecule docking scores. To balance computational cost and accuracy in large-scale docking campaigns, this parameter was set to a default value of 10. Similarly, we explored the *number per second search* parameter and set it to a default value of 5 (**Fig. S6F**).

Moreover, to assess generalizability beyond these three targets, we extended the evaluation to 43 additional protein targets from the DUDE-Z benchmark,^86^ each comprising actives and property-matched decoys at a 1:50 ratio. For consistency, 3D conformers of the full molecules were generated and subsequently decomposed into pairwise capped synthons for combinatorial docking.

### MINT-Dock implementation

The Enamine REAL Space used in this study is composed of 2.3 million synthons and encompasses nearly 103 billion small molecules. To fit three-component reactions to the CombiDOCK format, we converted three-component reactions to pseudo-two component reactions, which raises the total number of synthons to around 6.3 million. Since one building block can potentially react in multiple reactions, we mapped each unique (building block SMILES, reaction_id) pair to a unique identifier. In the following, we will use the term “synthon” to refer to each (building block SMILES, reaction_id) pair with its own identifier. We then cap each synthon according to each reaction it will participate in. Briefly, we replaced the terminal dummy atom with a connecting functional group corresponding to each reaction. We then used DOCK6 to compute the docking score of each synthon. If a synthon has multiple conformers with different docking scores, we took the minimum (most favorable) score. We capped the minimum synthon score to -10 and set a score of 10 for all synthon scores worse than -10. Three-component reactions were converted to two-component reactions, and the resulting synthons were followed by the same preprocessing procedure.

In the actual MCTS, each node is scored by a scoring function, which is a function of the node itself and the rollout number N, similar to the Synthemol implementation:^74^ 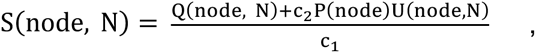 where for all non-product layer, Q(node, N) = ∑_cycles_ q(node, N) = ∑_cycles_ ∑_child nodes_ q(child node, N), which represents the exploitation score. If node is the product layer, then q(node, N) = f(S_node_, C_node_) = S_node_ – c_3_|C_node_ − C_empirical_|. S_node_, C_node_ are the docking score and the formal charge of the product contained in that node, and C_empirical_ is the empirical charge for a good molecule that could reasonably reach the target in vivo and bind with the target. c_3_ is a penalty hyperparameter that tunes how much penalty is placed on deviation in the product charge. 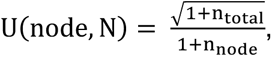 where n_node_ is how many times the node has been visited and n_total_ = ∑_sibling_ n_sibling_. This represents the exploration score followed from a prior MCTS scoring function.^136^ P(node) is the average of precomputed scores of synthons contained in that node, and c_2_ is an explore weight that tunes how much emphasis is placed on exploration versus exploitation. Finally, c_1_is a normalization hyperparameter that scales S(node, N). Since the DOCK6 score is retrieved at 6 decimal points precision, we also round S(node, N) to 6 decimal points.

We set the first layer nodes to the synthons that successfully docked in the target with a docking score below -10. In the first layer, we adopted a rolling visit strategy, where the tree first randomly selects a fraction of all available nodes for further expansion, which allows for more global explorations and provides speed-up for the tree search process. The tree then keeps the nodes that are estimated to have enough synthon_2_ to expand in the second layer and selects N_1_ (=500 or 1000) nodes with highest scores to continue in the second layer. In the second layer, for each selected first layer node, the tree searches for nodes that have synthon_2_ entries compatible with the synthon_1_ in that node, excludes nodes that have already been visited, and selects at most N_2_ (=50 for two-component reactions, =20 for three-component reactions) nodes with highest scores to continue to a single product layer, which then proceeds to parallel CombiDOCK calculation of all N_1_xN_2_ synthon combinations, where N_1_ Slurm jobs are submitted, each completing CombiDOCK calculation of one synthon_1_ with at most N_2_ synthon_2_. In practice, each round usually generates less than N_1_xN_2_ synthon combinations due to insufficient number of expandable synthon_2_ from a portion of selected reaction groups. After DOCK6 score has been computed for all products, the best score for each unique synthon combination, considering all possible conformations, is backpropagated to each parent layer to update n_node_ and q(node, N). We set c_emprical_ = +1 and a high c_3_ = 50 to penalize di-cations with favorable DOCK scores for all prospective screening campaigns.

For MINT-Dock for two targets, the DOCK6 scores of each synthon for each target are precomputed, and the score function becomes 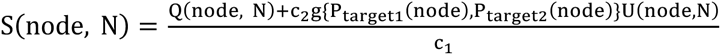 and for all non-product layer, Q(node, N) = ∑_cycles_ q(node, N) = ∑_cycles_ ∑_child nodes_ g{q_target1_(child node, N), q_target2_(child node, N)}. The expression of P and q for each target is the same in the single target case. g is a function that can be tuned according to the purpose of MCTS. For poly-pharmacology assays, we set 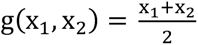 and for selectivity assays where we want the selected compounds to favor target_1_ over target_2_, we set 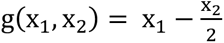 to balance between favorable docking scores for target_1_ and large docking score difference between target_1_ and target_2_. For poly-pharmacology screens, we also differentiate between the two targets by setting a primary target (target_1_) and an auxiliary target (target_2_). For compounds that are unable to be docked in the pocket of the primary target, their auxiliary target score is also set to a default low value.

Since MINT-Dock utilized a dynamic Slurm scheduling procedure to complete parallel docking, we approximated the average number of cores utilized for one complete MINT-Dock screen by the total DOCK time divided by the sum of maximum DOCK time across all Slurm jobs submitted per cycle. We summed the average number of utilized cores for two-component and three-component runs to get a total average number of utilized cores for single-target screens.

### MINT-Dock benchmarking

For this benchmarking, we compared the performance of MINT-Dock sampled molecules with randomly sampled molecules from an Enamine make-on-demand library of 141 reactions (131 two-component, 10 three-component).

For each target, we computed the dock score of each capped synthon for setting the initial heuristic strategy for MINT-Dock search with full molecule DOCK6 HDB docking.^84^ We capped the minimum synthon score to -10 and set a score of 10 for all synthon scores worse than -10. Here, we do not impose any charge penalty. We ran 60 cycles (1.5 million rollouts) for two-component reactions and 10 cycles (0.1 million rollouts) for three-component reactions with an explore weight of 0.001, hence constraining the theoretical maximum number of the generated compounds to roughly half of the random subset’s size. Relevant parameters are included in **Table S7.** For each target, MINT-Dock sampled between 3.2 to 9.0 million stereogenic molecules and eventually generated between 1.1 to 1.5 million compounds.

To procure the docking scores of random molecules, we curated 2.9 million unique compounds identified by synthon combinations from these 141 reactions totaling 5.3 million stereogenic molecules from full molecule db2 generation for DOCK6 HDB and 7.4 million stereogenic molecules from synthon db2 generation for CombiDOCK, while keeping the number of unique synthon combinations contributing to each reaction proportional to the full library. We then docked these molecules with either CombiDOCK or DOCK6 HDB. The adjusted enrichment ratio was calculated by the formula 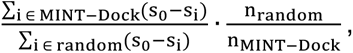 where s_0_ is the chosen dock score threshold, s_3_ is the best dock score for compound i, and n_MINT-Dock_ and n_random_ are the number of synthon pairs generated by MINT-Dock and by random subset respectively. For gross evaluation of MINT-Dock enrichment, we defined the score threshold for each target as max(s_MINT-Dock_, s_random_), where s_MINT-Dock_ is the docking score of the 1,001th ranked compound from MINT-Dock, and s_random_ is the docking score of the 2nd ranked compound from the random sample. To estimate the confidence interval, we performed bootstrapping with replacement on the MINT-Dock molecule scores and random molecule scores respectively. Namely, for each bootstrapping iteration, we calculate the adjusted enrichment using the pair of bootstrapped MINT-Dock sample and bootstrapped random sample. We repeated this process 1,000 times to gather an overall distribution to estimate the confidence interval.

### MINT-Dock backpropagation comparison

We tested whether backpropagation improves MINT-Dock’s ability to generate favorable molecules over successive search cycles. Measuring affinity improvement directly would be impractical, so we used docking score as a tractable proxy for search progress, since docking score correlates with experimental hit rate in three systems (AmpC beta-lactamase, D4 dopamine and σ_2_ receptors) in prior studies^91^ and remains the primary selection criterion in docking campaigns. We compared the number of good compounds between MINT-Dock with backpropagation implemented and MINT-Dock without backpropagation implemented, where in each rollout, the tree search is based on precomputed synthon scores only, on σ_2_ and VMAT2 receptors and on two multi-target objectives (**Fig. S1**). From the final pool of generated molecules, we compared the number of generated compounds with docking scores better than three given score thresholds respectively. The most lenient score threshold was chosen as -45 kcal/mol on the basis that this is the docking score at which the hit rate versus docking score curve for σ_2_ starts to show gradient in a previous large-scale virtual screening campaign.^27^ The most stringent threshold was chosen as -56.5 kcal/mol and -53.5 kcal/mol for σ_2_ and VMAT2 respectively based on the docking score distribution of the molecules from a 2 billion monocation library screen with CombiDOCK, where -56.5 is roughly the lowest score of the top 300,000 ranked compounds for the σ_2_ docking campaign, and -53.5 is roughly the lowest score of the top 1 million ranked compounds for the VMAT2 docking campaign. For multi-target comparison, the criterion for the number of good compounds was VAChT docking score below -45, σ_2_ docking score above -40 for selectivity, or both targets docking score below -45 for polypharmacology. Backpropagation consistently increased the yield of well-scoring molecules across multiple score cutoffs, and the advantage became more pronounced as the number of cycles increased, reaching 1.2 to 2.2-fold at the most stringent thresholds (**Fig. S2**). For selectivity (VAChT selective over σ_2_), we considered a compound well-scored if its VAChT docking score was < −45 kcal/mol and its σ_2_ docking score was > −40 kcal/mol. For polypharmacology, we considered a compound well-scored if its docking score was < −45 kcal/mol against both VAChT and σ_2_. Under these definitions, backpropagation improved screening results for well-scored two-component compounds to 2.4-fold in both selectivity and polypharmacology campaigns (**Fig. S2**). Notably, backpropagation produced a smaller gain for σ_2_ than for VMAT2, even though σ_2_ yielded more well-scoring compounds overall. We hypothesized that this reflects how well synthon-level docking approximates full-molecule docking for each system. Consistent with this, the root mean square deviation (RMSD) between synthon and full-molecule poses is generally lower for σ_2_ than for VMAT2 (**Fig. S3A–B**), indicating that precomputed synthon scores are a more reliable proxy for full-molecule scores in the σ_2_ campaign. This difference is especially pronounced for first-layer synthons (**Fig. S3C–D**), where the tree is widest and early selection is strongest, which suggests that the σ_2_ performance relies more on synthon priors and reduces the incremental benefit of correction by backpropagation. In contrast, for VMAT2, larger synthon-to-product pose discrepancies make product-level docking feedback more informative, leading to a stronger backpropagation effect.

### MINT-Dock synthon versus full-molecule RMSD calculation

For both σ_2_ and VMAT2, we took the top 300,000 ranked compounds out of their respective 10 million rollout MINT-Dock runs for two-component reactions after filtering for formal charges greater than +1.1 and removing compounds with duplicate SMILES. We then separated each full molecule into their synthon_1_ and synthon_2_ part poses using Chimera 1.18,^137^ maintaining the full structural information of the non-cap regions of the synthons. We calculated the RMSD between the heavy atom of each synthon component of the full molecule pose and those of the original best scoring capped synthon pose and then take either the first layer synthon RMSD or the minimum of the two RMSDs, one for each synthon component, for evaluating the distribution.

### MINT-Dock comparison with ML-MCTS

To assess whether coupling MCTS to physics-based docking improves generation relative to an ML surrogate, we compared MINT-Dock to an ML-guided MCTS baseline (“ML-MCTS”, adapted from Synthemol^74^). For both σ_2_ and VMAT2, MINT-Dock produced substantially more well-scored molecules exceeding any given score threshold, yielding higher enrichment across thresholds (**Fig. S4**). For VMAT2, we docked approximately 10 million randomly selected molecules from ZINC with DOCK6, 590,011 molecules had scores below -10, which we used to train a Chemprop model for predicting the scores of full molecules.^138^ For σ_2_, we docked approximately 2.5 million randomly selected molecules from ZINC with DOCK6 and selected 600,000 molecules that had scores below -10 to train a Chemprop model for predicting the scores of full molecules. We used these models to predict the scores for each unique SMILES to use to populate each synthon node. We adapted the Synthemol architecture so that it generates 10 molecules per cycle rather than 1 molecule per rollout to speed up the generation process. We generated 979,991 molecules for σ_2_ and 981,684 molecules for VMAT2 with 1 million rollouts on the two-component space with ML-MCTS. We generated 727,545 molecules for σ_2_ and 849,043 molecules for VMAT2 with 1 million rollouts on the two-component space with MINT-Dock. We then used CombiDOCK to calculate the actual scores of these molecules and compare between the two methods. The adjusted enrichment ratio was calculated by the formula presented in comparison with random docking.

### MINT-Dock-based screening campaigns

For all prospective screening campaigns, we accounted for the fact that the desired small molecule binders will need to permeate through cell membranes and cannot have high charges by assigning any molecule with a charge greater than +1 a penalty that is an affine function of its charge during backpropagation as described previously.

For σ_2_, we ran 400 cycles (10 million rollouts) on the two-component space and 200 cycles (2 million rollouts) on the three-component space, which sampled 120 million stereoisomers in total and generated 8.3 million and 2.0 million compounds respectively. For VMAT2, we ran 600 cycles (15 million rollouts) on the two-component space and 200 cycles (2 million rollouts) on the three-component space while filtering for high strain energy (total torsion strain energy of less than 8 units, maximum strain energy per torsion angle of less than 1.5 units) during docking to prevent the generation of many strained poses in the narrow pocket of VMAT2, which sampled 58 million stereoisomers in total and generated 13.6 million two-component and 2.0 million three-component compounds respectively. For VAChT versus σ_2_ selectivity assay, we ran 300 cycles (15 million rollouts) on the two-component space, which sampled 33 million stereoisomers and generated 12.9 million compounds. We also ran 100 cycles (2 million rollouts) on the three-component space for the purpose of backpropagation comparison only. For VAChT plus σ_2_ poly-pharmacology assay, we ran 400 cycles (10 million rollouts) on the two-component space only, while setting VAChT as the primary target, which sampled 31 million stereoisomers and generated 8.8 million compounds. We also ran 100 cycles (2 million rollouts) on the three-component space for the purpose of backpropagation comparison only. Relevant parameters are included in **Table S8-S9.** After the tree search had completed, we collected the compounds in the product layer. For backpropagation evaluation, we only performed a charge filter by removing compounds with formal charge greater than +1.1. For downstream pipeline of hit picking, we performed more elaborate filtering. In the single target setting, we filtered out compounds with undesirable charges and grouped the compounds by SMILES and selected the molecule with the most favorable adjusted docking score for each SMILES to maintain the diversity of the compounds for further hit picking.

### Post-docking processing

For σ_2_, the top-ranking 300,000 compounds from the 56-billion-compound CombiDOCK screen were retained for post-docking filtering. To remove molecules similar to known ligands, the ECFP4-based Tc value was computed against 2,232 σ_1/2_ ligands in ChEMBL (https://www.ebi.ac.uk/chembl/) and 574 σ_2_ ligands from S2RSLDB (https://www.researchdsf.unict.it/S2RSLDB). Compounds with an Tc ≥ 0.35 were removed, leaving 266,584 molecules. To prioritize poses that capture essential interaction with relevant residues, the remaining molecules were filtered using LUNA-based interaction calculations.^139^ Molecules that capture key interactions with D29 were retained. An additional constraint required fewer than 3 unsatisfied hydrogen bond acceptors and no unsatisfied hydrogen bond donors, narrowing the set to 91,574 compounds. To ensure a diverse set of compounds to select from, the remaining compounds were then clustered by their LUNA interaction fingerprints using the Butina algorithm (Tc ≥ 0.68), and cluster representatives were visually inspected. A total of 137 representatives were selected, and 122 were synthesized and experimentally tested.

The top 300,000 ranked compounds from the MINT-Dock screening were similarly retained for downstream strain filtering (total torsion strain energy of less than 8 units, maximum strain energy per torsion angle of less than 3 units, leading to 229,910 compounds), novelty filtering (Tanimoto similarity with known actives < 0.35, leading to 178,170 compounds), and interaction filtering with LUNA^139^ on compounds that form a salt-bridge/hydrogen-bond with D29, leading to 68,497 compounds. These filtered compounds were then clustered by their LUNA interaction fingerprints using the Butina algorithm (Tanimoto similarity ≥ 0.68), and cluster representatives were visually inspected. 225 were selected, and 206 were synthesized and tested. Because the same selection criteria were applied, 27 of the 206 compounds tested in the MINT-Dock campaign were also tested in the CombiDOCK campaign: Z9322686774, Z9322686448, Z9322686929, Z9322686891, Z9322686649, Z9322686503, Z9322686489, Z9322687006, Z9322686784, Z9322687611, Z9322686694, Z9322687001, Z9322687500, Z9322686991, Z9322687037, Z9322686403, Z9322687159, Z9322686415, Z9322686618, Z9322686480, Z9322687461, Z9322686557, Z9322687562, Z9322686657, Z9322686646, Z9322687053, and Z9322686947.

For both VMAT2 and VAChT, the top-ranking compounds from each brute-force screen were retained for post-docking filtering (top 100K for 3.6M; top 1M for 1.9B and CombiDOCK 103B). Compounds with an ECFP4 Tc ≥ 0.35 to known ligands (18 ligands for VMAT2; 38 ligands for VAChT) were removed, leaving 97,740, 978,464, and 993,861 molecules for VMAT2, and 97,693, 958,021, and 983,477 molecules for VAChT, respectively. The remaining molecules were filtered using LUNA-based interaction calculations, retaining only compounds that preserve target-specific bond interaction: for VMAT2, a key hydrogen bond with E312 and an additional hydrogen bond with R189, and a π-stacking interaction with F135; for VAChT, a key hydrogen bond with D398, and at least one additional hydrogen bond with S225, Y432, N302, or N47. An additional constraint required fewer than 3 unsatisfied hydrogen bond acceptors and no unsatisfied hydrogen bond donors, narrowing the sets to 282, 6,127 and 14,701 for VMAT2, and 2,397, 9,356 and 87,428 for VAChT. The filtered compounds were then clustered by their LUNA interaction fingerprints using the Butina algorithm (Tc ≥ 0.75), and cluster representatives were then visually inspected. Poses were selected for good shape complementarity to the binding site, consistency with established interaction patterns for known binders and excluded if they displayed structural artifacts, such as improper torsional geometries. For VMAT2, preference was given to poses that positioned a protonatable amine toward the conserved acidic residue E312, and maintained compact aromatic scaffolds, and formed well-defined π stacking interactions with F135, while avoiding strained conformations and improper placement of key functional groups. For VAChT, preference was given to poses featuring ring systems over flexible chains, as well as those exhibiting a bidentate binding mode with the binding site. After visual inspection, 78, 122, and 128 compounds were selected for VMAT2 and 83, 130, and 153 for VAChT. For VMAT2, 65, 111, 109 compounds were synthesized and tested experimentally. For VAChT, 64, 108, and 114 were synthesized and tested.

For VMAT2 MINT-Dock screening, we selected the top 1 million ranked compounds for downstream novelty filtering (Tanimoto similarity with 18 known actives < 0.35, leading to 959,047 compounds) and interaction filtering with LUNA on compounds that form a salt-bridge/hydrogen-bond with E312 and R189 and those that form hydrophobic interaction with F135, leading to 9,028 compounds. These filtered compounds were then clustered by their LUNA interaction fingerprints using the Butina algorithm (Tanimoto similarity ≥ 0.75), and cluster representatives were visually inspected. Criteria for prioritizing compounds were followed as previously. 124 compounds were selected, and 108 compounds were synthesized and tested. Because the same selection criteria were applied, 17 of the 108 compounds tested in the MINT-Dock campaign were also tested in the CombiDOCK campaign: Z1542734894, Z2481800190, Z8353197938, Z3360769253, Z9329608260, Z9327964514, Z5313551878, Z2839865511, Z3503238340, Z3367013938, Z1839981454, Z4275682064, Z1438338004, Z9332647693, Z9329608325, Z3294417719, Z1272055642.

For selectivity MINT-Dock screening, we filtered out compounds with undesirable charges and grouped the compounds by SMILES and selected the molecule with the largest docking score difference between VAChT and σ_2_ for each SMILES before applying another score filter of VAChT docking score < -45 kcal/mol and σ_2_ docking score > -40 kcal/mol, leading to 121,628 molecules. For each of these compounds, for pose against each target, we applied strain filtering (111,686 VAChT poses, 110,646 σ_2_ poses) and then novelty filtering (111,516 VAChT poses, 107,673 σ_2_ poses) as described for σ_2_ screening. For interaction filtering using LUNA, we filtered on compounds that form a salt-bridge/hydrogen-bond with the D398 residue of VAChT and do not form an interaction with the D29 residue of σ_2_, leading to 40,449 compounds in total. These filtered compounds were then clustered by their LUNA interaction fingerprints from their docked poses against VAChT using the Butina algorithm (Tanimoto similarity ≥ 0.75), and the VAChT docked poses of cluster representatives were visually inspected. Criteria for prioritizing compounds were followed as previously. 129 compounds were selected, and 114 compounds were synthesized and tested.

For poly-pharmacology MINT-Dock screening, we filtered out compounds with undesirable charges and grouped the compounds by SMILES and selected the molecule with the most favorable sum of docking score for VAChT and docking score for σ_2_ for each SMILES before applying another score filter of VAChT docking score < -45 kcal/mol and σ_2_ docking score < -45 kcal/mol. For each of these compounds, for pose against each target, we applied strain filtering (111,686 VAChT poses, 110,646 σ_2_ poses) and then novelty filtering (111,516 VAChT poses, 107,673 σ_2_ poses) as described for σ_2_ screening, leading to 260,576 compounds. For interaction filtering using LUNA, we filtered on compounds that both form a salt-bridge/hydrogen-bond with the D398 residue of VAChT and form a salt-bridge/hydrogen-bond with the D29 residue of σ_2_, leading to 80,555 compounds in total. These filtered compounds were then clustered by their LUNA interaction fingerprints from their docked poses against VAChT using the Butina algorithm (Tanimoto similarity ≥ 0.75), and cluster representatives were visually inspected. Compounds with poses satisfying criteria for both the VAChT pocket and the σ_2_ pocket were prioritized. 58 compounds were selected, and 53 compounds were synthesized and tested.

### Benchmarking CombiDOCK and MINT-Dock against the fragment-enrichment-based approach

To evaluate recall performance of CombiDOCK to V-SYNTHES at large-scale, we curated a benchmark library of ten million molecules generated from 5,000 randomly selected Syn1s and 2,000 Syn2s within a single amide-forming reaction type. Using the top 5000 scoring compounds (top 0.05% hits) from full-molecule docking as a reference, we assessed the recall performance of CombiDOCK and V-SYNTHES ^10^ across top ranking thresholds of 0.1%, 0.2%, 0.5%, 1%, 2% and 5% of the ten-million library for the σ_2_ receptor, VMAT2 and VAChT. As shown in **Fig. S15A**, CombiDOCK reliably reproduced the top-ranking molecules from full-molecule docking, with recall values for the top 5,000 compounds reaching 45% to 74% at the top 0.2% cutoff of its ranked lists. In contrast, fragment-enrichment approaches, which operate by docking individual synthons to identify promising anchors, enumerating only products containing those anchors, and then docking the reduced product set—achieved recalls of only 4% to 9% using the same docking engine and grids (**Fig. S15A**). Additionally, docking score correlations with full molecule docking remained high across all three targets (σ_2_: Pearson *R* = 0.89, Spearman *R* = 0.88; VMAT2: Pearson/Spearman *R* = 0.67; VAChT: Pearson *R* = 0.69, Spearman *R* = 0.70), underscoring strong rank-order performance.

For MINT-Dock, we compared it to the fragment-enrichment protocol above under matched-scale outputs for both σ_2_ and VMAT2. For a fair comparison of MINT-Dock with the fragment docking idea of enumerating all products available from top scoring synthons, we re-implemented the V-SYNTHES method as described below. After obtaining the synthon DOCK6 scores, we implemented an automatic pose filter that excluded synthon poses that were likely to clash with the protein. The clash was heuristically determined by extending each synthon pose from the centroid of its cap functional group by a 5 Å distance and checking if the extended coordinate was within a 2.8 Å distance with any heavy atom of the target protein. This 2.8 Å clash distance cutoff derived from the clash criterion established for all-atom contact analysis, where the van der Waals shells of two non-bonded heavy overlapped more than 0.4 Å,^140^ was selected as a minimal filter that only excludes synthons when their extension results in a severe steric collision with the target protein. We then selected the top scoring synthons. For both σ_2_ and VMAT2, the number of synthons we chose were selected so that the total number of stereoisomers sampled and the total number of compounds generated from fragment docking and MINT-Dock, where compounds from 10 million rollouts on the two-component space and compounds from 2 million rollouts on the three-component space were combined, respectively were similar. Namely, for σ_2_, we pooled 8.3 million two-component compounds (30 million stereoisomers) from 400 cycles (10 million rollout), and 2.0 million three-component compounds (30 million stereoisomers) from 200 cycles (2 million rollout) generated from MINT-Dock, and we pooled 7.6 million two-component compounds (30 million stereoisomers), and 2.6 million three-component compounds (31 million stereoisomers) generated from fragment docking. For VMAT2, we pooled 9.0 million two-component compounds (30 million stereoisomers) from the first 400 cycles (10 million rollout), and 2.0 million three-component compounds (14 million stereoisomers) generated from MINT-Dock, and we pooled 9.1 million two-component compounds (29 million stereoisomers), and 2.9 million three-component compounds (14 million stereoisomers) generated from fragment docking. Since here the purpose was to compare the two methods to evaluate their performances on both dock scores and interactions with the designated target, we performed the same pipeline on the set of molecules generated by fragment docking as we did on the set of molecules generated by MINT-Dock. For each target, we filtered out compounds with undesirable charges and grouped the compounds by SMILES and selected the molecule with the most favorable adjusted docking score for each SMILES. We then selected the top 300,000 ranked compounds, passed them through strain and novelty filtering, and performed interaction filtering customized for each target. The adjusted enrichment ratio was calculated by the formula presented in comparison with random docking.

For σ_2_ (10.3 million molecules from MINT-Dock and 10.2 million from fragment-enrichment docking), fragment-enrichment produced slightly better raw score distributions directly from docking (**Fig. S15B**). However, when we applied interaction-based triage designed to retain molecules consistent with key interactions critical for σ_2_ binding, MINT-Dock became increasingly favored. In particular, after filtering for a salt bridge with D29, MINT-Dock outperformed fragment-enrichment at most score thresholds, and the advantage widened further when we imposed a stricter criterion requiring an additional hydrogen-bond interaction (**Fig. S15B**). Composition analysis indicated that fragment-enrichment disproportionately selected high-scoring three-component products, which were less likely to satisfy these interaction filters than two-component products, explaining the reversal after interaction-based triage (**Fig. S16**). For VMAT2 (11.0 million molecules from MINT-Dock and 11.9 million from fragment-enrichment docking), MINT-Dock produced more well-scoring molecules than fragment-enrichment docking across score thresholds, both directly from docking and after applying filters tied to key VMAT2 interactions, including E312 and R189 (**Fig. S15C**).

Given CombiDOCK’s higher recall and closer agreement with full docking in the retrospective benchmarking analysis, we next evaluated how a fragment-enrichment-based virtual screening method performs in recovering true actives from our prospective CombiDOCK screens against σ_2_, VMAT2, and VAChT. We used the 103B Enamine REAL library and performed a similar V-SYNTHES-style full molecule enumeration and assessed what percentages of tested actives were included when top 0.02%, 0.05%, 0.1%, 0.5%, 1%, 2%, 5%, and 10% of the top ranked synthons were enumerated.

**Fig. S15D** summarizes recall at progressively higher selection thresholds for enumerated synthon libraries, where recall values represent the proportion of true actives present in the final library. The fragment-enrichment-based method performed similarly across the three targets, with poor recovery of actives at early stages of enrichment (top ≤ 0.1%), with a maximum recall rate of 8% against VAChT if the top 0.1% are enumerated. Recall improved monotonically with broader selections, surpassing 90% by the top 5%. Notably, for VAChT the most potent CombiDOCK hit (CSMS00228273820; IC50 = 12 nM) was not recovered until enumeration of the top 0.5% of synthons, and similarly, for σ_2_ and VMAT2 the most potent actives only appeared when the top 5-10% of synthons were selected, requiring enumeration and full docking of more than 500 million molecules (**Table S5**).

With the fragment approach, achieving complete recall of all 64 VAChT actives required enumerating and generating 3D conformers for > 1 billion full molecules. End-to-end, fragment-docking, full-molecule enumeration, followed by full-molecule docking, this takes 179 cluster-days on 2,000 CPUs. In contrast CombiDOCK docks synthons and assembles them into full molecules in a single pass, completing in 26 cluster-days. MINT-Dock achieves even greater efficiency by concentrating docking on the most productive regions of chemical space, typically completing a campaign in 3 to 4 days on approximately 100 CPU cores while generating molecules that better satisfy target-specific interaction criteria than fragment-enrichment at matched scale. Together, these comparisons establish that both CombiDOCK and MINT-Dock overcome the core limitation of fragment-enrichment approaches.

### DrugCLIP comparison

The recently published DrugCLIP is able to score over trillions of ligand-protein interactions per cluster day using 8 GPUs, which has enabled genome-wide docking, discovering potent binders prospectively for three biological targets.^6^ DrugCLIP reports that screening 500 million compounds against 10,000 proteins takes 1 cluster-day on 8 GPUs. However, under DrugCLIP’s stated computational complexity O(M+N) (with M targets and N compounds), scaling from 0.5 billion to 100 billion compounds implies 200 cluster-days on the same 8-GPU node, largely independent of whether M is 10,000 proteins or a single protein (because runtime is dominated by N). In addition, DrugCLIP’s ligand encoder (Uni-Mol^141^) requires precomputed 3D conformers generated using RDKit, and the paper describes a data augmentation strategy that can generate up to 10 conformations per molecule. Even assuming a conservative one conformer per molecule for library embedding, conformer generation becomes a major overhead at the 100-billion scale. For example, with 2,000 CPU cores and 0.2 s per conformer per CPU-core, generating one conformer per molecule for 100B molecules requires 116 cluster-days. Taken together, even under optimistic assumptions, the end-to-end runtime for the DrugCLIP retrieval step at 100 billion molecules on a single target is at least 316 cluster-days. In contrast, CombiDOCK requires only 1 cluster-day to prepare the synthon library and 30 cluster-days to screen 100 billion molecules against a single target, implying roughly a 10-fold faster end-to-end workflow at this scale. Moreover, DrugCLIP is fundamentally a retrieval approach: to make structure-based predictions, it must be paired with additional docking engines, which contributes to end-to-end wall time beyond the retrieval stage. On our targets, DrugCLIP predictions do not even recover the experimental ligand binding site. For σ_2_, DrugCLIP yields no predictions in [their public database](https://drug-the-whole-genome.yanyanlan.com/), which is surprising given σ_2_ is considered ligandable. For both VMAT2 and VAChT, visual inspection of the reported DrugCLIP pockets/hits indicates all predicted binding sites that are spatially distant from the canonical ligand site (**Fig. S11C-D**). We therefore re-docked the top DrugCLIP hits using the same docking engine used in this study and found they score substantially worse than CombiDOCK’s top-ranked compounds on both VMAT2 and VAChT (**Fig. S11C-D**).

### Analoging in the make-on-demand library

The most potent compound identified in our 103 billion CombiDOCK screen against VAchT was CSMS00228273820 with an IC_50_ = 12 nM. The most potent compound identified in our MINT-Dock screen for selectivity was Z9424360192 and Z2406174633 with an IC_50_ = 86 nM and 100 nM respectively. To optimize the potency of the three compounds we curated a set of close structural analogues from Enamine REAL Space make-on-demand library for testing. Using database search tool swp.docking.org,^142^ we queried 68 compounds with a maximum graph edit distance of 12 relative to the structure of CSMS00228273820, 1176 compounds with a maximum graph edit distance of 6 relative to the structure of Z9424360192 and 1128 compounds with a maximum graph edit distance of 7 relative to the structure of Z2406174633. Stereoisomers and protomers were enumerated for all compounds followed by 3D conformer generation. Conformers for all compounds were stored in DB2 format and were docked to both our VAChT and σ₂ models using DOCK6 HDB. Here we placed the primary focus on structural similarity of analogs with our parent compounds, with σ₂ selectivity as a secondary filter. For CSMS00228273820, we selected 21 analogues for synthesis, all of which were successfully synthesized and experimentally tested. For Z9424360192, we selected 22 analogs for synthesis, 20 were successfully synthesized and experimentally tested. For Z2406174633, we selected 32 analogs for synthesis, 29 were successfully synthesized and experimentally tested. Of these 29 analogs, Z2406174639, Z2406174433, and Z2405924153 were also analogs of CSMS00228273820.

### Expression and purification of VAChT

The coding sequence corresponding to human VAChT (residues 34 to 524) was subcloned into the pBMCL1 vector.^143^ The DNA sequence encoding MBP and the helical linker with the sequence (AEEEKRK)_2_ were incorporated at the N terminus of VAChT. Meanwhile, the flexible linker (GGGGS)_2_ was fused to the C-terminus of VAChT, together with the coding sequence for DARPinoff_7_^144^ (TransGen Biotech). This fully assembled construct was referred to as the VAChT^EM^ hereafter.

VAChT^EM^ was expressed in HEK293S GnTI− cells utilizing the BacMam system.^145^ Baculoviruses were generated through transfection of Sf9 cells. Following three successive rounds of amplification, the viruses were used for cell transduction. HEK293S cells, maintained at a density of 3×106 cells/ml, were subjected to infection via the addition of 10% third-passage baculoviruses, and subsequently cultured at 37°C for 8–12 h. Thereafter, 10 mM sodium butyrate was introduced to trigger protein expression, after which the cultures were further incubated at 30°C for an additional 48 h. The cells were harvested via centrifugation at 4,000 rpm for 20 min and then stored at −80°C.

To purify VAChT^EM^, cell pellets were resuspended in lysis buffer (50 mM HEPES pH 7.25, 300 mM NaCl, 15% glycerol) supplemented with 2 μg/ml DNase I and protease inhibitor cocktail (APExBIO). Cell lysis was then performed at 4°C for 3 h using 2% Lauryl maltose neopentyl glycol (LMNG), 0.2% Cholesteryl hemisuccinate (CHS), and 0.67% Glyco-dendrimer (GDN). Following centrifugation at 18,000 rpm for 40 min, the supernatant was subjected to incubation with anti-GFP nanobody affinity resin at 4°C for 2 h. The resin was subsequently washed using 40 column volumes of Buffer A (25 mM HEPES pH 7.25, 150 mM NaCl, 0.003% LMNG, 0.001% GDN, 0.0003% CHS). To remove the GFP tag and release VAChT^EM^ from the resin, GST-tagged PreScission Protease was utilized. The protease was thereafter removed via adsorption to Glutathione Beads (Smart-Lifesciences). The protein underwent further purification via size-exclusion chromatography (SEC), using a Superose 6 Increase 10/300 GL column (GE Healthcare) in Buffer A. Peak fraction containing VAChTEM was concentrated to a final concentration of ∼8 mg/ml using a 50 kDa cut-off centrifugal filter (Millipore). The protein sample was stored at −80°C until subsequent use.

### Cryo-EM sample preparation and data collection

The protein was pre-incubated with 600 μM CSM00228273820 or Z9424360192 at 4°C for 20 min before initiating cryo-EM sample preparation. A 3 μL aliquot of the protein sample was applied onto glow-discharged holey carbon grids (Quantifoil R0.6/1.0, Au300 mesh). Using a Vitrobot Mark IV (FEI), the grids were flash-frozen in liquid ethane cooled via liquid nitrogen under conditions of 8°C and 100% humidity. For the blotting step, the operational parameters were set as follows: a blot time of 3 s and a wait time of 10 s.

The grids were screened utilizing a 200 kV Talos Arctica microscope (FEI), which was outfitted with a Gatan K2 Summit detector. Acquisition of raw movie stacks was carried out with a 300 kV Titan Krios microscope (FEI) paired with a Gatan K3 camera. The data were recorded at a physical pixel dimension of 0.83 Å per pixel and a nominal defocus interval ranging from 1 to 2 μm, with each individual movie comprising 40 frames. The total exposure dose was around 60 e^−^/Å^2^. Ultimately, 9,261 and 6,847 micrographs were acquired in total for CSM00228273820-bound VAChT and Z9424360192-bound VAChT respectively. Comprehensive details of all data collection parameters are compiled in **Table S4**.

### Cryo-EM image processing

All data processing procedures were carried out solely within cryoSPARC.^146^ The image stacks were gain-normalized and corrected for beam-induced motion using Patch Motion Correction. Estimation of CTF parameters was accomplished through the Patch CTF Estimation tool. Micrographs that were unsuitable for subsequent analysis were excluded via manual inspection. Blob picking was utilized for initial particle picking and the generation of 2D templates. Following three cycles of 2D classification, particles of high quality were used for ab initio reconstruction and heterogeneous refinement processes. Low-pass filtered maps obtained from this stage guided further heterogeneous refinement, which was subsequently followed by non-uniform refinement^147^ of particles derived from the optimal classes. These refined particles served as reference templates for template-based picking or as training datasets for the Topaz particle-picking pipeline.^148^ Subsequent iterations of ab initio reconstruction, heterogeneous refinement, and additional heterogeneous refinement steps, guided by low-pass filtered maps from the prior blob picking step, were repeated to merge the optimal particle classes. Following the removal of duplicate particles, the resulting particle subsets underwent refinement via both non-uniform refinement and local refinement. Subsequently, the refined map and particles were subjected to seed-facilitated classification and refinement,^149^ with this process conducted against the full dataset of initially picked particles. A final round of local refinement, focused on the transporter, yielded a 3.6 Å-resolution map for the CSM00228273820-bound state and a 3.7 Å-resolution map for the Z9424360192-bound state (gold-standard FSC = 0.143).

### Model building and refinement

The vesamicol-bound VAChT structure (PDB: 8ZMR)^101^ was first docked into the cryo-EM density map using ChimeraX.^150^ Subsequently, real-space refinement was executed via PHENIX,^151^ while manual adjustments were made within Coot^152^ Iterative cycles of automatic and manual refinement were conducted until the final structural model was achieved. Estimation of the local resolution for the cryo-EM map was carried out using cryoSPARC.^146^ The geometric integrity of the structural model was validated through MolProbity.^140^ All structure-associated figures were generated utilizing ChimeraX.

### σ_2_ radioligand binding assay

Membranes for radioligand binding assays were prepared from Expi293 cells (Thermo Fisher Scientific) transfected with wild-type σ_2_ receptor using FectoPRO (Polyplus-transfection) according to the manufacturer’s instructions. Cells were harvested 72 hours post-transfection and pelleted by centrifugation. Cell pellets were resuspended in 20 mM HEPES (pH 7.5), 2 mM magnesium chloride, and 1:100,000 (vol:vol) benzonase nuclease (Sigma Aldrich), supplemented with cOmplete Mini EDTA-free Protease Inhibitor Mixture Tablets (Roche). Following Dounce homogenization, cells were centrifuged at 50,000 × g for 20 min at 4°C. The resulting membrane pellet was resuspended in 50 mM Tris (pH 8.0), subjected to final Dounce homogenization, aliquoted, flash frozen, and stored at −80°C until use.

For radioligand binding experiments membranes were thawed, homogenized, and incubated with 5 nM [^3^H]-DTG (PerkinElmer) in 100 μL binding reactions containing 50 mM Tris (pH 8.0), 0.1% (w/v) bovine serum albumin, and 50 nM PD-144418 to block σ_1_ receptor binding sites. For competition assays, competing ligands were added at 1 μM for single-point measurements. Reactions were incubated at 37°C for 2 hours with gentle agitation. Reactions were terminated by rapid filtration through glass fiber filters pre-treated with 0.3% polyethylenimine using a Brandel cell harvester. Filters were washed with ice-cold water, then individually soaked in 5 mL Cytoscint scintillation fluid (MP Biomedicals) overnight. Radioactivity was quantified using a Beckman Coulter LS6500 scintillation counter. All reactions were performed in triplicate using 96-well format.

### VMAT2 radioligand uptake assay

The uptake assays were performed as previously described.^98^ Human embryonic kidney (HEK293) cells (ATCC, CRL-3216) were transiently transfected with a VMAT2-encoding plasmid using Lipofectamine 3000 according to the manufacturer’s protocol. Cells were cultured at 37 °C for 16–18 h, supplemented with 10 mM sodium butyrate, and further incubated at 30 °C for an additional 24 h.

Assays were performed 48 h after transfection. The medium was aspirated, and cells were washed with ice-cold assay buffer (110 mM potassium gluconate, 5 mM glucose, 5 mM MgCl2, 1 mM ascorbic acid, 30 μM pargyline and 20 mM HEPES pH 7.4), then incubated with 10 μM digitonin in assay buffer for 5 min at room temperature and washed again with ice-cold assay buffer without digitonin. Permeabilized cells were incubated with an assay buffer containing 5 mM ATP with test compounds at a final concentration of 10 μM. After 10 min incubation at room temperature, 50 nM [^3^H]-dopamine (ViTrax Radiochemicals) was added, cells were subsequently incubated at room temperature for 20 min. The substrate–containing buffer was aspirated, cells were washed twice with ice-cold assay buffer and then solubilized in P2 lysis buffer (Qiagen). Scintillation liquid (Ultima Gold) was added to cell lysates to measure radioactivity using a Microbeta counter (PerkinElmer).

### VAChT radioligand binding assay

Radioligand binding experiments were performed utilizing crude cellular membranes harboring human VAChT. HEK293F cells that had been transiently transfected with full-length VAChT plasmid were cultured at 37°C for 16−18 h, after which 10 mM sodium butyrate was added and the cells were further incubated at 30°C for additional 24 h. Following cell collection, the cells were washed once with cold PBS and then resuspended in ice-cold SH buffer (10 mM HEPES pH 7.4, 320 mM sucrose, protease inhibitor cocktail) at a density of approximately 4×106 cells/mL. The resuspended cells were lysed via sonication, then centrifuged at 4,000 g for 5 min to eliminate unlysed cells and cellular debris. The resulting supernatant was then flash-frozen in liquid nitrogen and stored at −80°C for subsequent use.

In the [3H]-vesamicol competition binding assay, 50 μL aliquots of membrane suspensions were subjected to a 60-min incubation at 25°C with 150 μL of binding buffer (20 mM HEPES pH 7.4, 110 mM potassium tartrate, 1 mM ascorbic acid) containing 3 nM (−)-[3H]-vesamicol (30–60 Ci/mmol, American Radiolabeled Chemicals, Inc.) and 7.5 μM of the test compounds. Reactions were terminated via rapid vacuum filtration through GF/C glass fiber filters (Whatman) pretreated with 0.3% polyethyleneimine, with subsequent washing using 2 ml of ice-cold binding buffer. The radioactivity retained on the filters was quantified by scintillation counting in 2.5 mL of scintillation solution (PerkinElmer).

### Dose response curve

For σ_2_ top compounds measurement, the dose-response assays were performed in the same manner as the competition binding assays, except that test compounds were added at a range of concentrations to generate full displacement curves. Data were analyzed using GraphPad Prism 10 (http://www.graphpad.com). Kᵢ values were calculated using the Cheng–Prusoff equation as implemented in the software.

For VMAT2 top compounds measurement, the dose-response assays were performed similarly to the [^3^H]-dopamine uptake assay, except that the permeabilized cells were incubated with assay buffer containing 5 mM ATP with a series of specified concentrations of the test compounds. IC_50_ values were determined by fitting three-parameter dose–response curves using GraphPad Prism 10.

For VAChT top compounds measurement, the dose-response assays were conducted in a similar manner to the competition binding assay, with the exception that the membrane suspension was incubated with a series of specified concentrations of the test compounds. The IC_50_ values were determined by fitting logIC_50_ curves using GraphPad Prism 10.

### Microscale Thermophoresis (MST) analysis

The binding affinity between VAChT (and its mutants) and CSMS00228273820 was measured by MST experiments using a NanoTemper Monolith NT.115 instrument (NanoTemper Technologies). VAChT and mutants were first fluorescently labeled using the Red-NHS 2^nd^ Generation kit (NanoTemper Technologies) in the assay buffer (25 mM HEPES pH 7.4, 150 mM NaCl, 0.02% DDM and 0.002% CHS). For the binding assay, 50 nM of the labeled VAChT was then incubated with CSMS00228273820 at 16 serially diluted concentrations, ranging from 0.476 nM to 15.6 μM, prepared in the same assay buffer. After a 20-min incubation at 16°C, the mixture was then loaded into Standard Monolith Capillaries (NanoTemper Technologies). MST measurements were conducted using an excitation power of 20% and an MST power of 40%. Capillaries exhibiting aggregation or adsorption were excluded. The *K*_d_ value was calculated using the MO.Affinity Analysis software (NanoTemper Technologies).

### Make-on-demand synthesis

All the make-on-demand molecules were derived from Enamine REAL (https://enamine.net/compound-collections/real-compounds). See supplementary materials for the synthesis procedures and characterization of compounds.

### Bemis-Murcko scaffold analysis

Molecular scaffolds were generated using RDKit,^128^ following the Bemis–Murcko framework,^116^ to enable structural comparisons among compounds from different library screens.

### MINT-Dock comparison with large-scale docking

The large-scale docking campaigns for 3 targets were described previously. We added another σ_2_ docking campaign of 1.73 billion stereogenic molecules, which is a strict subset of the 1.85 billion molecules in prior VMAT2 and VAChT screening campaigns, for both an additional comparison with MINT-Dock and comparison with multi-target MINT-Dock. The less 0.12 billion molecules were not procured due to a reorganization of the ZINC22 database. For rank comparison, we coalesced the stereoisomers and assigned the best DOCK score for each unique ZINC ID from the large-scale docking campaign. For single-target comparison, we used the formula 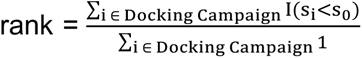 that is the fraction of docked molecules for each docking campaign with scores below a threshold s_0_ to represent its efficiency. The higher the rank is, the more efficient the campaign is. For VMAT2 comparison, we applied a post-docking strain filter of total torsion strain energy of less than 8 units, maximum strain energy per torsion angle of less than 1.5 units to molecule poses from the 1.85 billion large-scale docking campaign in order to match with the MINT-Dock docking setup. The 660-fold enrichment for σ_2_ was determined from using the score threshold of -56.5 kcal/mol, and the 130-fold enrichment for VMAT2 was determined from using the score threshold of -53.5 kcal/mol, same as in the “**MINT-Dock Backpropagation Comparison**” section. For rank comparison for selectivity screens, we used the formula 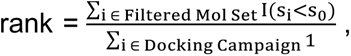 where “Filtered Mol Set” was defined to be the set of molecules with VAChT docking score below -45 kcal/mol and σ_2_ docking score above -40 kcal/mol. For rank comparison for poly-pharmacology screens, we used the formula 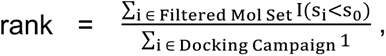 where “Filtered Mol Set” was defined to be the set of molecules with VAChT docking score below -45 kcal/mol and σ_2_ docking score below -45 kcal/mol.

### Technical details for CombiDOCK

CombiDOCK was implemented in the latest release of DOCK6 (version 6.14). It was built upon a hierarchical database (HDB) search routine, which traverses precomputed ligand conformations stored in a Dock Binary version 2 (DB2) format, in a similar way to that in DOCK3.^50,127^ Our method is based on HDB docking framework re-implemented in DOCK6^84^ but extends them by enabling hierarchical tree sampling of two molecular fragments in a sequential and combinatorial manner. Below are the technical details:

#### 1. DB2 file format in hierarchical organization

The DB2 file format stores ligand conformations in a highly compressed and hierarchical structure. In this format, each molecule is associated with one or more hierarchies, each built around a rigid segment, which serves as a reference frame for all conformational variations. For example, a molecule containing multiple rings will have each ring treated as an independent rigid segment, and a separate hierarchy will be generated for each. Thus, a molecule with three rings will have three corresponding hierarchies.

Each hierarchy is organized as a tree structure, where the root node represents the rigid segment, and subsequent nodes represent molecular segments in specific conformations. A complete branch of the tree corresponds to a fully grown molecular conformation. Since the rigid root segment is shared among all branches, it provides a fixed reference for the relative positions of different branches.

The standard DB2 file^127^ is composed of several sections:

- M (Header): General molecule information
- A (Atoms): Atom definitions
- X (Coordinates): Cartesian coordinates of atoms
- C (Conformers): Molecular segment information
- S (Sets): Branch or conformer information

This hierarchical organization enables efficient storage and rapid conformational search, making the DB2 format well-suited for ultra-large library docking. To support combinatorial docking of two synthons, CombiDOCK extends the DB2 format with an additional overlay section:

- **O (Overlay)**: Overlay segment information

The overlay section contains the overlay segment information, which is derived from its capping group. This information is used to align and fuse two separate fragments into a complete molecule during docking. By including an overlay section within the DB2 structure, CombiDOCK maintains the efficiency of hierarchical sampling and full molecule reconstruction.

Moreover, because each synthon can serve as either the first (Syn1) or the second (Syn2) fragment in the docking process, we prepared two separate DB2 files for each: In the Syn1 form, the number of ring systems determines the number of hierarchies. Each ring is treated as a rigid segment and used as the root of a hierarchy. In the Syn2 form, only a single hierarchy is generated, with the capping group defined as the rigid segment. This dual representation ensures that each synthon can be appropriately used for both stages of combinatorial docking.

#### 2. DB2 building workflow

The DB2 building workflow converts 2D molecular representations (full molecules or synthons) into 3D conformer ensembles stored in the hierarchical DB2 format, enabling efficient sampling during docking. The process involves six key stages: protonation and tautomerization state enumeration, stereoisomer enumeration, 3D conformer generation, partial charge and desolvation calculation, conformational expansion, and hierarchical data structuring.

- **Protonation and tautomerization state enumeration** Each input molecule (represented using SMILES string) is first processed using the Jchem suite (v.22.22.0, https://chemaxon.com/). The cxcalc and molconvert utilities are used to enumerate protonation and tautomerization states at physiological pH (7.4). Each unique protomer or tautomer is treated as an independent molecular entry for downstream processing.
- **Stereoisomer enumeration** For each protomer/tautomer, RDKit (v2023.09.1)^128^ is used to enumerate all possible stereoisomers based on undefined chiral centers. Each stereoisomer is retained as an independent molecular variant for downstream 3D conformer generation.
- **3D conformation generation** Each stereoisomer is converted into a 3D geometry using Corina Classic (v.4.4.0026, https://mn-am.com/products/corina/), which generates a single low-energy conformation while preserving stereochemistry. This initial geometry serves as the input for subsequent partial charge and solvation calculations.
- **Partial charge and desolvation calculations** AMSOL (version 7.1) is used to compute AM1-BCC partial atomic charges^153,154^ and per-atom desolvation energy decompositions using a Generalized Born (GB) solvation model. These charge and solvation descriptors are stored with the molecular data (mol2 format) and propagated into the DB2 structure for electrostatic and ligand solvation terms.
- **Conformational expansion** Using the Corina-generated 3D geometry as a reference, OMEGA (v.2019.10.2, https://www.eyesopen.com/omega)^129^ is employed to generate a hierarchical ensemble of conformers by systematically sampling torsional degrees of freedom. The resulting conformers are filtered by energy and RMSD thresholds (energy window = 12, RMSD cutoff = 0.5 Å) to remove redundant or high-energy conformations, yielding a compact yet representative conformational set.
- **Hierarchy construction and DB2 generation** The OMEGA-generated conformers are organized hierarchically according to their rigid segments using the mol2db2 utility from the DOCK 3.7 (https://dock.compbio.ucsf.edu/DOCK3.7/). Each rigid segment serves as the root of a hierarchy, while flexible torsions define successive branches.

- For multi-ring full molecules, each ring is treated as a separate rigid root, resulting in multiple hierarchies.
- For synthons, two DB2 representations are generated:

- Syn1 form – multiple hierarchies, one per ring system.
- Syn2 form – a single hierarchy rooted at the capping group. This dual representation allows each synthon to function as either the first (Syn1) or second (Syn2) fragment in the combinatorial docking process.

Hierarchical data are converted into DB2 format using the build_ligands.py utility from the DOCK 3.7. Each DB2 file contains five primary sections – M (Header), A (Atoms), X (Coordinates), C (Conformers), and S (Sets) – and, for synthons, an additional O (Overlay) section encoding atom indices for fragment fusion during docking. We modified build_ligands.py to incorporate RDKit for reading 3D molecular structures (mol2 files) and automatically detecting atom indices corresponding to the capping group, which enables accurate Overlay information for combinatorial docking.

#### 3. HDB search routine

The HDB search begins by aligning the rigid segment of the ligand to the receptor matching spheres. If the alignment is incompatible (e.g., score is not acceptable due to the steric clashes), the match is discarded, and the next alignment is considered.

For each conformational set (or branch set), segment conformations are evaluated sequentially. Each segment is first oriented based on the rigid segment position and then scored. If a segment is found to be incompatible (e.g., score is not acceptable due to the steric clashes), it is flagged. This prevents redundant evaluation if the same segment is encountered in other branches. Once an incompatibility is detected, the current search along these sub-branches is terminated, and the routine proceeds to the next branch. If all segment conformations in a branch are compatible, the entire branch is retained as a valid conformation. After all the branches are traversed, each valid conformation is minimized through rigid-body minimization to further refine the pose. The final refined poses are ranked by docking scores and the top-scoring pose is retained for virtual screening (VS).

#### We implemented CombiDOCK HDB search routine by enabling hierarchical search of two molecular fragments in a sequential and combinatorial manner

Same as in the standard HDB search, the rigid segment of the first synthon (Syn1) is oriented to the receptor matching spheres, and segment conformations are evaluated sequentially along each branch. Incompatibilities of the segment conformation terminate the evaluation of that branch, and the routine continues to the next until all branches are traversed. Valid Syn1 conformations are saved.

Next, for each valid Syn1 conformation, its capping group is used as the reference frame to orient the rigid segment of the second synthon (Syn2). The rigid segment of Syn2 is aligned by superposition of its capping group onto that of Syn1 using the Kabsch algorithm. After alignment, the search proceeds using the same segment-by-segment evaluation described above. All valid Syn2 conformations are stored for each Syn1 pose, and the resulting combinations are fused into complete molecules. These full-molecule poses are refined through rigid-body minimization and then ranked by docking score. The top-scoring pose is retained for VS.

#### 4. Fuse Syn1-Syn2 into full molecule

Each full molecule is constructed by fusing a pair of synthons (Syn1 and Syn2) using predefined overlay information extracted from DB2 files. This overlay section specifies the atom indices to be deleted from Syn1 and Syn2, as well as the atom indices from each synthon to be bonded to form the final molecule.

During the construction of the full molecule, atomic partial charges are rescaled using a sign-correcting approach. For example, if the computed total charge of the resulting molecule is +0.2 but the target total charge should be 0, the charge corrector is -0.2 (i.e., 0 – 0.2 = -0.2). Since this corrector is a negative value, only the negatively charged atoms are uniformly adjusted. Each negative atom receives a per-atom correction equal to the total charge corrector divided by the number of negatively charged atoms. This adjustment slightly increases the magnitude of the negative charges while preserving the overall charge distribution. In parallel, the atomic polar solvation energies are rescaled proportionally based on the ratio of corrected to original partial charges. This ensures consistency in energy values following charge adjustment.

#### 5. Combinatorial docking parameters

The docking parameters used for CombiDOCK are summarized in the table below and parameters specific to the CombiDOCK are highlighted in **Table S6**.

## Data availability

Active molecules reported here are available from J.L. or directly from Enamine (enaminestore.com). The structures of VAChT determined in complex with docking hits have been deposited in the Protein Data Bank under accession codes PDB: XXX (CSMS00228273820-bound) and PDB: XXX (Z9424360192-bound), and the corresponding cryo-EM maps are available from the Electron Microscopy Data Bank under accession codes EMD-XXXXX and EMD-XXXXX, respectively. The compounds docked from the ZINC make-on-demand library are freely available at http://zinc22.docking.org. Docking scores and SMILES for all screened molecules have been deposited on the Lyu Lab (Rockefeller University) Globus Share Collection: CombiDOCK_docking_data, and will be made publicly available upon publication. Any other data relating to this study are available from the corresponding authors on reasonable request. Source data are provided with this paper.

## Code availability

DOCK 3.8 code is freely available on [GitHub](https://github.com/docking-org/dock3). CombiDOCK is implemented in DOCK 6.13. The code will be freely available on [GitHub](https://github.com/Lyu-Lab/CombiDOCK_branch) upon publication. The MINT-Dock code will be available on [GitHub](https://github.com/jxZh0104/MINTDock/) upon publication.

## Acknowledgements

This work was supported by Rockefeller University start-up funds (J.L.), Irma T. Hirschl/Monique Weill-Caulier Trusts (J.L.), and the Stavros Niarchos Foundation (SNF) as part of its grant to the SNF Institute for Global Infectious Disease Research at The Rockefeller University (J.L.). J.L. is supported by the Searle Scholars Program. This research was also funded by the National Natural Science Foundation of China (to Z.Z., NO. 32471252) and the Beijing Natural Science Foundation (to Z.Z., NO. Z240013). The study was also supported in part by the Center for Life Sciences and the Qidong-SLS Innovation Fund (to Z.Z.). We thank the High-Performance Computing Resource Center (HPCRC) at The Rockefeller University (RRID:SCR_025889) for computational resources. We thank Trent Balius for assistance with the DOCK6 source code and helpful discussions; Aakash Davasam for providing AF3 benchmarking data for σ_2_ and SLC transporters; Dr. Jason Banfelder and Dr. Linelle Abueg for testing MINT-Dock code at Rockefeller HPCRC. We thank the Cryo-EM platform at the School of Life Sciences (SLS) of Peking University for helping with cryo-electron microscopy data collection. We thank Dr. Shitang Huang at the National Center for Protein Sciences of Peking University for assistance with the radioactive [^3^H]-vesamicol displacement experiments. We thank Dr. Brian Shoichet, Dr. Tarun Kapoor, Dr. Jue Chen, Dr. Seth Darst, Dr. Trent Balius, Dr. Jie Zhou, Dr. Fangyu Liu, and Dr. Hongyan Du for reading this work and helpful suggestions. We thank ChemAxon for a license to JChem, OpenEye Scientific software for a license to OEChem and Omega2, Molecular Networks for a license to Corina, Schrodinger LLC for the use of the prepwizard program in Maestro, Revvity Signals for the use of ChemDraw. Molecular graphics and analyses were performed with UCSF Chimera and UCSF ChimeraX. We polished and revised the manuscript with ChatGPT-5.3 and ChatGPT-5.4. We thank BioRender.com for graphic elements that were incorporated in our Figure 1B and 1F.

## Author contributions

J.L. conceived the study; C.Y. developed and implemented the CombiDOCK method in DOCK 6 and built the DOCKable files for the Synthon library; J.Z. and J.L. conceptualized the MINT-Dock algorithm; J.Z. developed the MINT-Dock algorithm; J.L. constructed docking grids for VMAT2 and VAChT, and performed subsequent in-stock and ZINC22 docking; C.Y. and B.L. performed CombiDOCK docking, chemoinformatics analyses, and ligand selection, with assistance from J.L.; J.Z. ran computational benchmarking for MINT-Dock with assistance from B.L.; J.Z. ran and analyzed MINT-Dock prospective campaigns with assistance from C.Y., B.L., and J.L.; Y.W. and C.Y. organized the large-scale docking data; Y.Z. and Z.Z. performed primary radioligand displacement assays, dose-response curve measurements, and cryo-EM structure determination for VAChT compounds; C.B. and A.A. performed primary radioligand displacement assays and dose-response curve measurements for compounds from the σ_2_ campaign and VAChT multi-target campaigns; X.C., S.P., and C.L. performed primary radioligand displacement assays and dose-response curve measurements for VMAT2 compounds; Y.S.M. and D.S.R. directed compound synthesis, purification, and characterization; J.L., Z.Z., C.L., and A.A. supervised the project; J.L., C.Y., J.Z., and B.L. wrote the manuscript with input from all authors.

