## Supplemental materials for "Combinatorial docking and molecular generation to navigate over 100-billion molecules for prospective ligand discovery"

### Supplemental information

**The PDF file includes:**

Supplemental figures S1 to S18
Supplemental tables S1 to S9

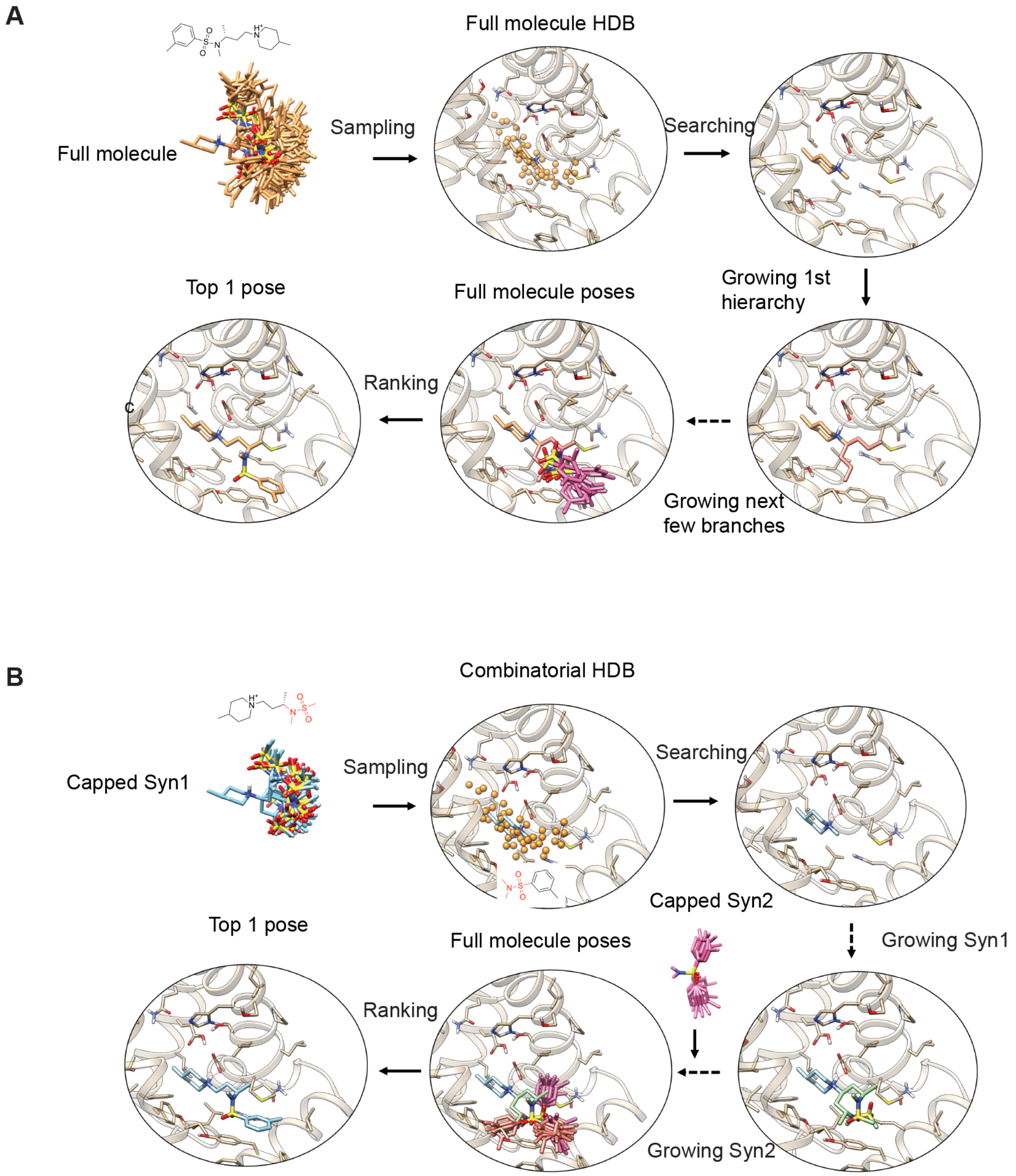


**Figure S1. Contrasting CombiDOCK with full-molecule docking**. (A) Traditional hierarchical database (HDB) docking in DOCK3 and its DOCK6 re-implementation operates on a hierarchical tree of precomputed conformers for each full molecule. (B) By contrast, CombiDOCK constructs two hierarchical trees corresponding to two molecular fragments (Syn1 and Syn2), which are then recombined during docking to reconstruct the full molecule.

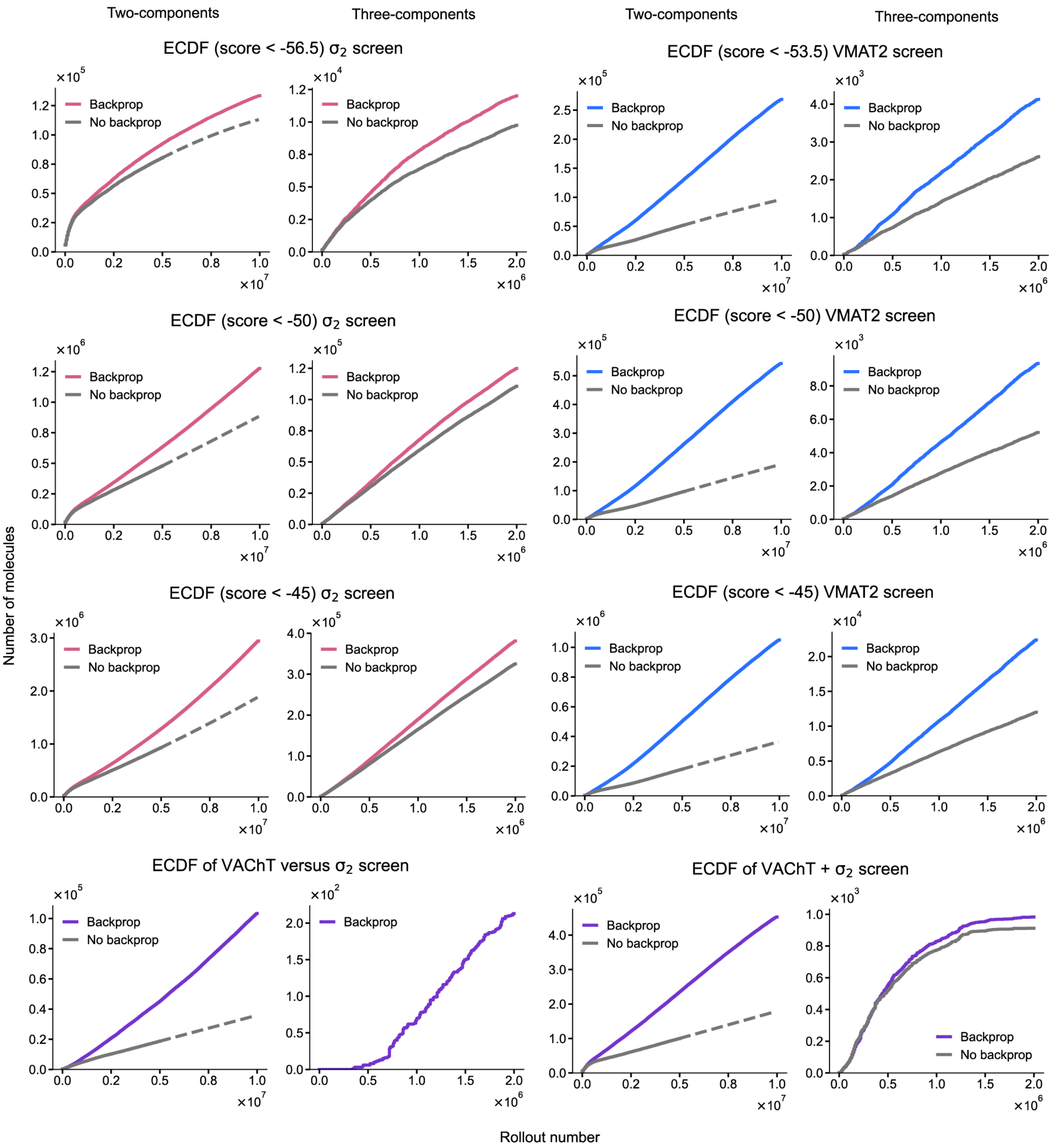


**Figure S2. Evaluation of backpropagation performance.** Comparison of the empirical cumulative distribution function (ECDF) of generated good compounds against $\sigma_2$, VMAT2 (docking score below a given score threshold), VAChT with selectivity constraints (VAChT docking score below -45, $\sigma_2$ docking score above -40), or VAChT with polypharmacology constraints (both targets docking score below -45) with or without backpropagation. Dashed line indicates projected trendlines by a polynomial extrapolation function. X-axis is the rollout number, which equals to the cycle count times the theoretical maximum number of unique synthon combinations that could be enumerated per cycle.

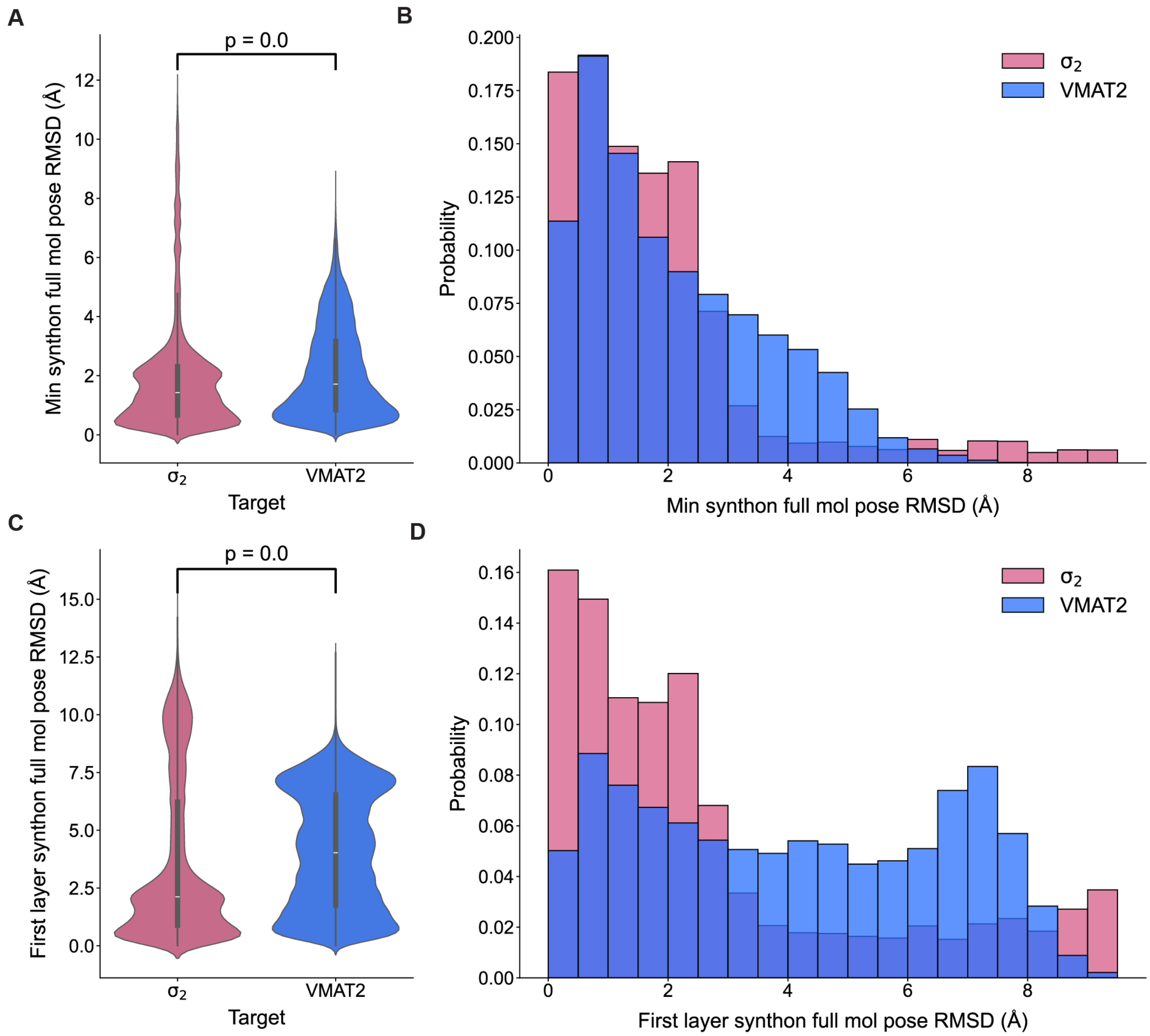


**Figure S3. Comparison of RMSD between synthon poses and full molecule poses across the top 300,000 ranked compounds.** (A) Comparison of the distribution of the minimum of the RMSD between $synthon_1$ and full molecule and the RMSD between $synthon_2$ and full molecule for each target. P-value was calculated by the two-sided Mann-Whitney U test with a significance level of 0.05. (B) Comparison of the density distribution of the minimum of the RMSD between synthon1 and full molecule and the RMSD between synthon2 and full molecule for each target. (C) Comparison of the distribution of the RMSD between the first layer synthon and full molecule for each target. P-value was calculated by the two-sided Mann-Whitney U test with a significance level of 0.05. (D) Comparison of the density distribution of the RMSD between the first layer synthon and full molecule for each target.

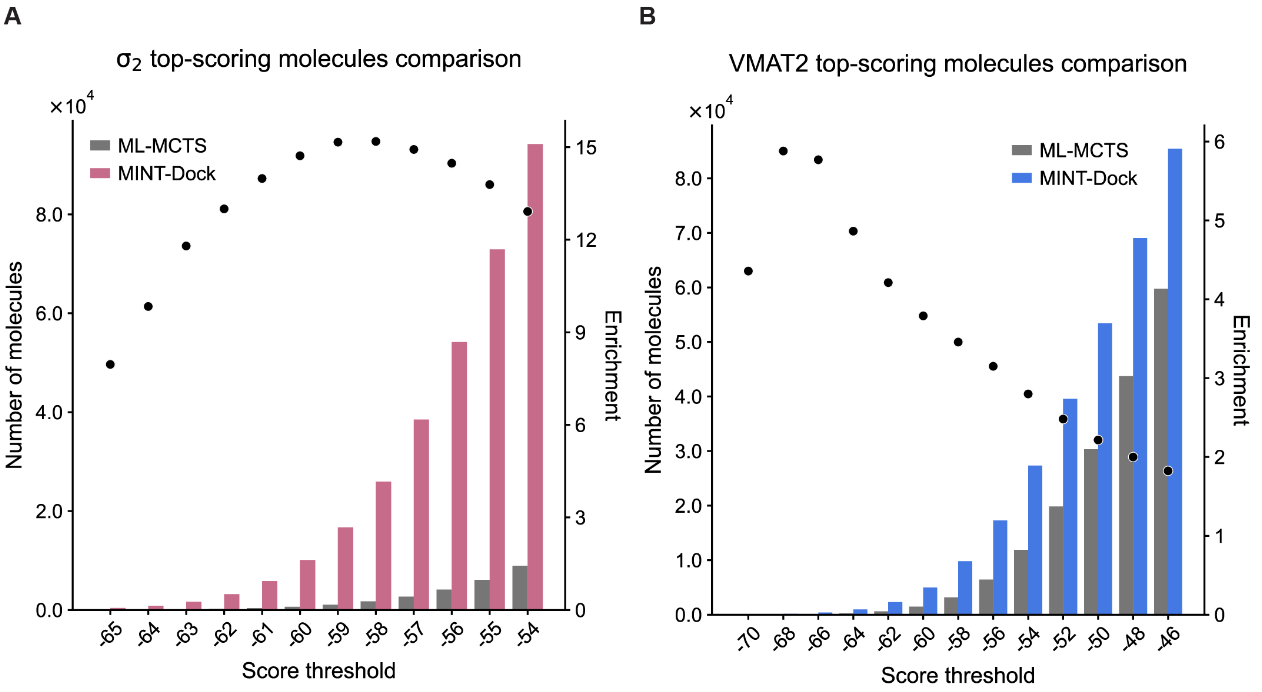


**Figure S4. Comparison between MINT-Dock and ML-MCTS of the number of well-scored molecules across different score thresholds.** (A) Comparison for $\sigma_2$. (B) Comparison for VMAT2.

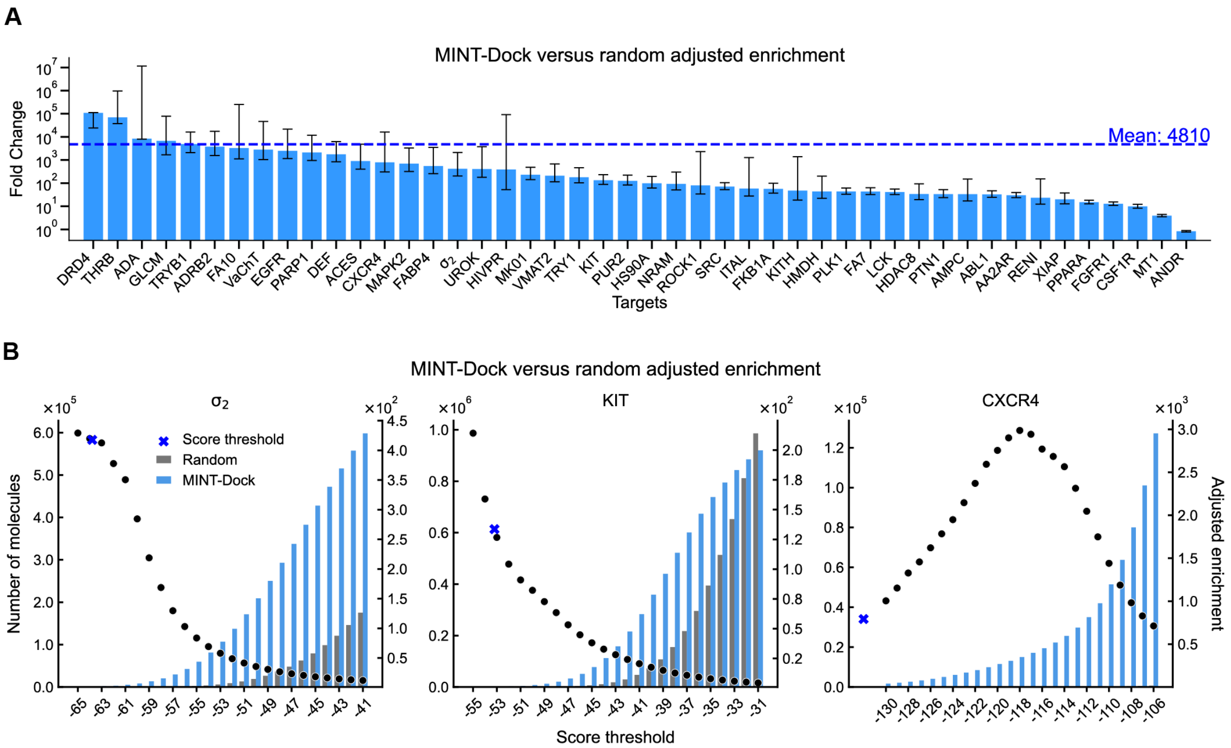


**Figure S5. Additional benchmarking of MINT-Dock in comparison with DOCK6 HDB sampled random molecules** (A) Summary plot of the adjustment enrichment ratio at a designated score threshold for all 46 targets, where the random library was docked by DOCK6 HDB. Error bars represent 90% confidence intervals, estimated by bootstrapping (1,000 iterations with replacement). (B) Plots of adjusted enrichment ratio versus docking score threshold for three example targets, where the random library was docked by DOCK6 HDB. It presents a more fine-grained enrichment comparison on three example targets. For each target, we count how many molecules achieve docking scores better than a series of thresholds and compare MINT-Dock to a size-matched random draw. The bar plots show that, across thresholds, MINT-Dock consistently yields many more molecules above any given score cutoff than random sampling, indicating that the search preferentially concentrates computation on productive regions of chemical space. The scatter points show that the adjusted enrichment is strongest at stringent cutoffs, where high-quality molecules are rare under random sampling and therefore most benefit from guided search.

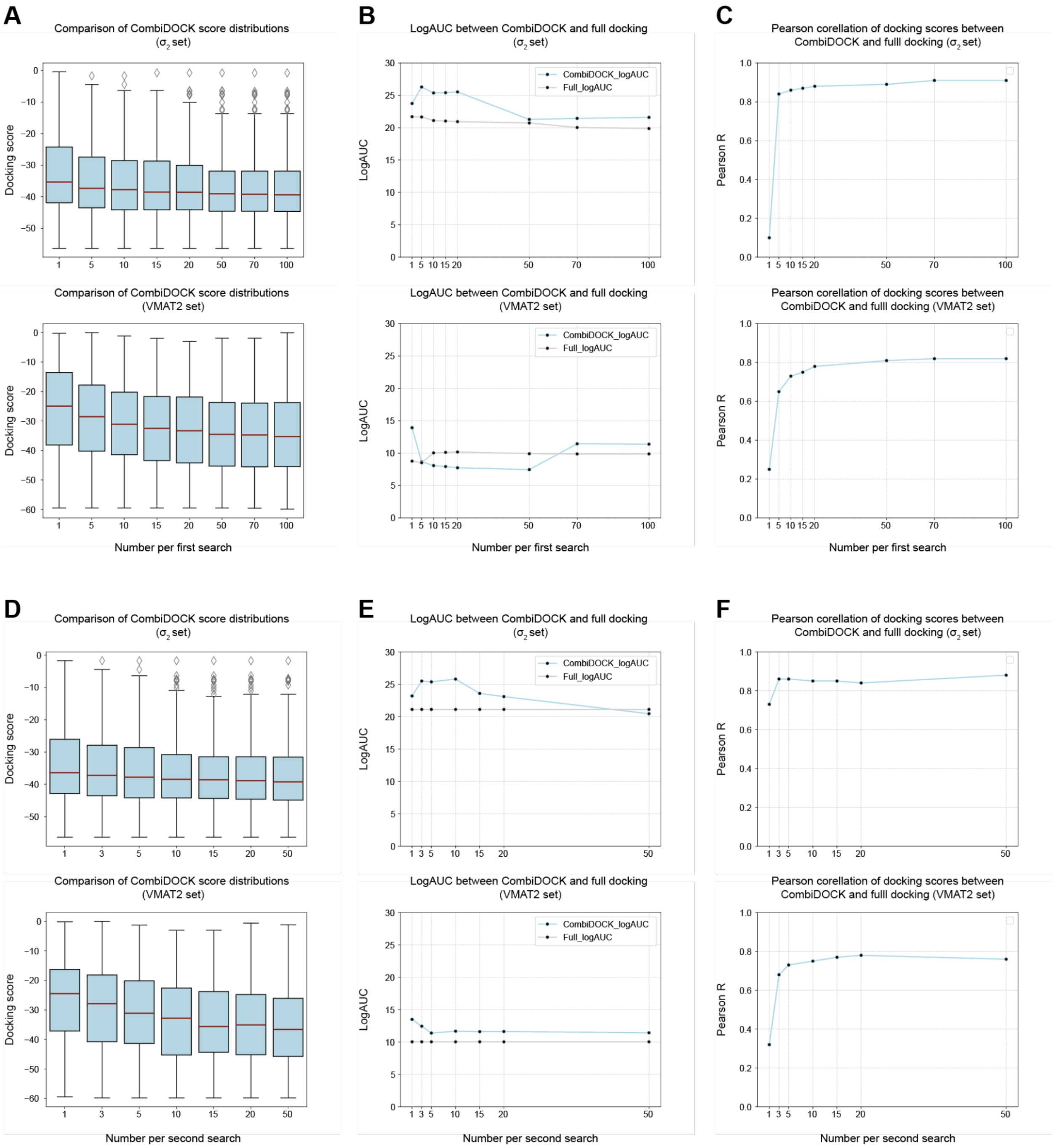


**Figure S6. Impact of the docking parameters on CombiDOCK performance across retrospective datasets.** (A) Distributions of CombiDOCK docking scores for the $\sigma_2$ (top) and VMAT2 (bottom) datasets as the "number per first search" parameter increases (1, 5, 10, 15, 20, 50, 70, 100). (B) Corresponding changes in logAUC for the $\sigma_2$ (top) and VMAT2 (bottom) datasets relative to full-molecule docking. (C) Changes in the Pearson correlation between CombiDOCK and full-molecule docking scores for the $\sigma_2$ (top) and VMAT2 (bottom) datasets. (D) Distributions of docking scores for the $\sigma_2$ (top) and VMAT2 (bottom) datasets as the "number per second search" parameter increases (1, 3, 5, 10, 15, 20, 50), with the "number per first search" fixed at its default value of 10. (E) Corresponding changes in logAUC relative to full-molecule docking. (F) Corresponding changes in Pearson correlations with full-molecule docking scores.

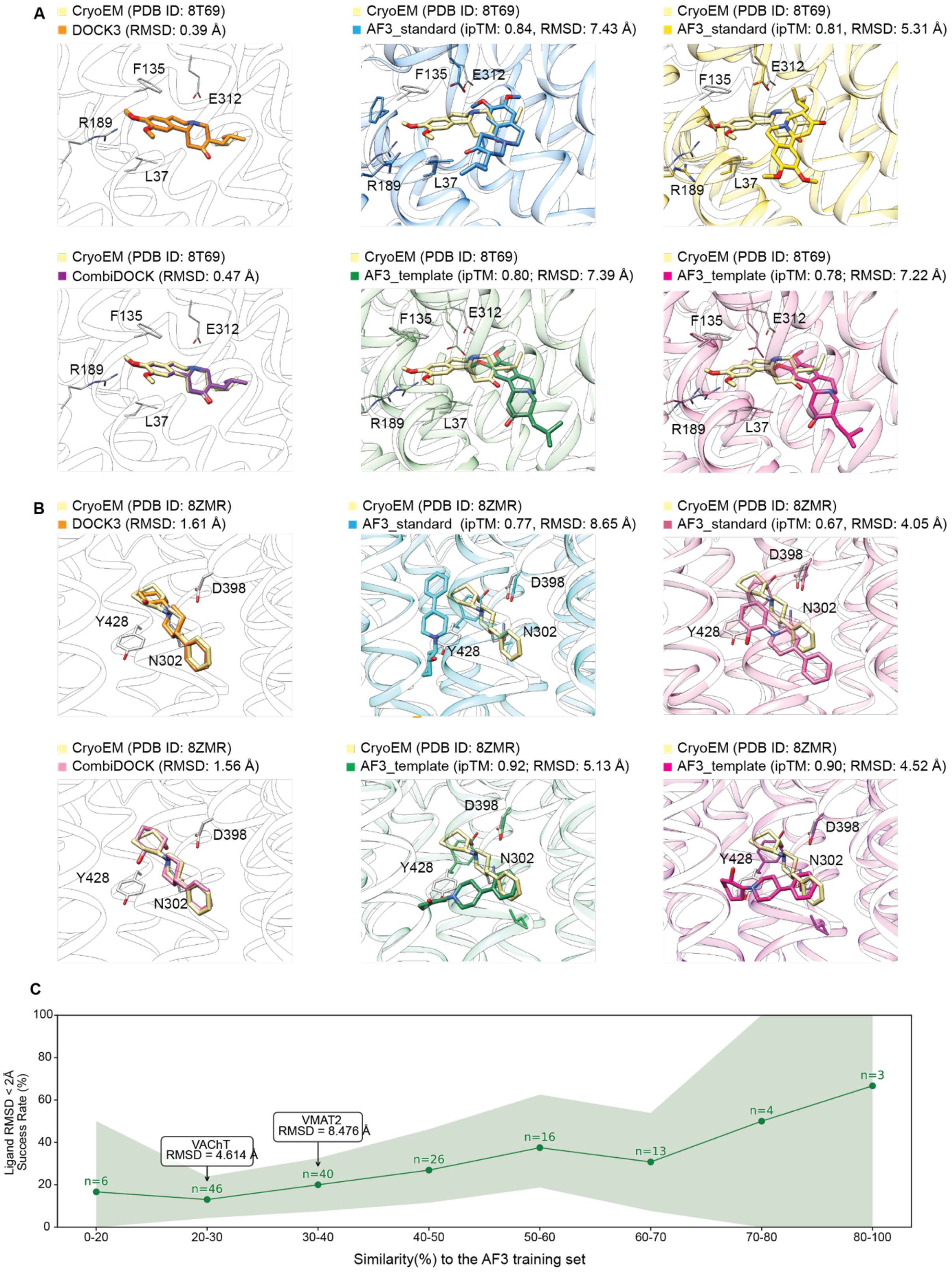


**Figure S7. Comparison of ligand binding poses predicted by DOCK3 and AlphaFold3 with experimental cryo-EM structures.** (A) Cryo-EM structure of VMAT2 bound to tetrabenazine (TBZ, PDB

ID: 8T69) overlaid with (upper left) DOCK3-predicted pose (RMSD = 0.39 Å), (upper middle) highest scoring AlphaFold3 model (ipTM = 0.84; RMSD = 7.43 Å), (upper right) lowest RMSD AlphaFold3 model (ipTM = 0.81; RMSD = 5.31 Å), (lower left) CombiDOCK-predicted pose (RMSD = 0.47 Å), (lower middle) highest scoring AlphaFold3_template model (ipTM = 0.80; RMSD = 7.39 Å), and (lower right) lowest RMSD AlphaFold3_template model (ipTM = 0.78; RMSD = 7.22 Å). (B) Cryo-EM structure of VAChT bound to vesamicol (PDB ID: 8ZMR) overlaid with (upper left) DOCK3-predicted pose (RMSD = 1.61 Å), (upper middle) highest scoring AlphaFold3 model (ipTM = 0.77; RMSD = 8.65 Å), (upper right) lowest RMSD AlphaFold3 model (ipTM = 0.67; RMSD = 4.05 Å), (lower left) CombiDOCK-predicted pose (RMSD = 1.56 Å), (lower middle) highest scoring AlphaFold3_template model (ipTM = 0.92; RMSD = 5.13 Å), and (lower right) lowest RMSD AlphaFold3_template model (ipTM = 0.90; RMSD = 4.52 Å). (C) In both cases, DOCK3 and CombiDOCK accurately recapitulate the native ligand binding poses, while AlphaFold3 predictions fail to capture the experimentally observed conformations. Benchmarking AF3 performance across the prediction of 154 SLC transporters unveils moderate correlation between prediction success and similarity of protein-ligand complex to AF3's training set.

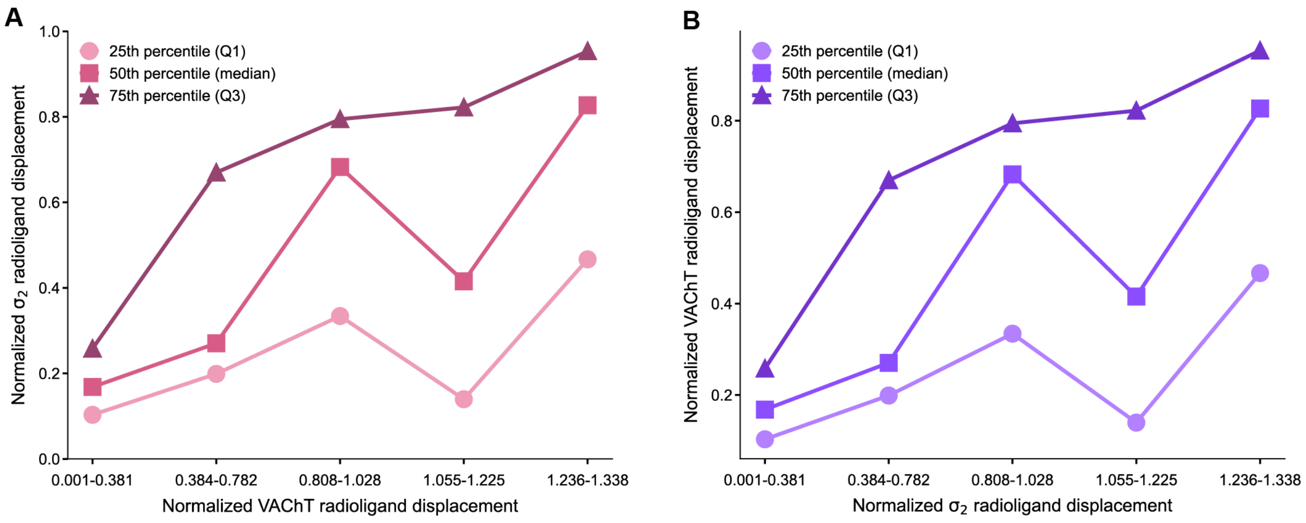


**Figure S8. Additional evaluation of VAChT campaigns.** (A) The 25%, 50%, 75% quartile of the radioligand displacement for $\sigma_2$ at different ranges of radioligand displacement for VAChT binned such that the number of compounds in each bin is roughly equal. (B) The 25%, 50%, 75% quartile of the radioligand displacement for VAChT at different ranges of radioligand displacement for $\sigma_2$ binned such that the number of compounds in each bin is roughly equal.

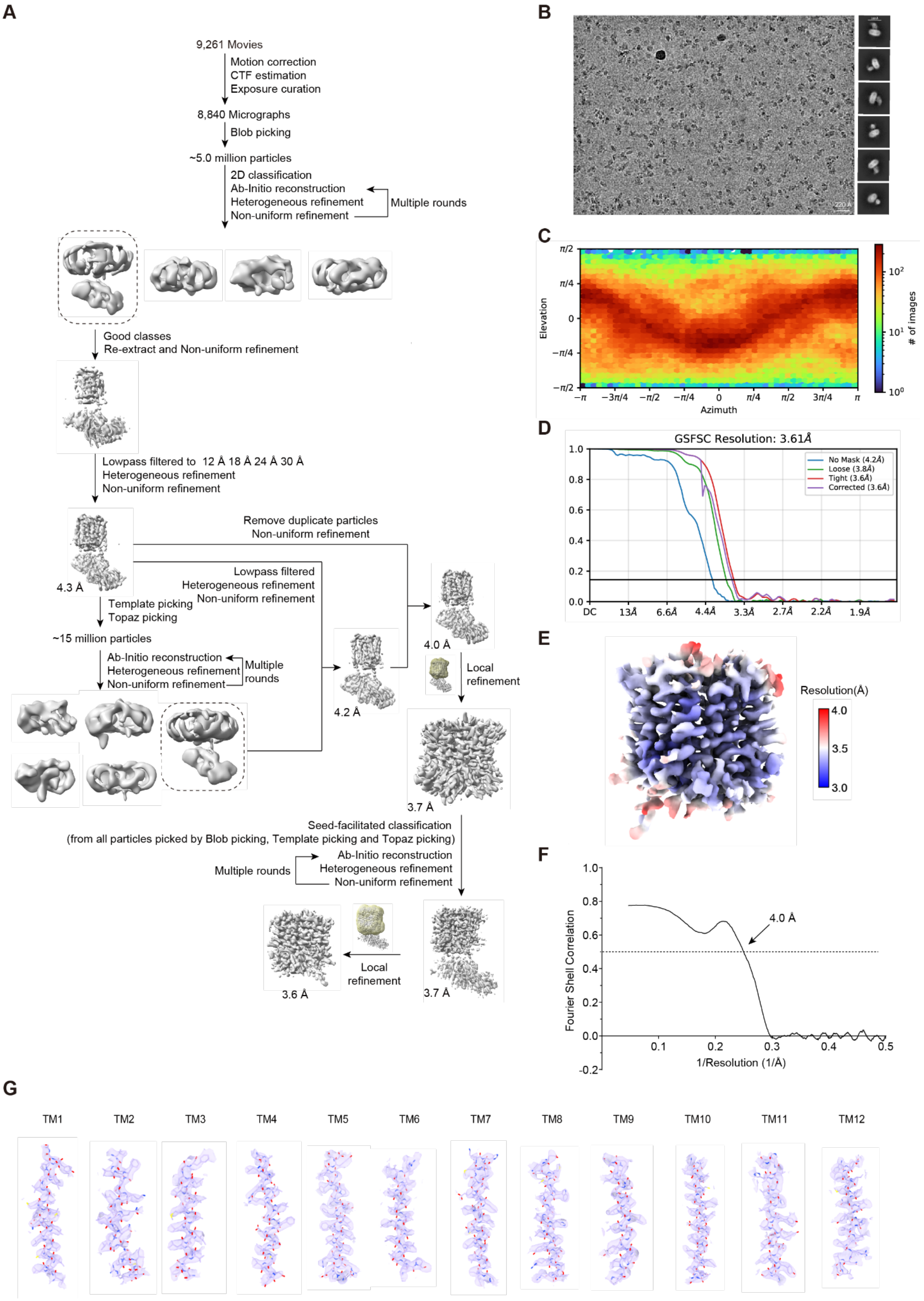

A
9,261 Movies
Motion correction
CTF estimation
Exposure curation
8,840 Micrographs
Blob picking
~5.0 million particles
2D classification
Ab-Initio reconstruction
Heterogeneous refinement
Non-uniform refinement
Multiple rounds
Good classes
Re-extract and Non-uniform refinement
Lowpass filtered to 12 Å 18 Å 24 Å 30 Å
Heterogeneous refinement
Non-uniform refinement
Remove duplicate particles
Non-uniform refinement
Lowpass filtered
Heterogeneous refinement
Non-uniform refinement
4.3 Å
Template picking
Topaz picking
~15 million particles
Ab-Initio reconstruction
Heterogeneous refinement
Non-uniform refinement
Multiple rounds
4.2 Å
4.0 Å
Local refinement
3.7 Å
Seed-facilitated classification
(from all particles picked by Blob picking, Template picking and Topaz picking)
Ab-Initio reconstruction
Heterogeneous refinement
Non-uniform refinement
Multiple rounds
Local refinement
3.6 Å
3.7 Å
B
220 Å
C
Elevation
Azimuth
### of images
D
GSFSC Resolution: 3.61Å
No Mask (4.2Å)
Loose (3.8Å)
Tight (3.6Å)
Corrected (3.6Å)
E
Resolution(Å)
4.0
3.5
3.0
F
Fourier Shell Correlation
1/Resolution (1/Å)
4.0 Å
G
TM1
TM2
TM3
TM4
TM5
TM6
TM7
TM8
TM9
TM10
TM11
TM12

**Figure S9. Cryo-EM data processing for CSM00228273820-bound VAChT.** (A) Schematic workflow of the data processing pipeline. (B) Representative cryo-EM micrograph (left) and selected 2D class averages (right). (C) Angular distribution graph of the final set of particles used for 3D reconstruction. (D) Fourier shell correlation curves indicating the overall resolution of the cryo-EM map, calculated between two half-maps. (E) Local resolution map of the final reconstruction. (F) Fourier shell correlation curve for model-map correlation. (G) Cryo-EM densities of the transmembrane helices, shown with the final atomic model.

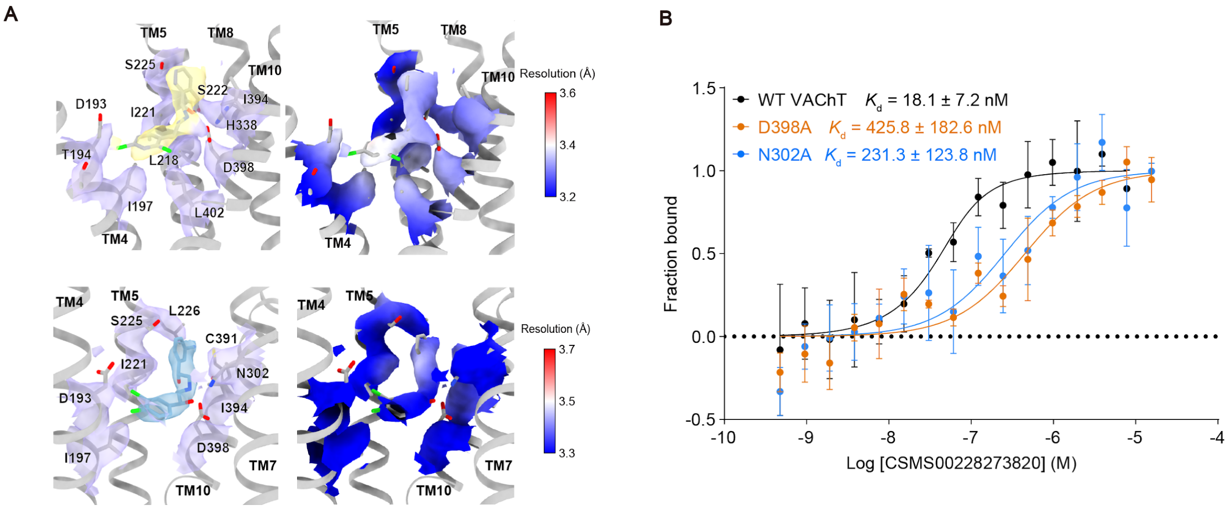


**Figure S10. Structural validation of VAChT binding with CSMS00228273820. MST analysis of the binding affinity between VAChT variants and CSMS00228273820.** (A) Close-up view of the VAChT binding pocket showing CSM00228273820 surrounded by transmembrane helices TM4, TM5, TM8, and TM10. The key interacting residues (T194, I197, I221, S222, S225, H338, I394, D398, and L402) are indicated (top left). Close-up view of the VAChT binding pocket showing Z9424360192 surrounded by transmembrane helices TM4, TM5, TM7, and TM10 (bottom left). The key interacting residues (I197, D193, I221, S225, L226, N302, C391, D398) are indicated. The right panel shows the local resolution map of the binding cavity, demonstrating well-resolved density around both ligands. (B) MST analysis of the binding affinity between VAChT variants and CSMS00228273820. Experiments were performed in triplicate (n = 3, data shown as mean ± SD).

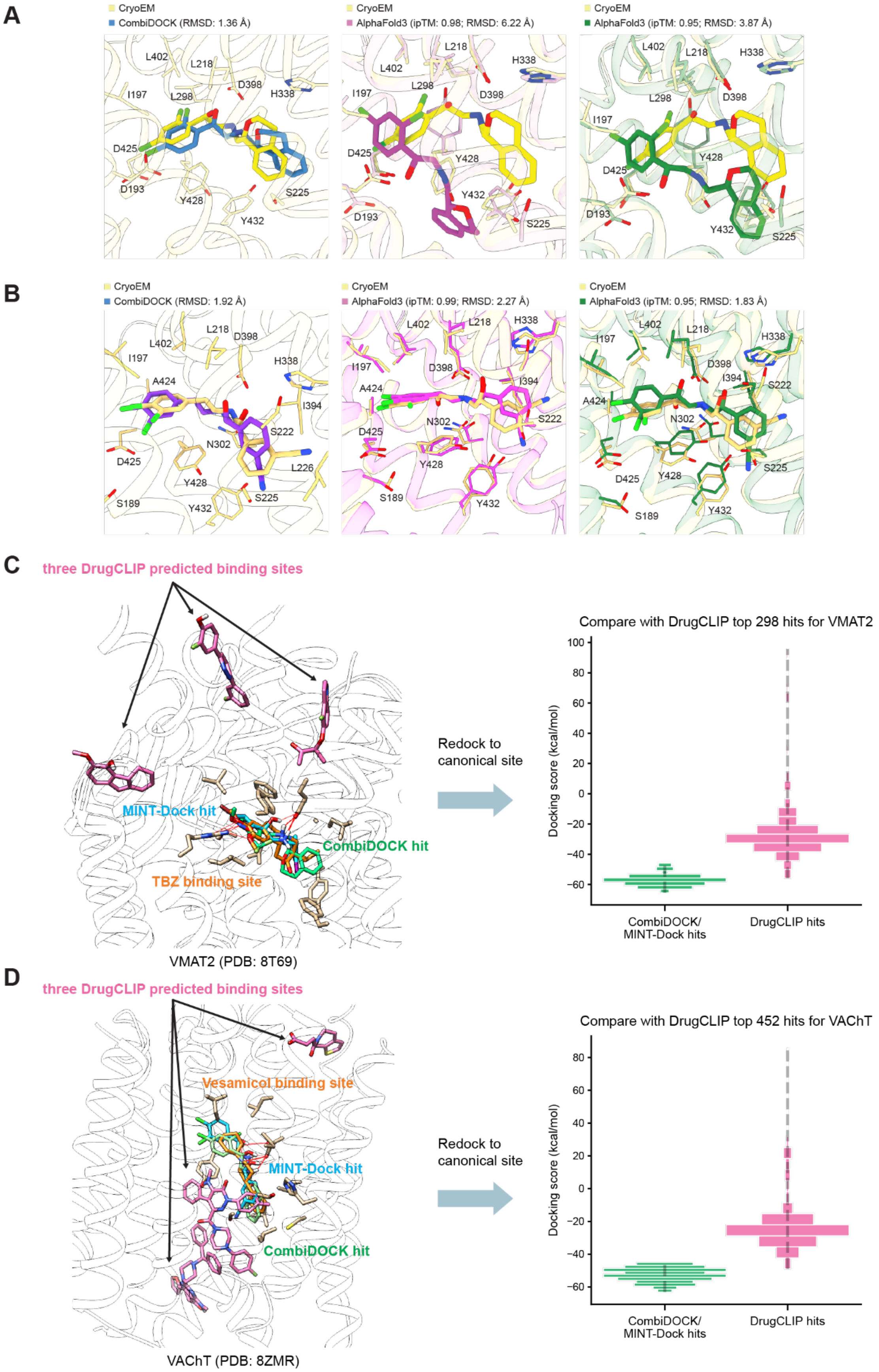

A
CryoEM
CombiDOCK (RMSD: 1.36 Å)
CryoEM
AlphaFold3 (ipTM: 0.98; RMSD: 6.22 Å)
CryoEM
AlphaFold3 (ipTM: 0.95; RMSD: 3.87 Å)
B
CryoEM
CombiDOCK (RMSD: 1.92 Å)
CryoEM
AlphaFold3 (ipTM: 0.99; RMSD: 2.27 Å)
CryoEM
AlphaFold3 (ipTM: 0.95; RMSD: 1.83 Å)
C
three DrugCLIP predicted binding sites
MINT-Dock hit
CombiDOCK hit
TBZ binding site
VMAT2 (PDB: 8T69)
Redock to canonical site
Compare with DrugCLIP top 298 hits for VMAT2
Docking score (kcal/mol)
CombiDOCK/ MINT-Dock hits
DrugCLIP hits
D
three DrugCLIP predicted binding sites
Vesamicol binding site
MINT-Dock hit
CombiDOCK hit
VAChT (PDB: 8ZMR)
Redock to canonical site
Compare with DrugCLIP top 452 hits for VAChT
Docking score (kcal/mol)
CombiDOCK/ MINT-Dock hits
DrugCLIP hits

**Figure S11. Comparison of ligand binding poses in VAChT and VMAT2 predicted by CombiDOCK and deep learning methods with experimental cryo-EM structures.** (A) Cryo-EM structure of CSM00228273820 bound to VAChT overlaid with (left) CombiDOCK-predicted pose (RMSD = 1.36 Å), (middle) highest scoring AlphaFold3 model (ipTM = 0.98; RMSD = 6.22 Å), and (right) lowest RMSD AlphaFold3 model (ipTM = 0.95; RMSD = 3.87 Å). CombiDOCK accurately recapitulates the native CSM00228273820 binding pose, while AlphaFold3 predictions fail to capture the native pose. (B) Cryo-EM structure of Z9424360192 bound to VAChT overlaid with (left) CombiDOCK-predicted pose (RMSD = 1.92 Å), (middle) highest scoring AlphaFold3 model (ipTM = 0.99; RMSD = 2.27 Å), and (right) lowest RMSD AlphaFold3 model (ipTM = 0.95; RMSD = 1.83 Å). Both CombiDOCK and AF3 accurately recapitulate the native Z9424360192 binding pose. (C) DrugCLIP predicted binding sites in VMAT2 (PDB: 8T69), compared to the canonical binding site of known VMAT2 inhibitor TBZ and our best CombiDOCK and MINT-Dock hit. Visual inspection of predicted DrugCLIP hits reveal that the method fails to capture the canonical binding site (left). An inferior docking score distribution is observed when redocking top DrugCLIP hits to the canonical site when compared to CombiDOCK and MINT-Dock experimental hits (normalized radioligand displacement < 0.5) (right). (D) DrugCLIP predicted binding sites in VAChT (PDB: 8ZMR), compared to the canonical binding site of known VAChT inhibitor vesamicol and our best CombiDOCK and MINT-Dock hits. Similarly to our VMAT2 DrugCLIP analysis, through visual inspection, VAChT DrugCLIP hits are observed to be spatially distant to the canonical binding site (left). Furthermore, when redocked to the canonical site, top DrugCLIP hits are observed to have a worse docking score distribution when compared to top CombiDOCK and MINT-Dock experimental hits (normalized radioligand displacement < 0.5) (right).

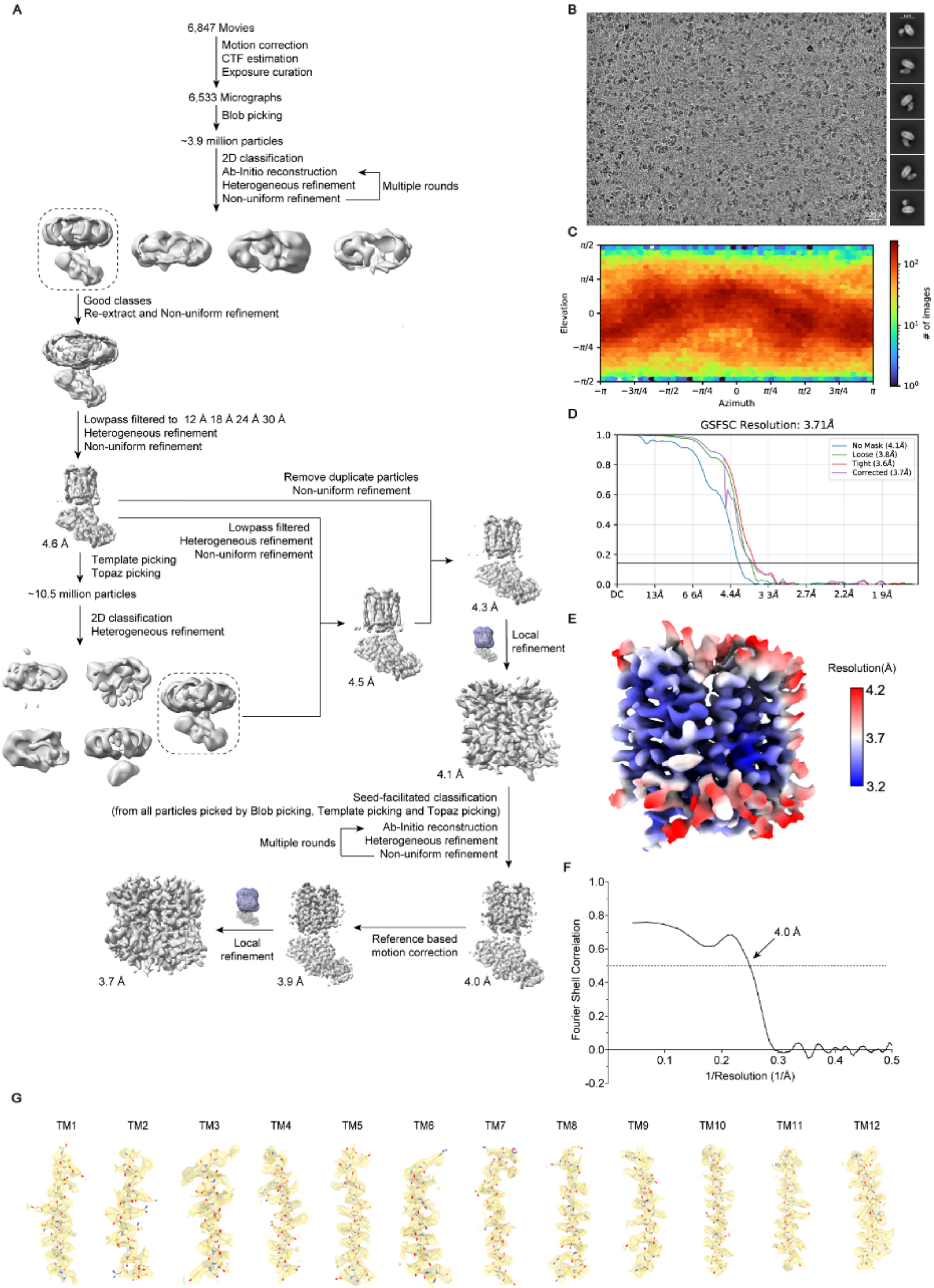
A
6,847 Movies
Motion correction
CTF estimation
Exposure curation
6,533 Micrographs
Blob picking
~3.9 million particles
2D classification
Ab-Initio reconstruction
Heterogeneous refinement
Non-uniform refinement
Multiple rounds
Good classes
Re-extract and Non-uniform refinement
Lowpass filtered to 12 Å 18 Å 24 Å 30 Å
Heterogeneous refinement
Non-uniform refinement
4.6 Å
Remove duplicate particles
Non-uniform refinement
Lowpass filtered
Heterogeneous refinement
Non-uniform refinement
Template picking
Topaz picking
~10.5 million particles
2D classification
Heterogeneous refinement
4.5 Å
4.3 Å
Local refinement
4.1 Å
Seed-facilitated classification
(from all particles picked by Blob picking, Template picking and Topaz picking)
Ab-Initio reconstruction
Heterogeneous refinement
Non-uniform refinement
Multiple rounds
Reference based motion correction
Local refinement
3.7 Å
3.9 Å
4.0 Å
B
C
Elevation
Azimuth
### of images
D
GSFSC Resolution: 3.71Å
No Mask (4.1Å)
Loose (3.8Å)
Tight (3.6Å)
Corrected (3.7Å)
E
Resolution(Å)
4.2
3.7
3.2
F
Fourier Shell Correlation
1/Resolution (1/Å)
4.0 Å
G
TM1
TM2
TM3
TM4
TM5
TM6
TM7
TM8
TM9
TM10
TM11
TM12

**Figure S12. Cryo-EM data processing for Z9424360192-bound VAChT.** (A) Image processing workflow for VAChT-Z9424360192 complex. (B) Representative raw micrograph (left) and selected 2D class averages (right). (C) Angular distribution of particles used for final 3D reconstruction. (D) Fourier shell correlation (FSC) curves between two half maps demonstrating map resolution at 0.143 criterion. (E) Local resolution estimation of the cryo-EM density map. (F) FSC curve calculated between the cryo-EM map and structural model. (G) Cryo-EM densities for transmembrane (TM) helices.

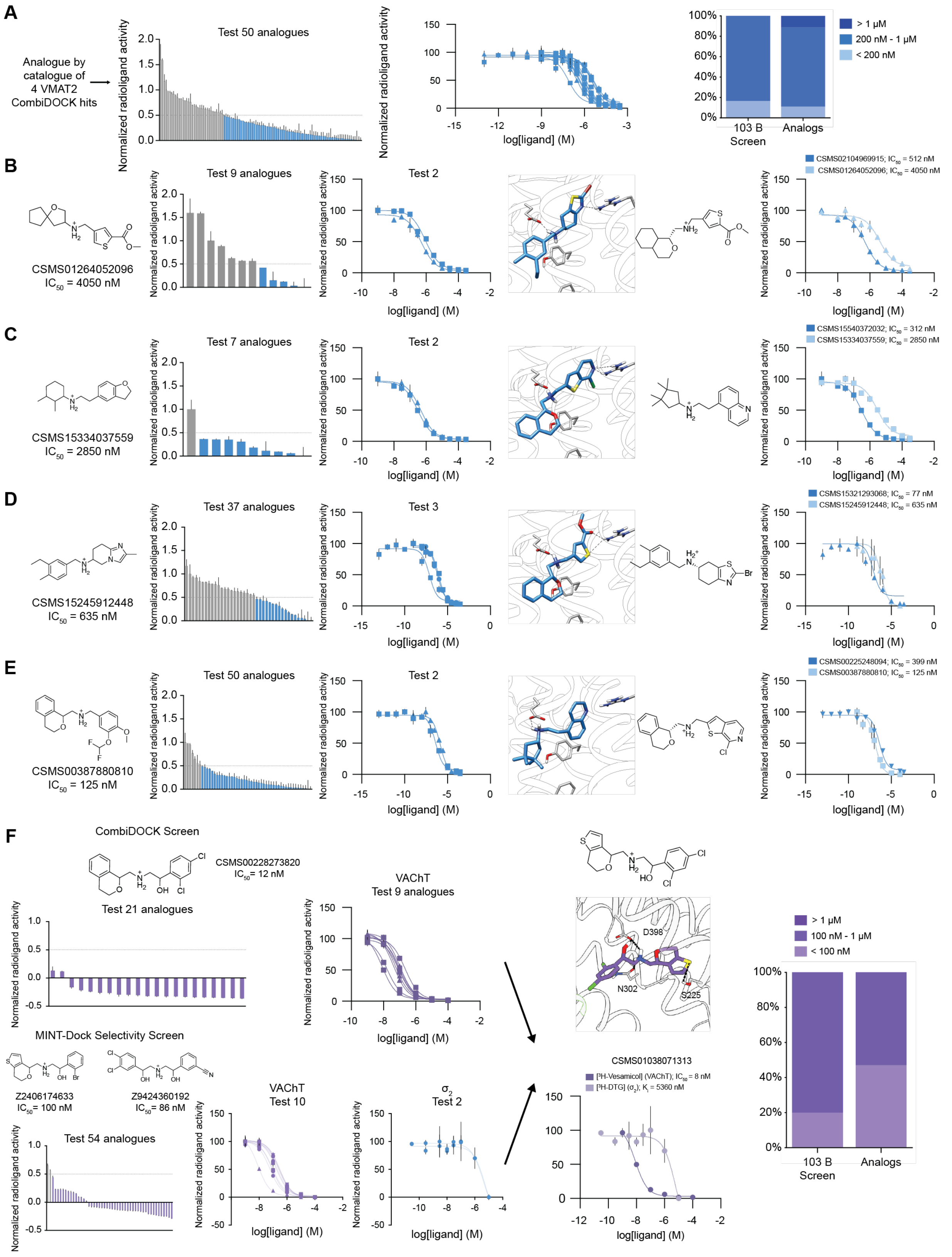

A
Analogue by catalogue of 4 VMAT2 CombiDOCK hits
Test 50 analogues
Normalized radioligand activity
log[ligand] (M)
100%
80%
60%
40%
20%
0%
103 B Screen
Analogs
> 1 μM
200 nM - 1 μM
< 200 nM
B
CSMS01264052096
IC50 = 4050 nM
Test 9 analogues
Test 2
CSMS02104969915; IC50 = 512 nM
CSMS01264052096; IC50 = 4050 nM
C
CSMS15334037559
IC50 = 2850 nM
Test 7 analogues
Test 2
CSMS15540372032; IC50 = 312 nM
CSMS15334037559; IC50 = 2850 nM
D
CSMS15245912448
IC50 = 635 nM
Test 37 analogues
Test 3
CSMS15321293068; IC50 = 77 nM
CSMS15245912448; IC50 = 635 nM
E
CSMS00387880810
IC50 = 125 nM
Test 50 analogues
Test 2
CSMS00225248094; IC50 = 399 nM
CSMS00387880810; IC50 = 125 nM
F
CombiDOCK Screen
CSMS00228273820
IC50 = 12 nM
Test 21 analogues
VAChT
Test 9 analogues
MINT-Dock Selectivity Screen
Z2406174633
IC50 = 100 nM
Z9424360192
IC50 = 86 nM
Test 54 analogues
VAChT
Test 10
σ2
Test 2
D398
N302
S225
CSMS01038071313
[3H-Vesamicol] (VAChT); IC50 = 8 nM
[3H-DTG] (σ2); Ki = 5360 nM
> 1 μM
100 nM - 1 μM
< 100 nM
103 B Screen
Analogs

**Figure S13. Analogue maturation of discovered VMAT2 and VAChT binders.** (A) 103 close-structural analogues of four VMAT2 hits from the 103B screen were synthesized and tested in a [$^3$H]-DA uptake assay, using >50% [$^3$H]-DA displacement as a hit threshold. 72 out of the 103 analogues were hits (left). The $IC_{50}$ was determined for 9 of the best binders representing affinities ranging from the low-micromolar to the hundreds of nanomolar range (middle). The $IC_{50}$ distribution did not shift significantly when compared to the 103 billion CombiDOCK screen against VMAT2 (right). (B) Nine analogues were curated and tested for low-affinity representative CSMS01264052096. Four of these analogues exhibited >50% inhibition (left). The $IC_{50}$ was determined for the top 2 binders, CSMS02104969915 with an $IC_{50}$ = 521 nM, and CSMS15184673800 with an $IC_{50}$ = 887 nM (middle). This represents an 8-fold improvement in potency from the parent compound CSMS01264052096 (right). (C) Seven analogues were curated and tested for our second low-affinity representative CSMS15334037559, all exhibited >50% inhibition (left). The $IC_{50}$ was determined for two binders, CSMS15334037147 with an $IC_{50}$ of 472 nM, and CSMS15540372032 with an $IC_{50}$ of 312 nM (middle). This represents a 9-fold improvement in potency from the parent compound CSMS15334037559 (right). (D), 37 analogues were synthesized and tested for high-potency representative CSMS15245912448, 17 of which exhibited >50% inhibition (left). The $IC_{50}$ was determined for three of the best analogues, with an $IC_{50}$ ranging between 77 nM to 963 nM (middle). The most potent compound, CSMS15321293068 with an $IC_{50}$ of 77 nM represents an 8-fold improvement in potency (right). (E) 50 analogues were synthesized and tested for our most potent high-affinity representative, CSMS00387880810, 44 of which exhibited >50% inhibition (left). The $IC_{50}$ was determined for two analogues, CSMS00225248094 with an $IC_{50}$ of 407 nM, and CSCS12219892184 with an $IC_{50}$ of 1410 nM (middle). The most potent analogue CSMS00225248094, did not exceed potencies exhibited by the parent compound, CSMS00387880810 (right). (F) 21 close structural analogues were curated for the most potent hit discovered from the VAChT CombiDOCK 103 billion screen, CSMS00228273820 ($IC_{50}$ = 12 nM) (top left) The21 analogues tested in a [$^3$H]-vesamicol binding assay, using >50% [$^3$H]-vesamicol displacement as a hit threshold.. Dosage response curves for 9 of the best binders were determined to have an $IC_{50}$ range between 8 and 221 nM (top middle). Similarly, for two MINT-Dock selective hits, the same analog search was conducted (22 Z9424360192 analogs; 32 Z2406174633 analogs) (bottom left). Dosage response curve for 10 of the best binders were determined to have an $IC_{50}$ range between 8 and 300 nM for VAChT, while maintaining selectivity over $\sigma_2$ (bottom middle). Congeneric expansion of series CSMS00228273820 and series Z2406174633, facilitated discovery of CSMS01038071313, the most potent analogue from both the CombiDOCK analogues and MINT-Dock selective analogues (top right). The most potent analogue, CSMS01038071313 ($IC_{50}$ = 8 nM), was observed to maintain some degree of VAChT/$\sigma_2$ selectivity (bottom right). The $IC_{50}$ distribution was observed to evolve towards more potent analogues from the 103 billion CombiDOCK screen to the analoging screens (right).

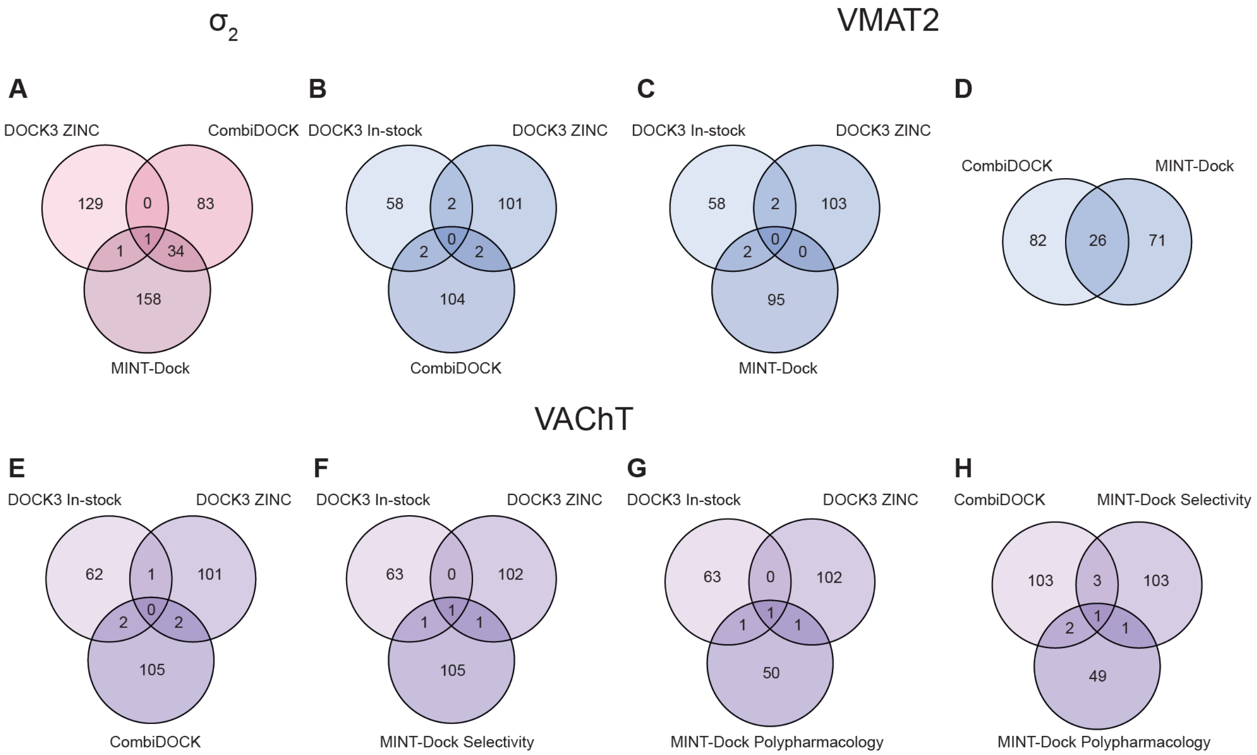


**Figure S14. Novelty analysis of tested molecules across different screens.** Venn diagrams illustrating the overlap of Bemis-Murcko scaffolds among molecules tested for σ2, VMAT2, and VAChT targets across multiple virtual screening campaigns using DOCK3, CombiDOCK, and MINT-Dock guided search. (A-B) Scaffolds overlap for all tested compounds for the following σ2 screens: DOCK3 ZINC 490 million screen, CombiDOCK 56 billion screen, MINT-Dock 103 billion screen. (C-D) Scaffolds overlap for all tested compounds for the following VMAT2 screens: DOCK3 in-stock 3.6 million screen, DOCK3 ZINC 490 million screen, CombiDOCK 56 billion screen, MINT-Dock 103 billion screen. (E-H) Scaffolds overlap for all tested compounds for the following VAChT screens: DOCK3 in-stock 3.6 million screen, DOCK3 ZINC 490 million screen, CombiDOCK 56 billion screen, MINT-Dock 103 billion selectivity screen, MINT-Dock 103 billion polypharmacology screen.

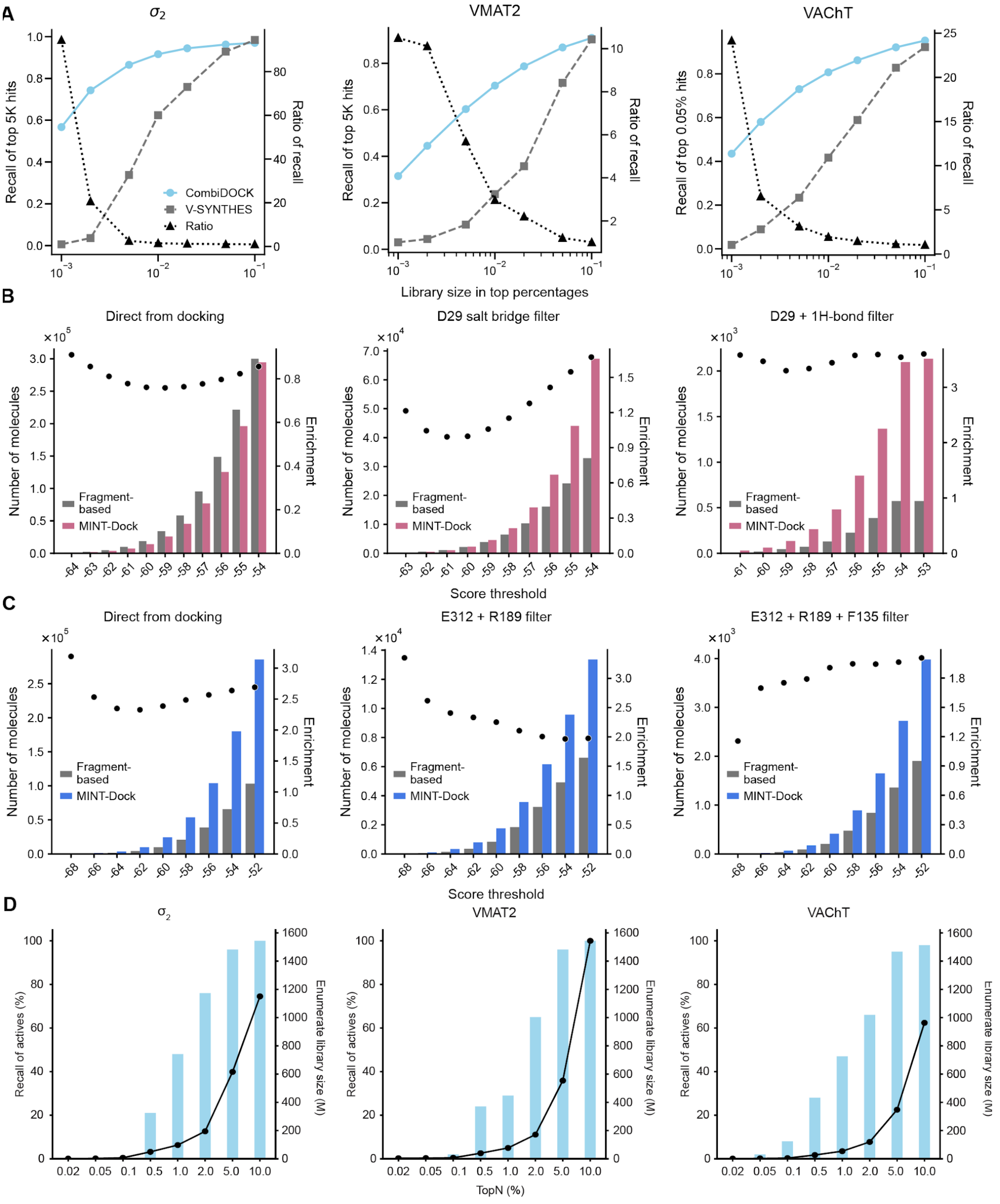


**Figure S15. Summary benchmarks of CombiDOCK/MINT-Dock comparison with fragment-based docking methods**. (A) Recall performance between CombiDOCK and V-SYNTHES was evaluated for the targets $\sigma_2$, VMAT2, and VAChT, respectively. The top 0.05% of compounds (top 5,000 molecules from a 10-million compound library) from the full-molecule docking were used as reference hits. (B) Comparison between MINT-Dock and fragment-based docking of the number of hits among top 300,000 scoring compounds and after passing each interaction filter at each score threshold for $\sigma_2$. (C) Comparison between

MINT-Dock and fragment docking of the number of hits among Top 300,000 scoring compounds and after passing each interaction filter at each score threshold for VMAT2. (D) Plots show recall of CombiDOCK actives (left y-axis) and corresponding enumerated library sizes (right y-axis) as a function of the top-N% threshold. Results are presented for three targets: $\sigma_2$, VMAT2, and VAChT. For each target, recall increases with a broader top-N% inclusion, reflecting enhanced recovery of active compounds, while enumerated library size (in millions) scales proportionally with the threshold.

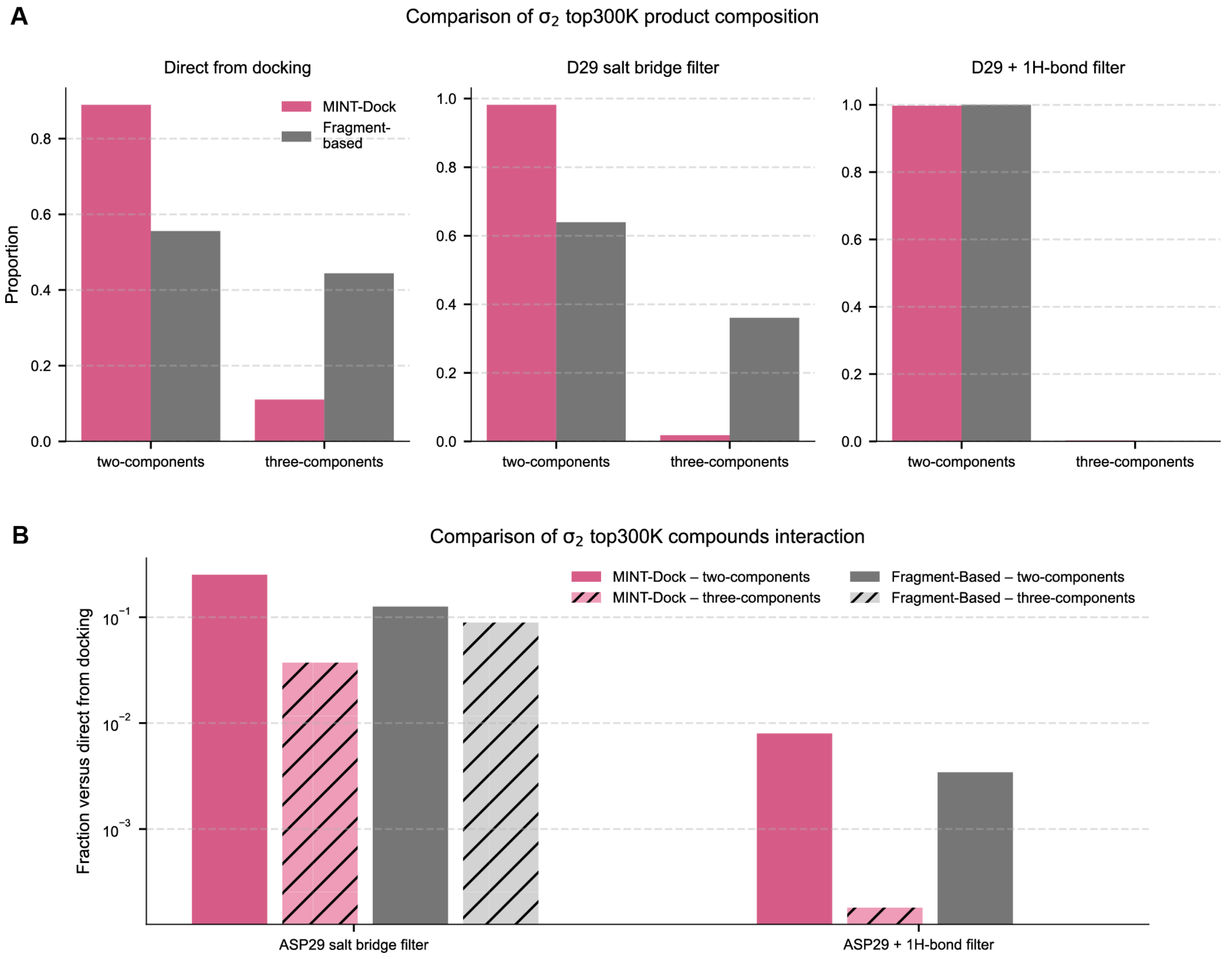


**Figure S16. Composition analysis of top 300,000 compounds from $\sigma_2$ MINT-Dock and fragment-based screen.** (A) Comparison of the proportions of two-component and three-component products among all compounds at different stages of filtering for each method. (B) Comparison of the ratios of two-component and three-component products that passed interaction filters for each method.

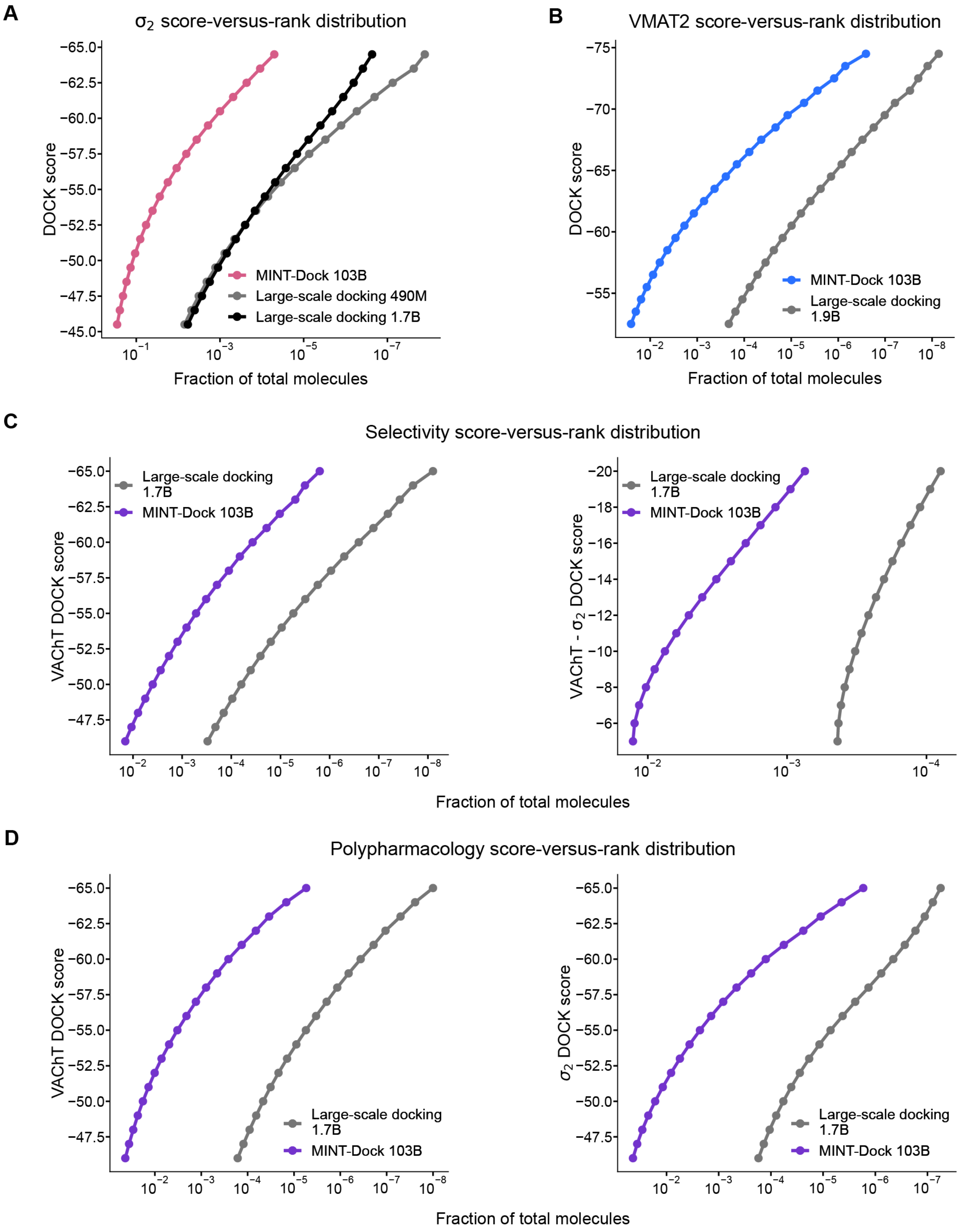


**Figure S17. MINT-Dock improves efficiency across prospective campaigns.** (A) Comparison of the score-versus-rank distribution between two large-scale docking campaigns and MINT-Dock for $\sigma_2$. (B) Comparison of the score-versus-rank distribution between two large-scale docking campaigns and MINT-

Dock for VMAT2. (C) Comparison of the VAChT-score-versus-rank distribution and the distribution of the difference in two targets' docking scores versus rank between large-scale docking and MINT-Dock for selectivity screening. (D) Comparison of the VAChT-score-versus-rank distribution and $\sigma_2$-score-versus-rank distribution between large-scale docking and MINT-Dock for poly-pharmacology screening.

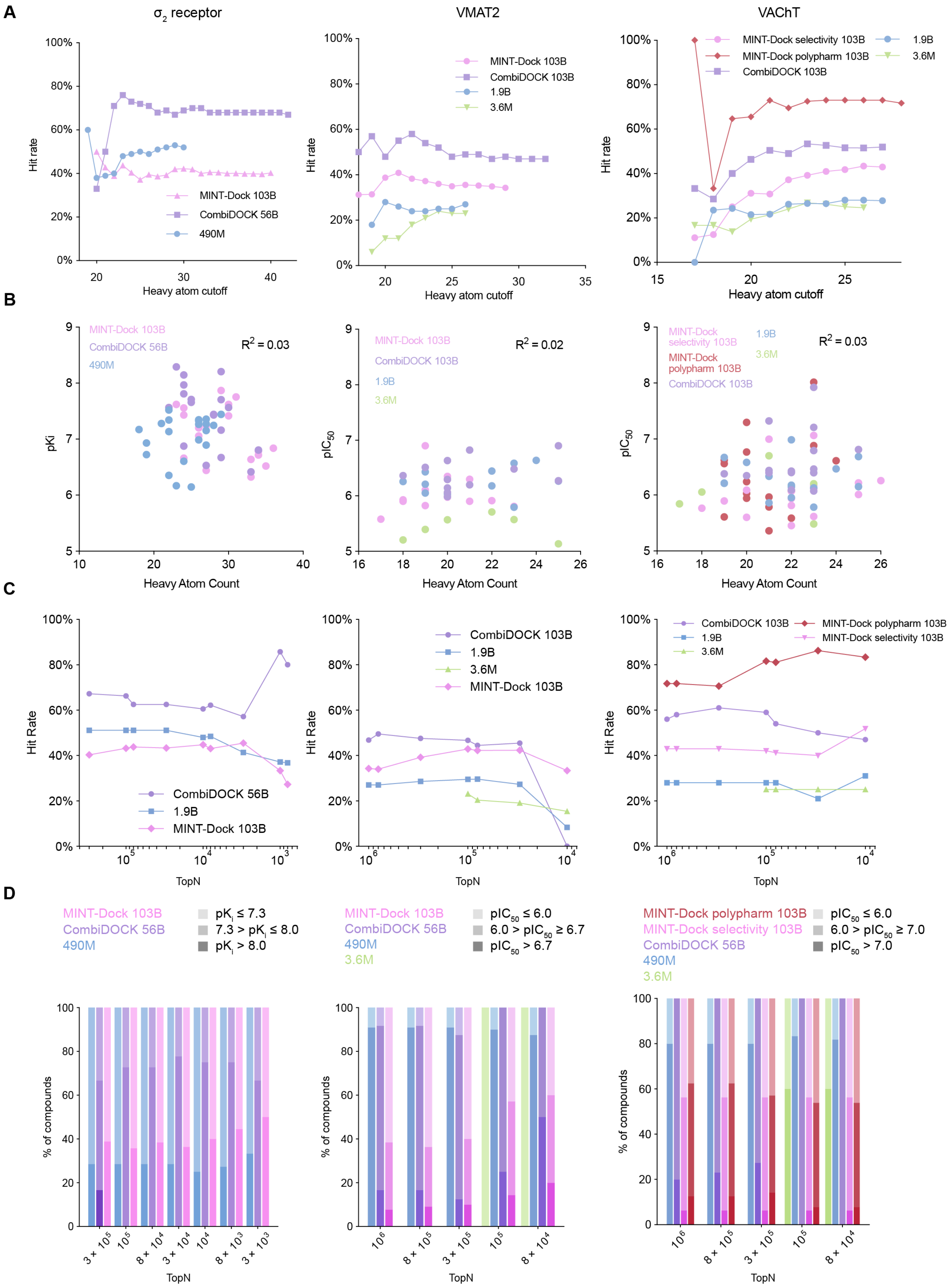

A
σ2 receptor
VMAT2
VAChT
Hit rate
Heavy atom cutoff
MINT-Dock 103B
CombiDOCK 56B
490M
CombiDOCK 103B
1.9B
3.6M
MINT-Dock selectivity 103B
MINT-Dock polypharm 103B
B
pKi
pIC50
Heavy Atom Count
R2 = 0.03
R2 = 0.02
R2 = 0.03
C
Hit Rate
TopN
D
pKi ≤ 7.3
7.3 > pKi ≤ 8.0
pKi > 8.0
pIC50 ≤ 6.0
6.0 > pIC50 ≥ 6.7
pIC50 > 6.7
6.0 > pIC50 ≥ 7.0
pIC50 > 7.0
% of compounds
TopN

**Figure S18. Ligand size effects and global ranking-based ligand discovery performance across different screening campaigns.** (A) Hit rates as a function of increasing ligand heavy-atom cutoffs for $\sigma_2$ (left), VMAT2 (middle), and VAChT (right) across libraries of different sizes. (B) Correlations between measured binding affinities and ligand heavy-atom counts for $\sigma_2$ (left), VMAT2 (middle), and VAChT (right). (C) Hit rate evolution as a function of the global ranking position of tested compounds within the topN set for $\sigma_2$ (left), VMAT2 (middle), and VAChT (right) campaigns, evaluated under controlled overlapping-ranking constraints. (D) Affinity distribution as a function of the global ranking position of tested compounds within the topN set for $\sigma_2$ (left), VMAT2 (middle), and VAChT (right) campaigns, under controlled overlapping-ranking constraints.

| Docking strategy | Library preparation time | Library storage | Docking time |
|---|---|---|---|
| On-the-fly sampling (e.g., Autodock Vina) | ~1.6 years (1 s/mol) | 200 TB (2 KB/mol) | ~48 years (30 s/mol)[a] |
| Pre-sampling (e.g., DOCK3.7)[b] | ~48 years (30 s/mol) | 50,000 TB (0.5 MB/mol) | ~1.6 years (1 s/mol)[a] |
| CombiDOCK | ~1 day | 1TB | 0.07 s/mol |

**Table S1. Estimated total costs of docking a 100-billion-compound library on a 2000-core cluster.** [a]curated from (Bender et al, 2021, doi: 10.1038/s41596-021-00597-z) [b]referred to as traditional docking methods in **Fig. 1A.**

| Protein type | Number of Targets | Targets |
|---|---|---|
| Other enzymes | 13 | ACES, ADA, AMPC, DEF, GLCM, HDAC8, HMDH, PARP1, PUR2, NRAM, FKB1A, PTN1, KITH |
| Protein kinases | 11 | ABL1, KIT, EGFR, FGFR1, CSF1R, SRC, LCK, PLK1, MAPK2 , MK01, ROCK1 |
| Proteases | 8 | HIVPR, RENI, TRY1, TRYB1, UROK, FA7, FA10, THRB |
| GPCRs | 5 | ADRB2, DRD4, AA2AR, CXCR4, MT1 |
| Nuclear receptors | 2 | ANDR, PPARA |
| Transporters | 2 | VMAT2, VAChT |
| Miscellaneous | 5 | XIAP, HS90A, ITAL, FABP4, $\sigma_2$ |
| Total | 46 | |

**Table S2. Protein-type classification of the 46 benchmark targets.**

| Docking methods | Average LogAUC[a] |
|---|---|
| DLIGAND2 | 9.64 |
| AutoDock Vina | 9.87 |
| ΔVinaRF20 | 9.45 |
| DLIGAND | 8.16 |
| X-ScoreHM | 7.75 |
| ID-Score | 2.23 |
| **DOCK6 HDB docking** | **28.5** |
| **CombiDOCK** | **27.2** |

**Table S3. LogAUC performance of different scoring functions or docking engine on DUDE-Z targets.** [a]The data of other docking methods is curated from (Chen, Pin, et al, 2019, doi: 10.1186/s13321-019-0373-4)

| | CSMS00228273820-bound hVAChT (EMDB-xxxxx) (PDB xxxx) | Z9424360192-bound hVAChT (EMDB-xxxxx) (PDB xxxx) |
|---|---|---|
| **Data collection and processing** | | |
| Magnification | 105,000 | 105,000 |
| Voltage (kV) | 300 | 300 |
| Electron exposure ($e^{-}/Å^2$) | 60 | 60 |
| Defocus range (μm) | 1.0−2.0 | 1.0−2.0 |
| Pixel size (Å) | 0.83 | 0.83 |
| Symmetry imposed | C1 | C1 |
| Initial particle images (no.) | 19,648,366 | 14,352,039 |
| Final particle images (no.) | 212,055 | 213,036 |
| Map resolution (Å) | 3.6 | 3.7 |
| FSC threshold | 0.143 | 0.143 |
| Map resolution range (Å) | 3.0−3.9 | 3.2−4.3 |
| **Refinement** | | |
| Initial model used (PDB code) | 8ZMR | 8ZMR |
| Model resolution (Å) | 4.0 | 4.0 |
| FSC threshold | 0.5 | 0.5 |
| Model resolution range (Å) | - | - |
| Map sharpening *B* factor ($Å^2$) | -191.7 | -160.0 |
| Model composition | | |
| Non-hydrogen atoms | 3,012 | 3,012 |
| Protein residues | 395 | 395 |
| Ligands | 1 (CSM00228273820) | 1 (Z9424360192) |
| *B* factors ($Å^2$) | | |
| Protein | 65.02 | 70.49 |
| Ligand | 65.12 | 68.55 |
| R.m.s. deviations | | |
| Bond lengths (Å) | 0.006 | 0.006 |
| Bond angles (°) | 1.090 | 1.115 |
| Validation | | |
| MolProbity score | 1.47 | 1.57 |
| Clashscore | 4.90 | 5.39 |
| Poor rotamers (%) | 0.00 | 0.00 |
| Ramachandran plot | | |
| Favored (%) | 96.66 | 95.89 |
| Allowed (%) | 3.34 | 4.11 |
| Disallowed (%) | 0.00 | 0.00 |

**Table S4. Cryo-EM data collection, refinement, and validation statistics for CSMS00228273820 and Z9424360192 bound hVAChT.**

| | Top 0.02% (500) | Top 0.05% (1295) | Top 0.1% (2590) | Top 0.2% (5180) | Top 0.5% (12951) | Top 1% (25902) | Top 2% (51803) | Top 5% (129508) | Top 10% (259016) |
|---|---|---|---|---|---|---|---|---|---|
| $\sigma_2$ | 0% (0) | 0% (0) | 0% (0) | 4% (3) | 21% (17) | 48% (39) | 76% (62) | 96% (79) | 100% (82) |
| Library Size ($\sigma_2$) | 0.88 M | 2.15 M | 6.69 M | 10.8M | 49.2 M | 97.2 M | 194.5 M | 615.7 M | 1150.6 M |
| VMAT2 | 0% (0) | 0% (0) | 2% (1) | 14% (7) | 24% (12) | 29% (15) | 65% (33) | 96% (49) | 100% (51) |
| Library Size (VMAT2) | 3.31 M | 4.63 M | 6.65 M | 16.0M | 39.5 M | 76.7 M | 171.2 M | 553.8 M | 1544.1 M |
| VAChT | 0% (0) | 2% (1) | 8% (5) | 14% (9) | 28% (18) | 47% (30) | 66% (42) | 95% (61) | 98% (63) |
| Library Size (VAChT) | 0.87 M | 2.10 M | 4.20 M | 10.6M | 26.7 M | 53.8 M | 119.2 M | 347.5 M | 963.3 M |

**Table S5. V-SYNTHES recall performance on CombiDOCK-identified actives.**

| | | |
|---|---|---|
| num_per_first_search | Number of Syn1 poses kept for each Syn1 hierarchy per orientation | 10 |
| num_per_second_search | Number of full poses kept for each Syn2 hierarchy coupled with each Syn1 pose | 5 |
| hdb_db2_search_score_threshold_first | The score cutoff for each segment of the branch in Syn1 | 100.0 |
| hdb_db2_search_score_threshold_second | The score cutoff for each segment of the branch in Syn2 | 100.0 |
| total_strain | Total strain energy threshold for each conformation | 8.0 |
| max_strain | Maximum strain energy threshold for each conformation | 1.5 |
| receptor_site_file | Specifies the sphere file | dock6files/matching_spheres.sph |
| max_orientations | The maximum number of orientations for the rigid segment to match the receptor spheres | 1000 |
| dock3.5_score_primary | Use the dock3.5 scoring as the primary scoring function | yes |
| dock3.5_grd_prefix | Path to files containing dock3.5 grid | dock6files/chem52 |
| dock3.5_vdw_score | Calculate vdw interactions | yes |
| dock3.5_electrostatic_score | Calculate electrostatic interactions | yes |
| dock3.5_ligand_desolvation_score | Calculate volume-based ligand desolvation from solvation grids | volume |
| dock3.5_solvent_occlusion_file | Path to solvent grids when desolvation score is turned on | dock6files/ligand.desolv.heavy |
| minimize_ligand | Whether minimize the docked ligand conformation or not | yes |
| simplex_max_iterations | Maximum number of iterations in simplex minimization | 1000 |
| score_threshold | Only poses with scores less than this value will be saved to the output mol2 file | -30.0 |

**Table S6. Parameters for CombiDOCK.** [a]These parameters were used in the ultra-large library docking campaign.

| | 2comp | | 3comp | |
|---|---|---|---|---|
| desired_charge | +3 | | | |
| wrong_charge_penalty | 50 | | | |
| desired_charge_no_penalty_limit | -3 | | | |
| hard_charge_diff_threshold | 6 | | | |
| num_expand_nodes | 200000 | 100000 | 160000 | 50000 |
| explore_weight | 0.001 | 0.002 | 0.001 | 0.002 |
| hardcode_scale_score | 40 | | | |
| default_error_value | 0 | | | |
| n_selection_1st_synthon | 500 | | | |
| n_selection_2nd_synthon | 50 | | 20 | |
| total_strain | 20 (default) | | | |
| max_strain | 10 (default) | | | |
| num_new_molecules_threshold | 15000 | | 6000 | |
| num_bad_cycles_threshold | 10 | | | |
| num_rollout | 1.5M | | 0.1M | |

**Table S7. Relevant parameters for MINT-Dock benchmarking on 46 targets.**

| | $\sigma_2$ | | VMAT2 | |
|---|---|---|---|---|
| | **2comp** | **3comp** | **2comp** | **3comp** |
| desired_charge | +1 | | | |
| wrong_charge_penalty | 50 | | | |
| desired_charge_no_penalty_limit | 0 | | | |
| hard_charge_diff_threshold | 1 | | | |
| num_expand_nodes | 200000 | 100000 | 160000 | 50000 |
| explore_weight | 0.001 | 0.002 | 0.001 | 0.002 |
| hardcode_scale_score | $40e^{-0.01}$ = 39.6019933499667 | | | |
| default_error_value | 0 | | | |
| n_selection_1st_synthon | 500 | | | |
| n_selection_2nd_synthon | 50 | 20 | 50 | 20 |
| total_strain | 20 (default) | | 8 | |
| max_strain | 10 (default) | | 1.5 | |
| num_new_molecules_threshold | 15000 | 6000 | 15000 | 6000 |
| num_bad_cycles_threshold | 10 | | | |
| num_rollout | 10M | 2M | 15M | 2M |

**Table S8. Relevant parameters for single-target MINT-Dock prospective screening campaigns.**

| | Prospective | | Backpropagation Comparison | |
|---|---|---|---|---|
| | **Selectivity** | **Polypharm** | **Selectivity** | **Polypharm** |
| | 2comp | 2comp | 3comp | 3comp |
| **desired_charge** | 1 | | 1 | |
| **wrong_charge_penalty** | 50 | | 50 | |
| **desired_charge_no_penalty_limit** | 0 | | 0 | |
| **hard_charge_diff_threshold** | 1 | | 1 | |
| **num_expand_nodes** | 160000 | | 200000 | 100000 |
| **explore_weight** | 0.0001 | 0.001 | 0.0001 | 0.0001 |
| **hardcode_scale_score** | $40e^{-0.01} = 39.6019933499667$ | | | |
| **default_error_value** | -0.1 | | | |
| **target_multiplier_ontarget** | 2 | 1 | 2 | 1 |
| **target_multiplier_offtarget** | 0 | 1 | 0 | 1 |
| **n_selection_1st_synthon** | 1000 | 500 | | |
| **n_selection_2nd_synthon** | 50 | | 20 | |
| **total_strain** | 20 (default) | | | |
| **max_strain** | 10 (default) | | | |
| **num_new_molecules_threshold** | 30000 | 15000 | 6000 | |
| **num_bad_cycles_threshold** | 10 | | | |
| **num_rollout** | 15M | 10M | 2M | 2M |

**Table S9. Relevant parameters for multi-target MINT-Dock prospective screening campaigns.**
